# Continuous and discrete decoding of overt speech with electroencephalography

**DOI:** 10.1101/2024.05.23.595510

**Authors:** Alexander Craik, Heather Dial, Jose Luis Contreras-Vidal

**Affiliations:** Department of Electrical and Computer Engineering, University of Houston; Department of Communication Sciences and Disorders, University of Houston; NSF IUCRC BRAIN, University of Houston

**Keywords:** Electroencephalography (EEG), deep learning, convolutional neural networks, recurrent neural networks, attention networks, speech decoding, Electromyography (EMG) removal

## Abstract

Neurological disorders affecting speech production adversely impact quality of life for over 7 million individuals in the US. Traditional speech interfaces like eye-tracking devices and P300 spellers are slow and unnatural for these patients. An alternative solution, speech Brain-Computer Interfaces (BCIs), directly decodes speech characteristics, offering a more natural communication mechanism. This research explores the feasibility of decoding speech features using non-invasive EEG. Nine neurologically intact participants were equipped with a 63-channel EEG system with additional sensors to eliminate eye artifacts. Participants read aloud sentences displayed on a screen selected for phonetic similarity to the English language. Deep learning models, including Convolutional Neural Networks and Recurrent Neural Networks with/without attention modules, were optimized with a focus on minimizing trainable parameters and utilizing small input window sizes. These models were employed for discrete and continuous speech decoding tasks, achieving above-chance participant-independent decoding performance for discrete classes and continuous characteristics of the produced audio signal. A frequency sub-band analysis highlighted the significance of certain frequency bands (delta, theta, and gamma) for decoding performance, and a perturbation analysis identified crucial channels. Assessed channel selection methods did not significantly improve performance, but they still outperformed chance levels, suggesting a distributed representation of speech information encoded in the EEG signals. Leave-One-Out training demonstrated the feasibility of utilizing common speech neural correlates, reducing data collection requirements from individual participants.

## 1. Introduction

In 2019, over 7 million individuals in the U.S. faced significant communication barriers due to neurological conditions like Amyotrophic Lateral Sclerosis (ALS) and stroke-induced aphasia, substantially affecting their quality of life [1]. Traditional communication aids, reliant on the energy-intensive selection of letters or words, offer a slow and unnatural interaction process. Brain-computer interface systems (BCIs) present a transformative prospect for these individuals, leveraging neural modalities to decode speech in a more intuitive manner. Despite the inherent complexities of speech processes and previous limitations of machine learning, recent advancements in deep learning have led to progress in speech synthesis from neural signals, especially when decoding invasive electrocorticography (ECoG) [2].

Decoding continuous overt speech from ECoG, focusing on audio signal characteristics rather than discrete phonemes or words [3], marks a pivotal shift towards a universal BCI tool. However, invasive methodologies involve inherent risks and limitations related to surgery and signal degradation over time [4]. Non-invasive Electroencephalography (EEG), a more accessible neural modality that reduces the need for surgery, is challenged by lower spatial resolution and substantial noise from electromyography (EMG), motion, and electrooculography (EOG) artifacts [5]. Addressing these limitations is crucial, necessitating advanced denoising techniques coupled with sophisticated deep learning models for decoding.

The investigation into language processing through EEG has historically been skewed towards comprehension rather than production, primarily due to the hurdle of EMG contamination [6]. This contamination not only risks erroneous interpretations by decoding algorithms but also potentially overlooks critical information in the gamma frequency band, which has been shown to contain information associated with overt speech [7, 2, 3, 8, 9]. Effective EMG removal, therefore, is paramount, as it may help to enhance the signal-to-noise ratio (SNR) and decoding efficacy, especially within the promising lower gamma band (30 – 70Hz) [10, 11] where there is significant overlap with the EMG frequency contamination [12, 13, 14, 15]. Despite challenges posed by lower SNR and artifact interference [16], recent strides in denoising techniques and deep learning-based decoding methodologies [17] have begun to mitigate these limitations, showcasing potential in phoneme [18] and word classification [11].

This research stands at the confluence of pioneering EMG decontamination and advanced decoding techniques, aiming to transform EEG-based speech BCIs into viable, universal tools for enhancing communication for those affected by neurological disorders. This research aims to refine and validate a de-noising framework for speech-related artifact removal from EEG, and to explore the feasibility of deep learning-based decoding of both continuous features and discrete classes of overt speech through EEG. This approach not only addresses the technical challenges presented by EMG contamination but also aligns with the overarching goal of enabling more natural and efficient communication for individuals with speech impairments [19].

## 2. Background

Electromyographic (EMG) contamination in electroencephalography (EEG) signal processing represents a significant hurdle, particularly within the realm of EEG-based speech decoding efforts. The challenge stems from the spectral overlap between EEG and EMG signals, where EMG artifacts can affect frequencies as low as 13 Hz, complicating efforts to cleanly separate neural signals pertinent to speech processing [12, 13, 14, 15]. Sophisticated signal processing methodologies are required, as merely filtering out EMG-associated frequency ranges risks the elimination of essential EEG data. Approaches to EMG removal are broadly categorized into filtering methods—including low pass filters and adaptive filters—and Blind Source Separation (BSS) techniques, such as Canonical Correlation Analysis (CCA) and Independent Component Analysis (ICA). ICA is considered a cornerstone in EEG signal processing despite its limitations due to the assumptions of source count restriction, statistical independence, and non-Gaussianity of signals, which are not always applicable in speech-related EEG signal processing [5, 20, 21, 22, 23, 24, 25].

In light of these challenges, alternative strategies such as Empirical Mode Decomposition (EMD) and its enhanced variant, Ensemble EMD (EEMD), have been proposed to facilitate more effective artifact removal by decomposing signals into intrinsic oscillatory components prior to source separation techniques. These methods show promise in reducing mode-mixing and preserving critical EEG information, essential for accurate speech decoding [26, 27].

Decoding overt speech from EEG signals is an emerging field with the potential to revolutionize communication aids for individuals with speech impairments. The endeavor is multi-faceted, encompassing the collection and analysis of neural data, the development of sophisticated decoding algorithms, and the overcoming of methodological and technological limitations observed in previous research. Reviews of existing studies highlight several recurring challenges, including limited phonetic diversity, the use of low-density EEG systems, and insufficient attention to comprehensive EMG cleaning in the preprocessing stages [28, 29, 30, 31, 32, 33, 34].

Noteworthy attempts at speech decoding from EEG include efforts to classify hand-crafted features from binary classes of phonemes and words, and studies focusing on repeating specific commands or sentences. These efforts often face limitations in phonetic diversity, underlining the necessity for datasets that capture a broad spectrum of speech nuances [28, 29, 30, 31, 32, 33]. For instance, studies employing high-density EEG systems in controlled environments have generated quality datasets but have been limited by their experimental scope and the depth of decoding attempts [30, 31].

Critically, none of the preliminary research endeavors undertook extensive EMG cleaning, a gap that this research aims to fill. The proposed methodology seeks to collect a comprehensive dataset using a high-density EEG system meticulously aligned with recorded audio signals. This dataset not only encompasses a wide range of phonetic diversity but also ensures a cleaner signal for decoding through advanced EMG artifact removal techniques, thereby addressing the dual challenges of signal purity and decoding accuracy. By integrating advanced EMG cleaning techniques with a robust decoding framework, the aim is to push the boundaries of what is possible in non-invasive speech BCI technology, offering new avenues for communication for those unable to speak.

## 3. Methods and materials

### 3.1. Data collection

This data was collected at the University of Houston under Study #3321, which was approved by the University of Houston Institutional Review board. Nine participants were recruited and fitted with a 63-channel Brain Products ActiChamp Plus EEG system [35]. Four passive electrooculography (EOG) sensors were placed around the eyes for use in the removal of eye artifacts. Two additional sensors were relocated from EEG locations O1 and O2 to positions around the mouth for paralabial EMG collection, specifically at the superior and inferior orbicularis oris. The location of the facial EMG sensors is based on previous research into EMG channels for speech recognition. In [36], the authors optimized EMG channel locations for an sEMG speech synthesis framework, finding the orbicularis oris as one of seven optimum channel selections for speech decoding. In [37], where the authors optimized sensor locations for the specific application of removing EMG artifacts for an overt speech task, the authors found that the coherence between the EMG sensors and the speech envelope was highest at the orbicularis oris muscles. A Brain Products StimTrak was used for audio collection, which, with a common trigger box, allows for effective synchronization between the EEG and audio signals. Each participant was seated comfortably in a WhisperRoom [38], a room designed to minimize the presence of background noise in the audio signal. Synchronized EEG, EMG, EOG, and audio were collected with a sampling frequency of 25,000 Hz.

### 3.2. Experimental protocol

The protocol involved periods of rest followed by the audible repetition of a sentence presented on a display. This protocol is presented with Figure 1. In this protocol, each block begins with a minute of rest before cycling through 52 selected sentences of overt speech. Following each produced sentence, the participant rests for five seconds. Each participant successfully completed between 3–4 blocks per session. Five of nine participants underwent two collection sessions (eight total blocks), with four opting for a single session due to scheduling constraints. The graphical interface was developed with Matlab toolbox Psychtoolbox [39].

**Figure 1:**
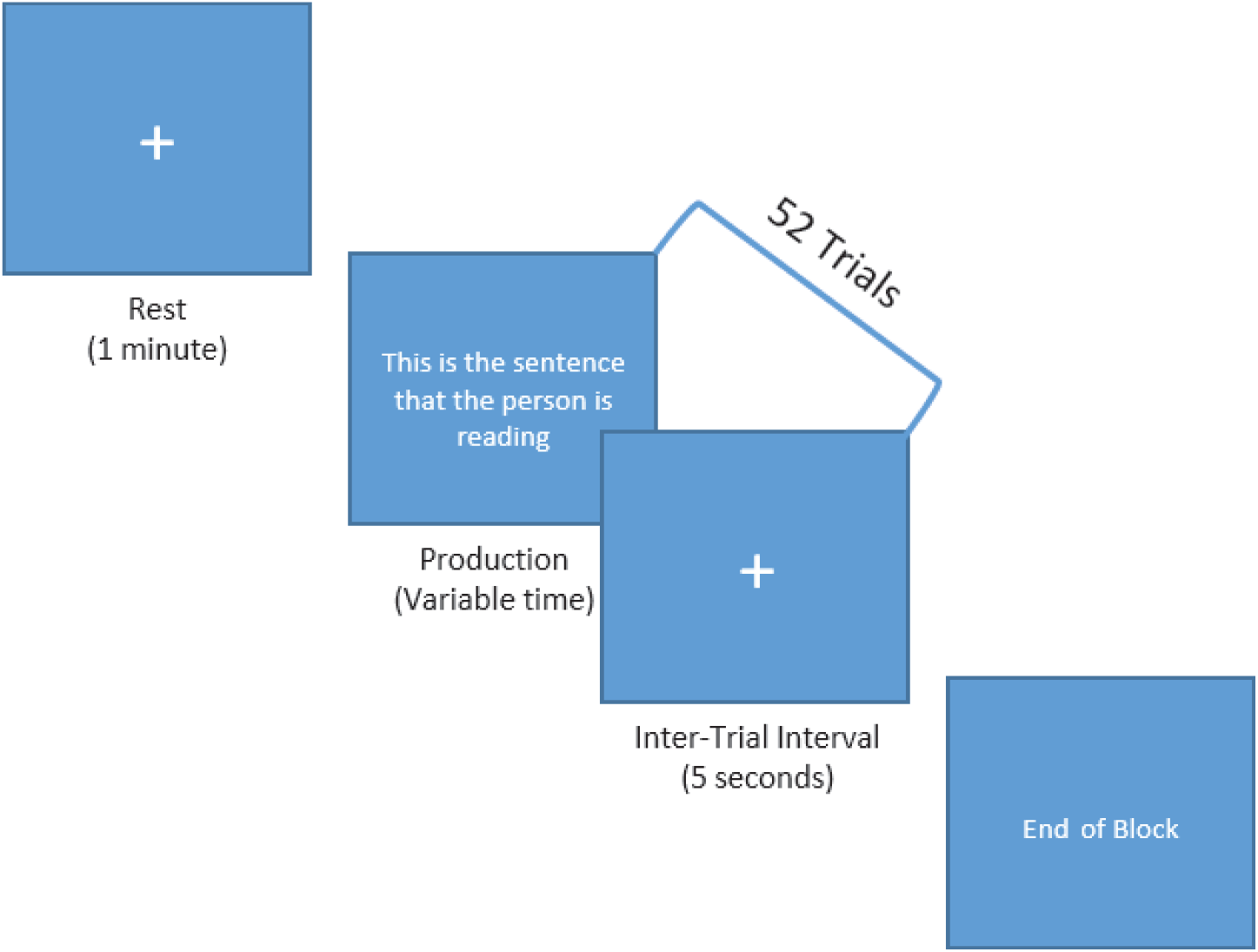
The block design for the overt speech experimental protocol.

In [3], the sentences were selected from the MOCHA-TIMIT database [40], which is a collection of 460 sentences with a phoneme representation that closely matches that of the English language. Instead, to increase the chances of success for discrete prediction, three well-researched passages were selected for repetition: the Rainbow passage, the Caterpillar passage, and the Grandfather passage. The grandfather passage [41] is a public domain text frequently used to gather speech samples, as it contains nearly all English phonemes. In this study, the extended Grandfather passage was used as the original formulation did not include the /UH/ phoneme, and phonemic coverage is desirable for the end application of full speech spectrum synthesis. The Rainbow passage [41] was similarly designed to include the variety of sounds and mouth movements used in the English language. The Caterpillar passage was also designed for phonemic similarities to the English language, with this passage leading the other two in phonemic balance [41]. To avoid auditory event related potentials, no audible triggering was employed. Instead, all protocol prompts are displayed visually. All sentences were displayed in pseudo-random order to prevent any anticipatory responses from the participant. Following collection, a Hidden Markov Model toolkit (Montreal Forced Aligner [42]) was used to automatically find the time points that correspond to the onsets of individual phoneme production (following the ARPAbet phoneme naming convention [43]). Transcripts for all participants were manually corrected at the word level prior to processing with the forced aligner. The resulting phoneme counts following 1, 4, and 8 blocks of collection are presented with Tables 5 and 6 in the Section 9.1 of the Appendix.

### 3.3. Signal processing prior to EMG removal

In preparation for EMG removal analysis, EEG signals underwent a preprocessing pipeline, adapted from Makoto’s recommended processing pipeline [44]. Initial steps included resampling signals to 200 Hz and applying both low and high-pass filters to accommodate a broad spectrum of analysis, up to the gamma frequency band (30 – 100 Hz). Despite initial efforts to remove power line noise using EEGLab’s Zapline function [45], persistent contamination necessitated the implementation of a notch filter specifically for the EOG sensors (Section 9.4 of the Appendix).

A major contamination source in EEG experiments is related to eye movement, and denoising of eye movement artifacts was accomplished with an adaptive noise cancellation (ANC) framework, specifically the H-infinity sub-optimal ANC, which employs EOG sensor data to estimate and remove noise without eliminating underlying EEG information [46]. Optimal parameterization of this ANC approach was derived from extensive prior research, setting parameters *γ*, *p0*, and *q0* based on their efficacy at a 200 Hz sampling frequency, to best discriminate between eye movement artifacts and neural activity [47].

While the resampling function includes a low pass filter, a secondary low pass filter at 100 Hz was then applied to mitigate any potential high-frequency artifacts introduced during the initial processing steps. To counteract the risk of distributing channel-specific noise across the EEG dataset, Artifact Subspace Reconstruction (ASR) was utilized to identify and correct for aberrant signals, ensuring that the preprocessed EEG was primed for the most effective EMG artifact removal possible [48]. Following ASR, Common Average Reference (CAR) method was applied for signal normalization. This pipeline is represented in Figure 2.

**Figure 2:**
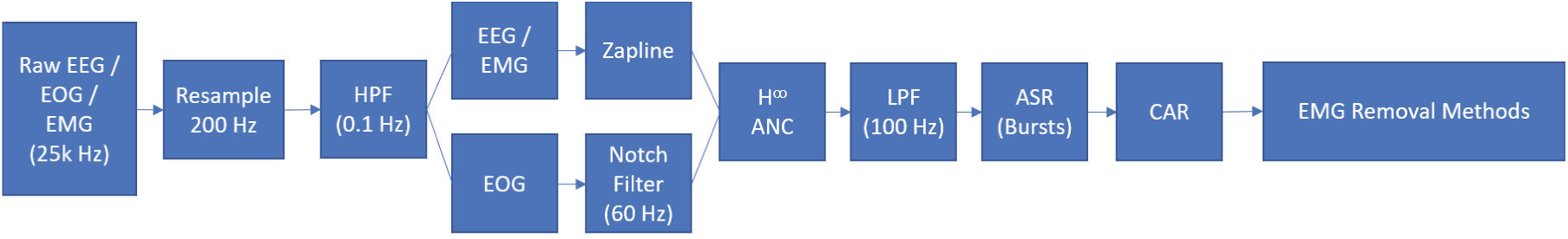
The preprocessing pipeline employed prior to the EMG removal method analysis.

### 3.4. EMG removal methods

The process of EMG artifact removal from EEG data is crucial for assessing artifactual influence on the accuracy of speech decoding. The process for this research involved extensive parameter selection and methodological implementation across various EMG cleaning methods. The implementation details were based on adaptations from EEGLab for ICA and CCA methods, and from the ReMAE toolbox [49] for EEMD, an extensive toolbox designed to address the challenges of EMG artifact interference. The methods and threshold types are presented with Figure 3.

**Figure 3:**
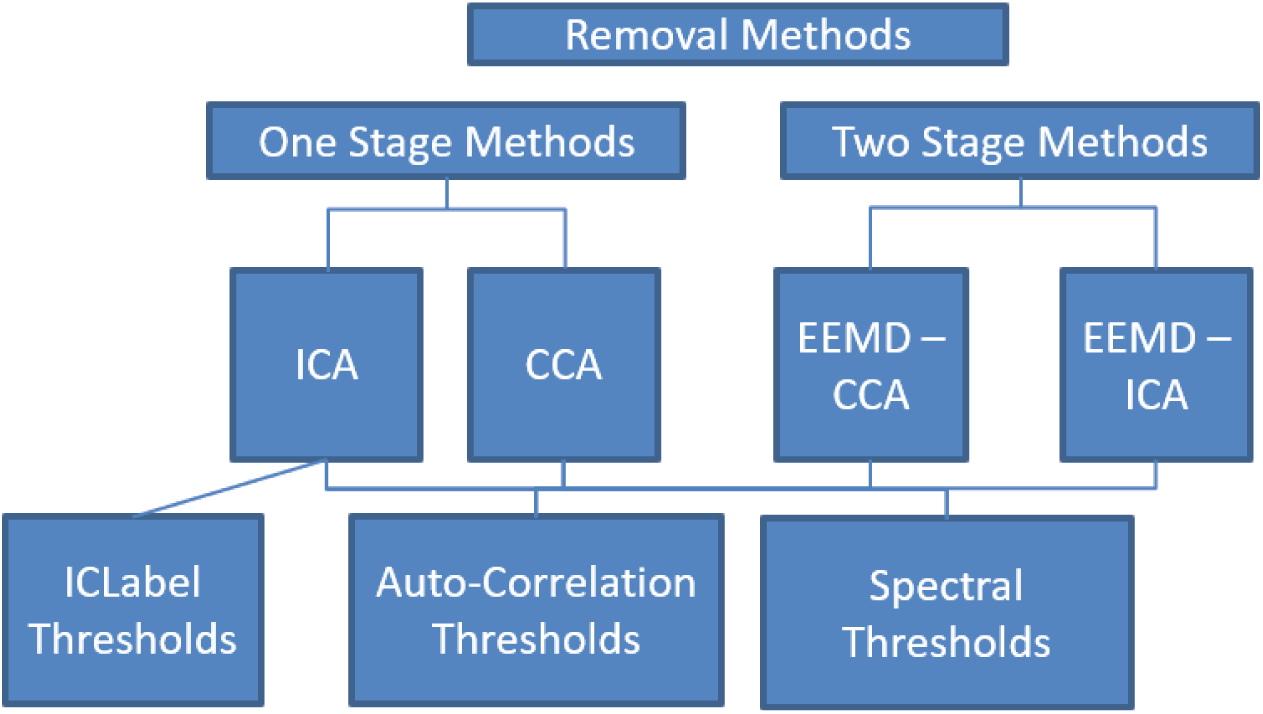
EMG removal methods and artifactual component threshold rejection types.

Single-Stage Blind Source Separation (BSS) techniques (Section 9.5.1 of the Appendix), including ICA and CCA, were specifically chosen for their reported efficiency and accuracy [50, 51]. ICA implementations utilized the SOBI variant for its superior computational speed and accuracy over other ICA forms like JADE, Infomax, and FastICA [52]. To reduce computational demands for CCA implementations, this research employed a one-point time lag.

EEMD Parameter Selections (Section 9.5.2 of the Appendix) involved defining the noise level, the number of ensembles, and the number of Intrinsic Mode Functions (IMFs) to effectively decompose each channel. A noise level of 0.2 times the standard deviation of input data and 10 ensembles were chosen based on prior research findings, with six IMFs identified as an optimal count for effective EEG signal discrimination [53]. A crucial step following EEMD decomposition was the selection of a subset of IMFs for further BSS analysis, with an autocorrelation value of 0.95 serving as the threshold for isolating potentially artifactual components. As described in [54], the threshold selection for the first stage in the EEMD-BSS process is not as important, as this simply collects potentially artifactual components, and does not directly remove data.

An Artifactual Component Assessment (Section 9.5.3 of the Appendix) was conducted through the evaluation of EEG components to identify and remove EMG artifacts. This assessment utilized both spectral content and autocorrelation thresholds, applying these to components derived from all EMG removal methods (ICA, CCA, EEMD-ICA, EEMD-CCA). The spectral thresholds, based on the power within specific frequency ranges [55], and autocorrelation thresholds [54, 56] were set to distinguish EEG from EMG data, enabling a comprehensive and comparative analysis across different EMG removal methods.

To ensure the robustness and reliability of the analysis, Artifactual Component Rejection Threshold Selection (Section 9.5.4 of the Appendix) employed Kmeans clustering [57] to determine threshold levels that would yield distinctly cleaned datasets. Relying on threshold intervals designed from past research resulted in datasets that were very similar, if not identical. Instead, this approach facilitated a nuanced differentiation of artifacts, accounting for variability across participants and sessions.

The Effectiveness of EMG Cleaning Methods (Section 9.5.5 of the Appendix) was evaluated across various metrics, including signal RMSE, spectral analysis, and the correlation of metrics to overt speech decoding performance. This involved comparing cleaned EEG data against pre-processed signals to assess the impact of EMG removal on signal quality and speech decoding accuracy.

For detailed implementation details, parameter selections, and a thorough evaluation of EMG cleaning methods’ effectiveness, refer to the Appendix. This comprehensive approach underscores the nuanced and methodical efforts undertaken to refine EEG data for enhanced speech decoding, illustrating the intricate balance between artifact removal and data integrity preservation.

### 3.5. Selection of EMG removal methods for decoding analysis

The comprehensive assessment of EMG cleaning methods within EEG data analysis for speech decoding tasks emphasized nuanced threshold levels and detailed metric evaluations, crucial for optimizing the removal process. Through a Kmeans clustering approach, participant-averaged thresholds demonstrated low variance, indicating consistent component characteristics across individuals. These thresholds spanned across various cleaning methods, including ICA, CCA, and EEMD, showing alignment with previous research benchmarks and suggesting tailored adjustments for artifact rejection based on either spectral content or autocorrelation criteria.

Signal RMSE metrics (Section 9.5.6 of the Appendix) revealed that two-stage methods generally induced less signal distortion compared to one-stage methods, particularly highlighting the precision of correlation-based rejection methods in targeting speech-related artifacts. However, ICLabel showcased the least signal distortion, suggesting a less aggressive cleaning approach may sometimes be advantageous. The correlation analysis underscored a relationship between cleaning methods and speech decoding performance, advocating for a balanced approach in artifact removal to retain essential EEG data quality.

A Spectral RMSE and Relative Spectral Change analysis (Section 9.5.7 of the Appendix) further delineated the effectiveness of two-stage methods, especially in low-frequency bands where lesser distortion was noted, potentially reflecting more effective preservation of EEG data. Notably, the choice of component rejection method and threshold levels significantly influenced the outcome, with autocorrelation rejection showing efficacy in removing higher-frequency gamma activity without excessively distorting essential EEG information.

A novel aspect of the evaluation involved the reconstructed artifact analysis and source analysis (Section 9.5.8 of the Appendix), providing insights into the spatial effectiveness of EMG removal methods and their capacity to preserve critical speech-related EEG components. Specifically, two-stage methods and certain threshold levels were identified as more conducive to maintaining essential data within speech production regions of the brain, such as Broca’s and Wernicke’s areas.

The Decoding Performance Analysis (Section 9.5.9 of the Appendix) directly correlated these methodological nuances with the practical outcome of speech decoding accuracy based on relatively simple CNN and RNN deep learning architectures. Findings pointed towards the selection of specific datasets and cleaning methods, such as EEMD-ICA and EEMD-CCA with medium or low thresholds, which showed promise in enhancing decoding performance by effectively balancing artifact removal with the preservation of vital EEG data.

Based on these findings, it is suggested that the datasets with the highest likelihood of more completely removing EMG contamination were based on either EEMD-ICA or EEMD-CCA. Datasets that have the greatest likelihood of obtaining high performance for speech-related decoding tasks were EEMD-CCA with medium thresholds, EEMD-ICA with low PSD thresholds, ICLabel with high thresholds (low cleaning), and the pre-processed signal itself. These four datasets were employed for the following more extensive decoding analysis.

### 3.6. Discrete class formulation

While there is support for the feasibility of continuous prediction of acoustic characteristics in invasive recordings [3], the large size of the collected datasets, the number of participants, and the resulting long training times made direct optimization of a model for continuous acoustic prediction infeasible. Instead, models were trained and optimized for the decoding of discrete phonetic and articulatory classes. This empirically-supported assumption was that the resulting models would be optimized for speech feature extraction and that the feature extraction architecture design could be used directly for acoustic characteristic prediction with minor adjustments to the model, specifically with the dense and output layers.

Given this discrete-to-continuous architecture optimization mechanism, the stimuli were selected for this proposed protocol to give a full representation of the speech spectrum, rather than equal counts of phonemes, as this made any continuous decoding model more applicable to real-world scenarios, where phoneme representation matches that of the English language. In the experimental protocol, each block was composed of a single repetition each for the Rainbow passage, Caterpillar passage, and extended Grandfather passage. The audio data was then segmented into phonemes using the Montreal Forced Aligner (MFA) [42], a tool that aligns speech recordings with their transcriptions at the phoneme level. MFA employs forced alignment to match phonetic segments in the speech signal to their corresponding phoneme labels in the text, providing precise synchronization and temporal information about phoneme occurrence and duration. The resulting collection of phonemes is separated into different articulatory and phonetic classes. In the current study, the analysis focused on the presence of voice, the place of articulation for consonants, the place of articulation for vowels, the manner of articulation for consonants, and whether a phoneme is classified as a vowel or a consonant. In addition to articulatory features of the phoneme, direct classification of individual phonemes was also performed.

#### 3.6.1. Presence of voice

The presence of voice is a binary feature indicating whether the vocal folds vibrate during production of a phoneme [58]. There are two ways in which this class formulation may be considered: with or without vowels. The presence of voice is not a discriminant feature for vowels generally, as the articulation of all vowels includes the vibration of the vocal folds, which may affect the performance of a classifier depending on the inclusion of vowels into the class as a whole. For this reason, this research only assessed decoding performance on the presence of voice for consonants.

#### 3.6.2. Place of consonant articulation

The place of consonant articulation refers to where the airflow is restricted in the articulatory system to produce consonant sounds [59]. Different regions within the vocal tract are involved in creating these restrictions for various consonants. Labial consonants involve the constriction of airflow when the two lips come together (bilabial) or when the bottom lip presses against the upper row of teeth (labiodental). Examples include [p], [b], [m], [v], and [f]. Dental consonants are produced by pressing the tongue against the teeth, as in the case of [th] and [dh]. Alveolar consonants are formed by pressing the tip of the tongue against the alveolar ridge, which is the area of the roof of the mouth just behind the teeth. Examples include various sounds such as [t], [d], [n], [s], [z], [l], and [r]. Palatal consonants are created by pressing the tongue against the back of the hard palate, which is the region behind the alveolar ridge. Consonants such as [sh], [ch], [zh], [jh], and [y] fall into this category. Velar consonants, like [k], [g], and [ng], are produced by pressing the back of the tongue against the soft part at the back of the roof of the mouth.

#### 3.6.3. Manner of consonant articulation

While the place of articulation indicates where the articulation occurs within the vocal tract, the manner of consonant articulation refers to how a consonant sound is produced [59]. There are six main categories of consonant manners: stops, fricatives, affricates, approximants, and nasals. Stops are consonants where the airflow is completely blocked for a brief moment. They can be further classified into voiced stops, such as [b], [d], [g], and unvoiced stops, such as [p], [t], [k]. Fricatives are consonants in which the airflow is constricted, causing friction. They can be divided into different subcategories based on the place of articulation.

Labiodental fricatives include [f] and [v]. Dental fricatives include [th] and [dh]. Palate-alveolar fricatives include sounds like [sh] and [zh]. Affricates are consonants that begin with a stop and transition into a fricative sound. Examples include [ch] and [jh]. Approximants are consonants where the articulators are close together, but the airflow is not constricted enough to create significant turbulence. Examples include [y], [w], [r], and [l]. Nasals are consonants in which the airflow is directed through the nasal cavity. Examples include [m], [n], and [ng].

#### 3.6.4. Place of vowel articulation

Vowels can be classified based on the position of the articulators during their production [60]. Front vowels are produced when the tongue is positioned towards the front of the mouth. Examples of front vowels include [iy], [ih], [ey], [eh], and [ae]. Center vowels are articulated with the tongue positioned in the central area of the mouth. Examples of center vowels include [ah] and [er]. Back vowels are produced with the tongue positioned towards the back of the mouth. Examples of back vowels include [ao], [ow], [uw], and [uh].

### 3.7. Model architectures

To reduce the search space for neural network model optimization, constraints imposed in this study include window size and articulatory class selection. The input data was fixed at a 0.3-second window size, corresponding to 60 time points at a 200 Hz sampling rate, based on event-related potentials’ timings relative to phoneme onset, as seen in prior work [61, 3]. Optimization focused on the Place of Articulation for Vowels due to its lower sample size, facilitating faster training and a balanced distribution [41, 60]. The authors note that initial empirical testing provided support that performance changes due to architecture differences matched well across classes, and only computational constraints prevented optimization across all classes.

Convolutional Neural Networks (CNNs), renowned for processing grid-like data [62], employ convolutional layers with filters to detect data patterns, using hierarchical layers and downsampling through pooling to reduce dimensionality and computational demand. Strategies such as Batch Normalization [63] and dropout layers [64] combat overfitting by normalizing activations and randomly deactivating neurons, respectively. CNN optimization explored architecture aspects like the number of layers, filters per layer, filter size, residual connections, and final feature vector size, aiming to balance feature detection capability with manageable parameter counts and computational efficiency. The incorporation of separable convolutional layers and spatial depth-wise convolutional layers as employed in EEGNet [65] alongside initial temporal pooling significantly reduces trainable parameters, vital for EEG data limitations.

Recurrent Neural Networks (RNNs) excel in sequential data processing, utilizing an internal state to capture temporal dependencies, essential for language modeling and speech recognition [66]. Similar to CNNs, RNN optimization included dropout and layer normalization to mitigate overfitting, focusing on the number of layers, types and number of units, bidirectionality, residual connections, and final feature vector sizing. RNN types assessed include Long Short-Term Memory (LSTM) [67] and Gated Recurrent Unit (GRU) [68] cells, with attention to trainable parameter counts versus temporal dependency capturing capacity.

Attention modules, integrated with CNN or RNN architectures, enhance model performance by allowing dynamic focus on relevant input data parts, employing multiple attention heads to capture diverse information types [69]. This approach’s innovation lies in assessing whether attention modules can improve EEG-based speech synthesis models’ performance without overly complicating or extending training times. The optimization explored the module’s configuration, including the type of feature extraction, number of encoder blocks, attention heads, head sizes, and dense decoding units, assessing the balance between attention detail and computational demands.

### 3.8. Discrete classification with the optimized models

The study advances speech BCI research by employing optimized CNNs, RNNs, and attention models for discrete classification of articulatory kinematic classes and individual phonemes. This approach investigated the distinguishability of phonemes and phoneme characteristics, potentially enhancing discrete speech BCIs. Key methodologies include isolating frequency bands to identify those critical for overt speech decoding and assessing EMG removal methods’ effectiveness. In this study, the following frequency ranges are employed for Delta (0.1-4 Hz), Theta (4-8 Hz), Alpha (8-12 Hz), Beta (12-31 Hz), Lower Gamma (31-70), and Upper Gamma (70-100 Hz). This spectral assessment is presented with Figure 4.

**Figure 4:**
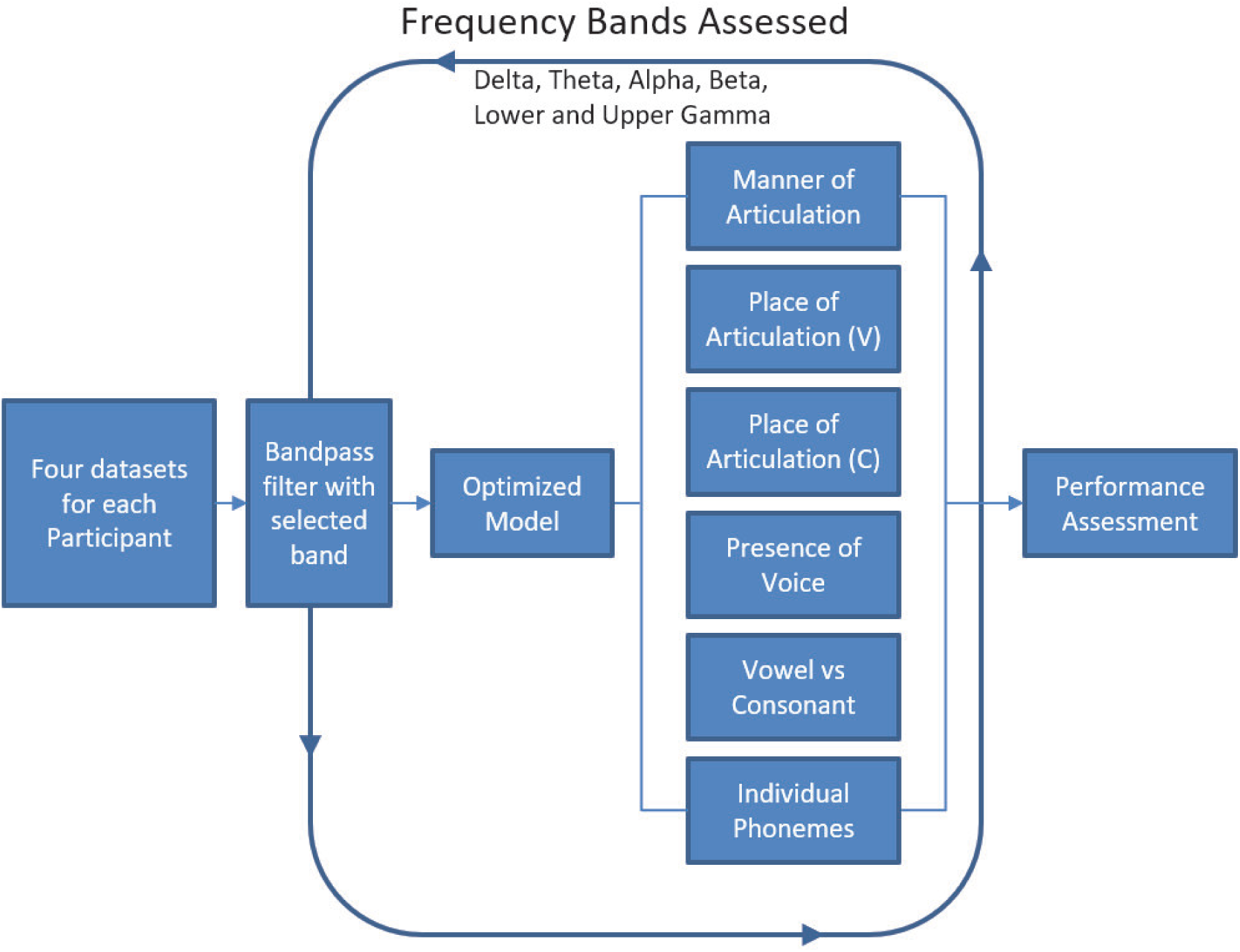
Frequency band assessment. V = vowel; C = consonant.

To further examine the impact of EMG removal, a perturbation analysis was conducted, assessing performance changes when individual EEG channels are zeroed out. This method evaluates EMG removal efficacy based on performance dips upon the removal of channels near potential noise sources, such as EEG electrode locations at FP1 and FP2, indicating less effective EMG removal for channels with higher EMG contamination. This perturbation analysis is presented with Figure 5.

**Figure 5:**
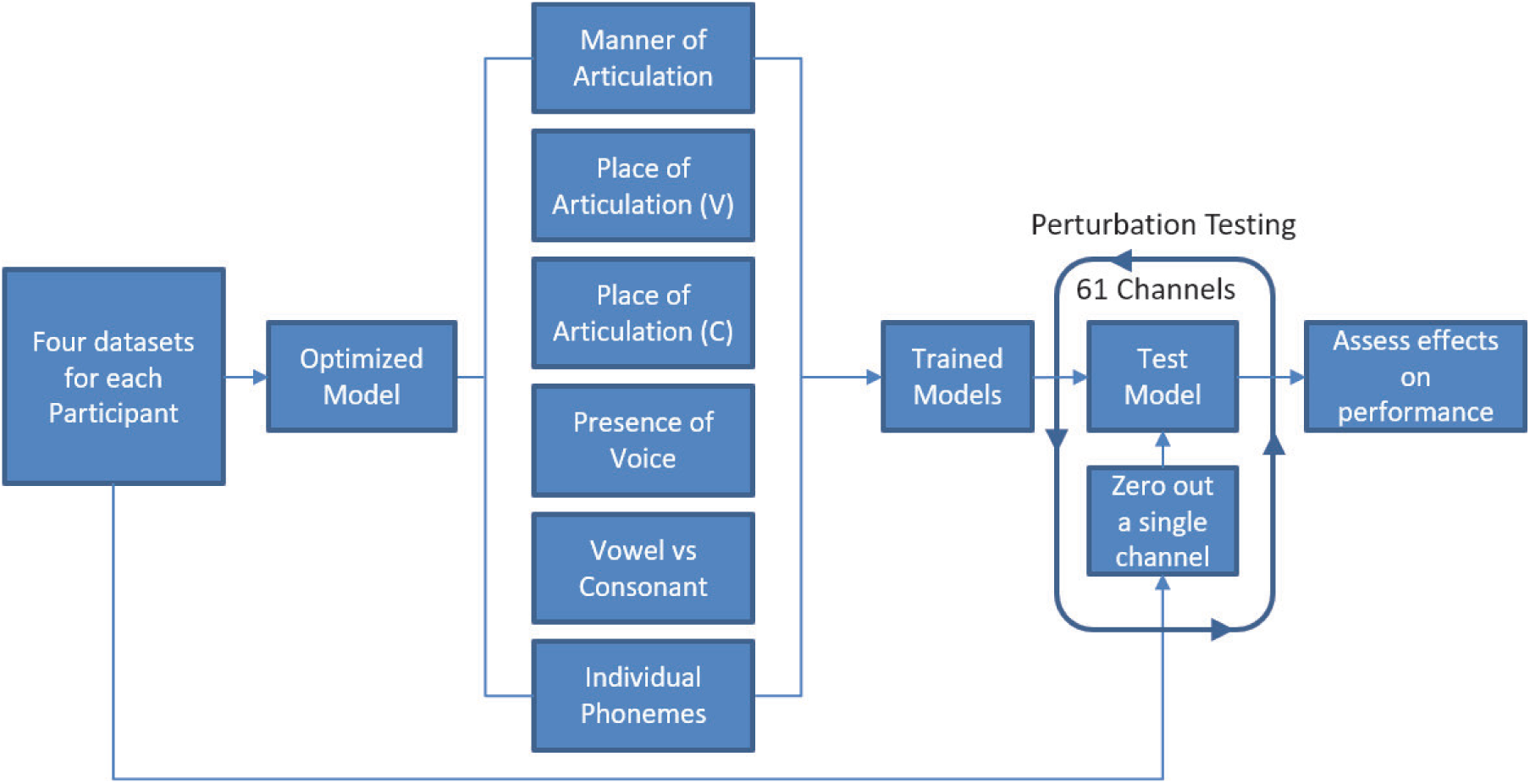
Perturbation analysis. V = vowel; C = consonant.

Channel selection strategies were explored to address challenges in decoding due to the large number of EEG channels. Reducing the number of necessary EEG electrodes is an important goal for both clinical and commercial applications, as the reduced electrode count would help to reduce the amount of effort needed for system setup, as well as minimize the size and cost of the EEG system generally. To assess different selection strategies, manual selection based on literature and an automated method using Common Spatial Pattern (CSP) analysis were compared. The manual approach selected channels close to areas known for speech processing, including the Post-central gyrus, Wernicke’s area, Pre-motor cortex, Broca’s area, and auditory cortices [70]. CSP analysis, implemented via the MNE Python toolkit [71], identified channels contributing most to discriminative spatial patterns, optimizing the decoding task’s channel set for each dataset and class. The top 11 channels were selected based on CSP filter values, ensuring comparability with manually selected channels. This channel selection analysis is presented with Figure 6.

**Figure 6:**
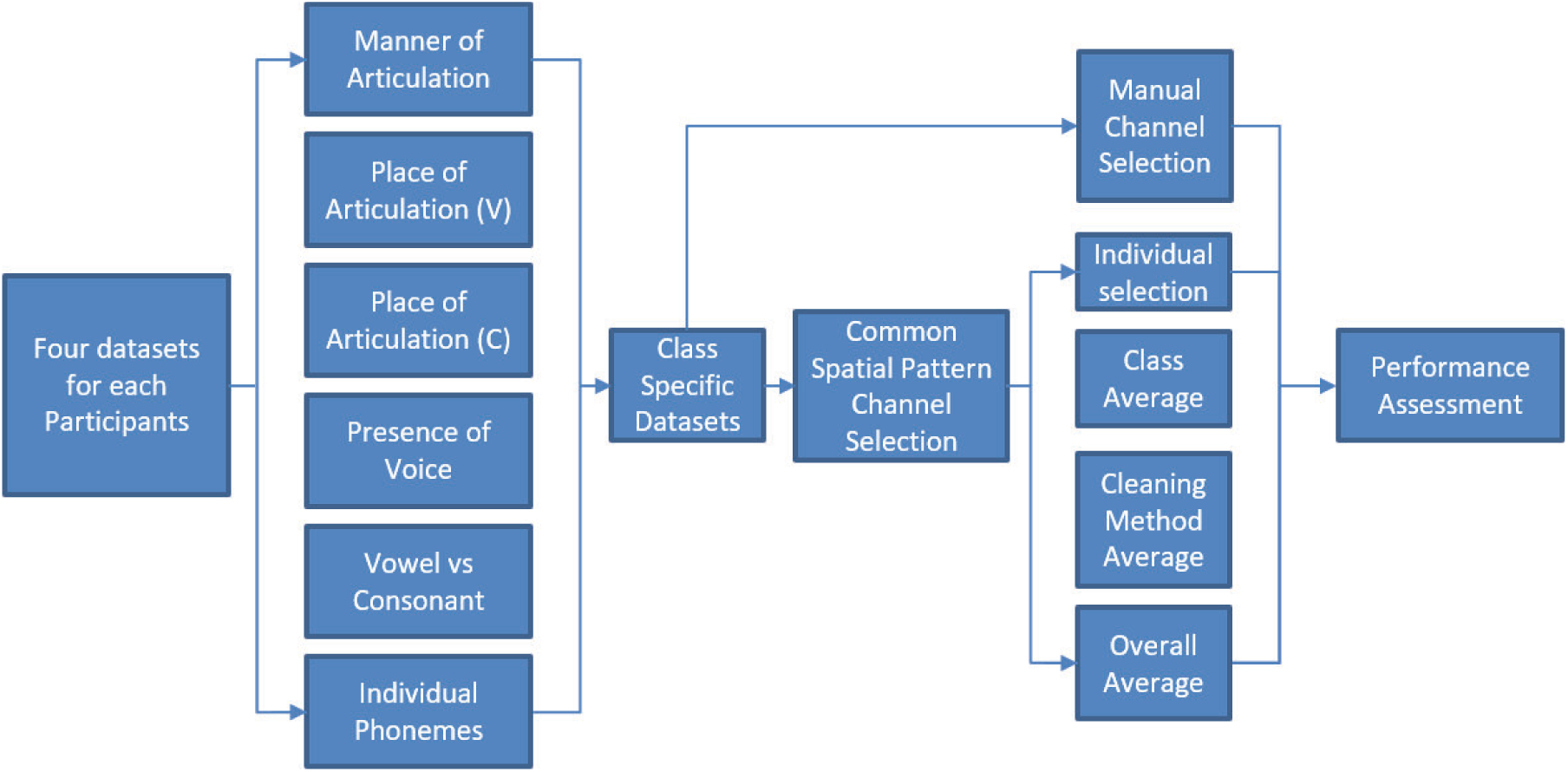
Channel selection analysis. V = vowel; C = consonant.

Leave-One-Out (LOO) testing assessed whether data from other participants could improve decoding for a specific participant. This involved classifying a participant’s data using models trained on data from others, with variations in the inclusion of the participant’s data in training or validation datasets. This method examined cross-participant generalizability and the impact of individual data on model accuracy, comparing results to participant-independent decoding accuracies. The LOO analysis is presented with Figure 7.

**Figure 7:**
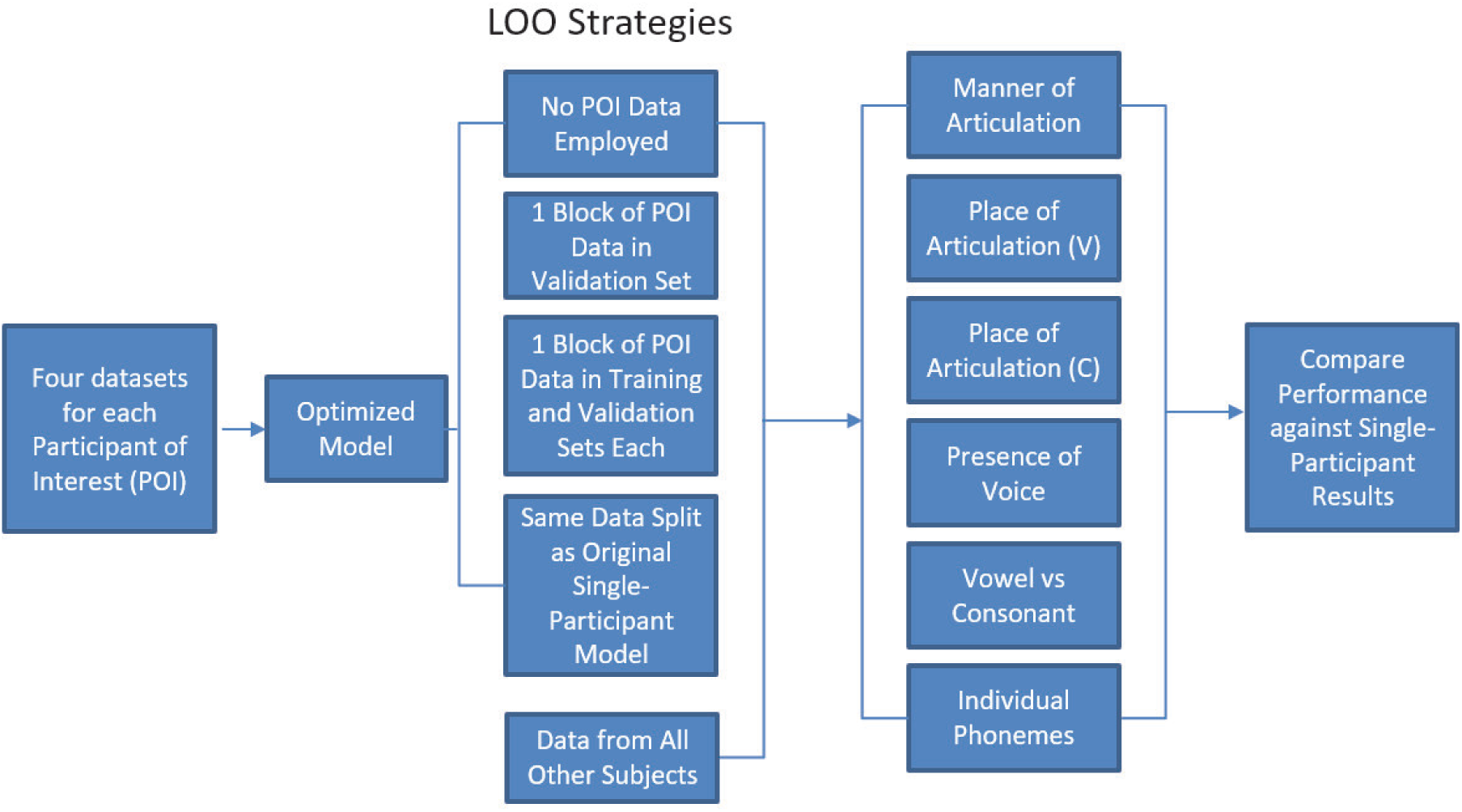
Leave-One-Out training assessment. V = vowel; C = consonant.

### 3.9. Continuous predictions with the optimized models

The continuous decoding of overt speech in this research focused on the prediction of Mel-Frequency Cepstral Coefficients (MFCCs), which are coefficients of a modified power spectral density curve of the acoustic signal [72]. The MFC is a modified power spectral density curve where the frequency bins are not evenly spaced. Instead, the frequency bins are designed to more closely match the perceptual ranges of the human auditory system. 25 MFCC’s were extracted from the audio signal at 200 Hz and synchronized with the EEG data collected through the protocol. 0.3 seconds of EEG data were extracted and centered around every time point of the MFCCs for prediction. Performance comparisons were made by calculating the Mel-Cepstral Distortion (MCD) [73] between the predicted and true MFCCs (lower MCDs imply more effective speech synthesis). The equation for Mel-Cepstral Distortion is,

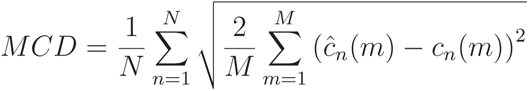

where N is the total number of time steps, M is the number of cepstral coefficients (in this case is 25), ĉn(m) represents the MFCs of the synthesized or predicted speech for the n-th frame and m-th coefficient, and cn(m) represents the MFCs of the original for the n-th frame and m-th coefficient.

The four selected EEG datasets per person used in the model optimization framework were again employed here to allow for a better understanding of how the EMG cleaning method affected continuous prediction performance. The three models optimized for discrete classification were also used here, with the sole difference being the final prediction layer. To handle the different output formats, for continuous predictions, the final layer employed a linear activation to produce continuous variable predictions, rather than the softmax activation employed for discrete classification, which outputs a single integer value representing the decoded class. The general continuous feature decoding pipeline is presented with Figure 8.

**Figure 8:**
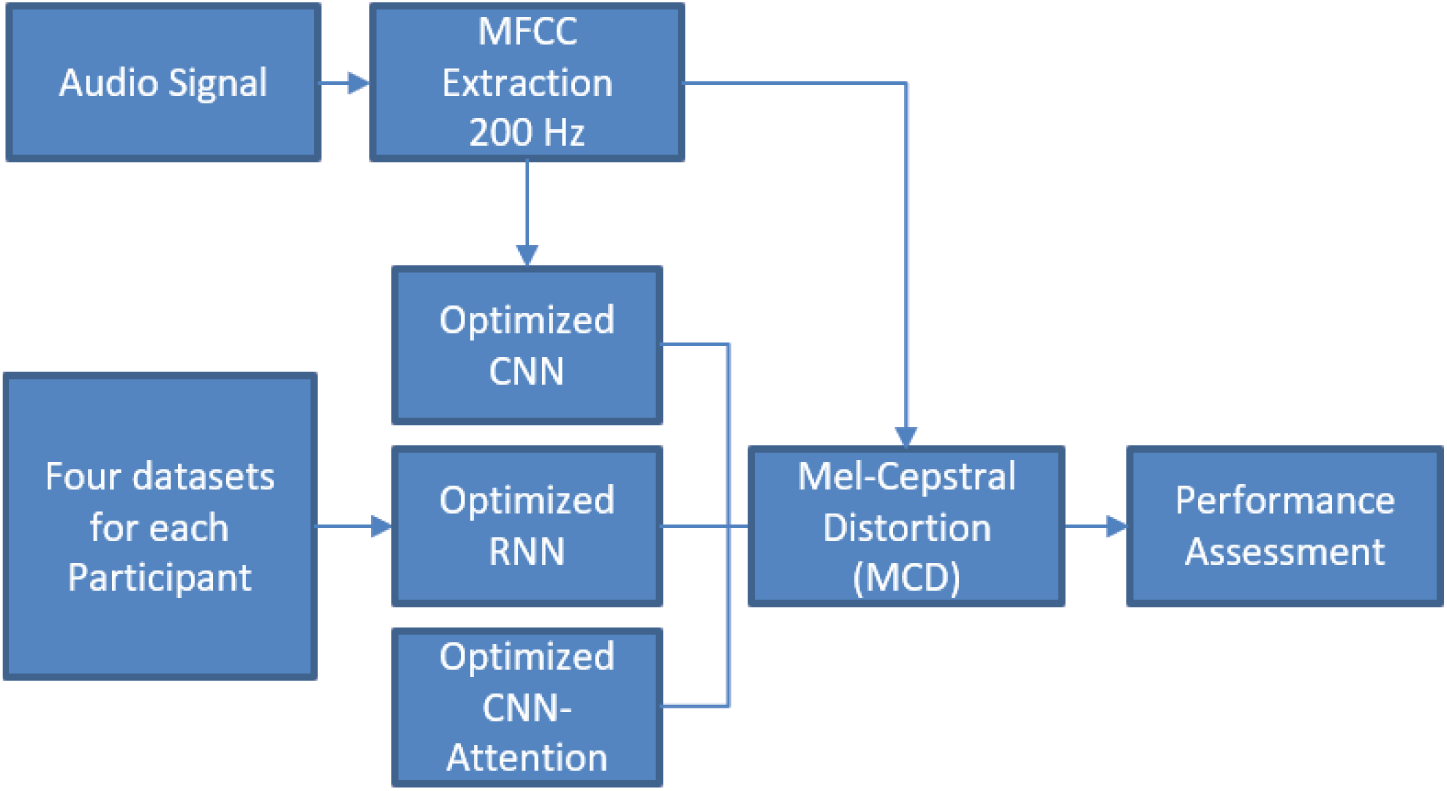
Continuous feature decoding pipeline. MFCC = Mel-frequency cepstral coefficient. CNN = convolutional neural network. RNN = recurrent neural network.

## 4. Results

### 4.1. Optimized architectures

This section details the results from the model optimization framework. All models were created with the Python deep learning library Tensorflow [74]. Model performances were compared against a chance model. Calculating a chance level based simply on the number of classes is not appropriate and would significantly inflate the performances of the trained models. Instead, the chance model was designed with Sklearn’s Dummy Classifier [75] with a stratified strategy. The stratified strategy forces the Dummy Classifier to learn the class distribution from the input data. During testing, the model randomly selects from this distribution. This was an appropriate selection for calculating chance level, as neither the phonemes nor the classes are equally balanced.

#### 4.1.1. Convolutional neural network

Table 1 presents the results from the optimization framework for the CNN architecture. Average performances across all participants are presented as performance above the chance model (i.e., model performance - chance model performance).

**Table 1:**
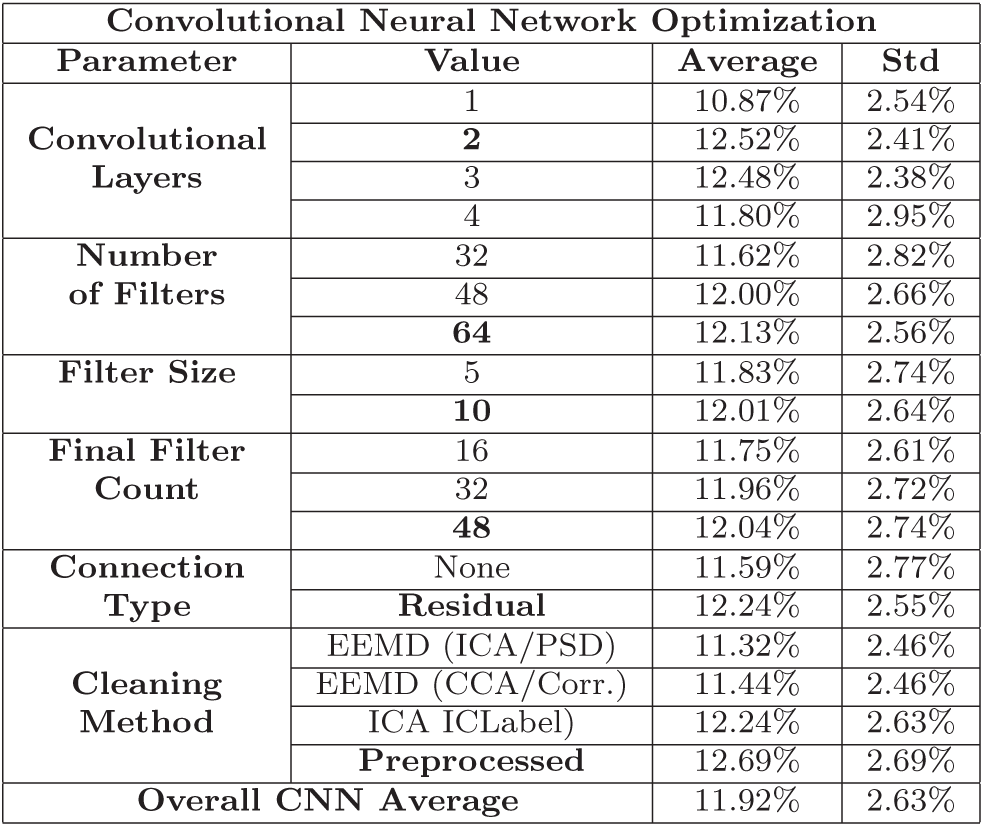
Optimization of the convolutional neural network models. Performance is presented as accuracy above the chance model.

| Convolutional Neural Network Optimization |  |  |  |
| --- | --- | --- | --- |
| Parameter | Value | Average | Std |
| Convolutional Layers | 1 | 10.87% | 2.54% |
|  | **2** | 12.52% | 2.41% |
|  | 3 | 12.48% | 2.38% |
|  | 4 | 11.80% | 2.95% |
| Number of Filters | 32 | 11.62% | 2.82% |
|  | 48 | 12.00% | 2.66% |
|  | **64** | 12.13% | 2.56% |
| Filter Size | 5 | 11.83% | 2.74% |
|  | **10** | 12.01% | 2.64% |
| Final Filter Count | 16 | 11.75% | 2.61% |
|  | 32 | 11.96% | 2.72% |
|  | **48** | 12.04% | 2.74% |
| Connection Type | None | 11.59% | 2.77% |
|  | **Residual** | 12.24% | 2.55% |
| Cleaning Method | EEMD (ICA/PSD) | 11.32% | 2.46% |
|  | EEMD (CCA/Corr.) | 11.44% | 2.46% |
|  | ICA ICLabel | 12.24% | 2.63% |
|  | **Preprocessed** | 12.69% | 2.69% |
| Overall CNN Average |  | 11.92% | 2.63% |

The optimization framework resulted in a CNN with three convolutional blocks, 64 filters, a filter size of 10, and a final filter count of 48. The network included residual connections between the second and third convolutional blocks. The cleaning method that results in the highest model performance is the ICA method with ICLabel EMG removal, but all cleaning methods underperformed against the solely pre-processed data, potentially indicating that the still-present EMG contamination in the solely pre-processed data helped decoding performance. The final CNN architecture is presented with Figure 9.

**Figure 9:**
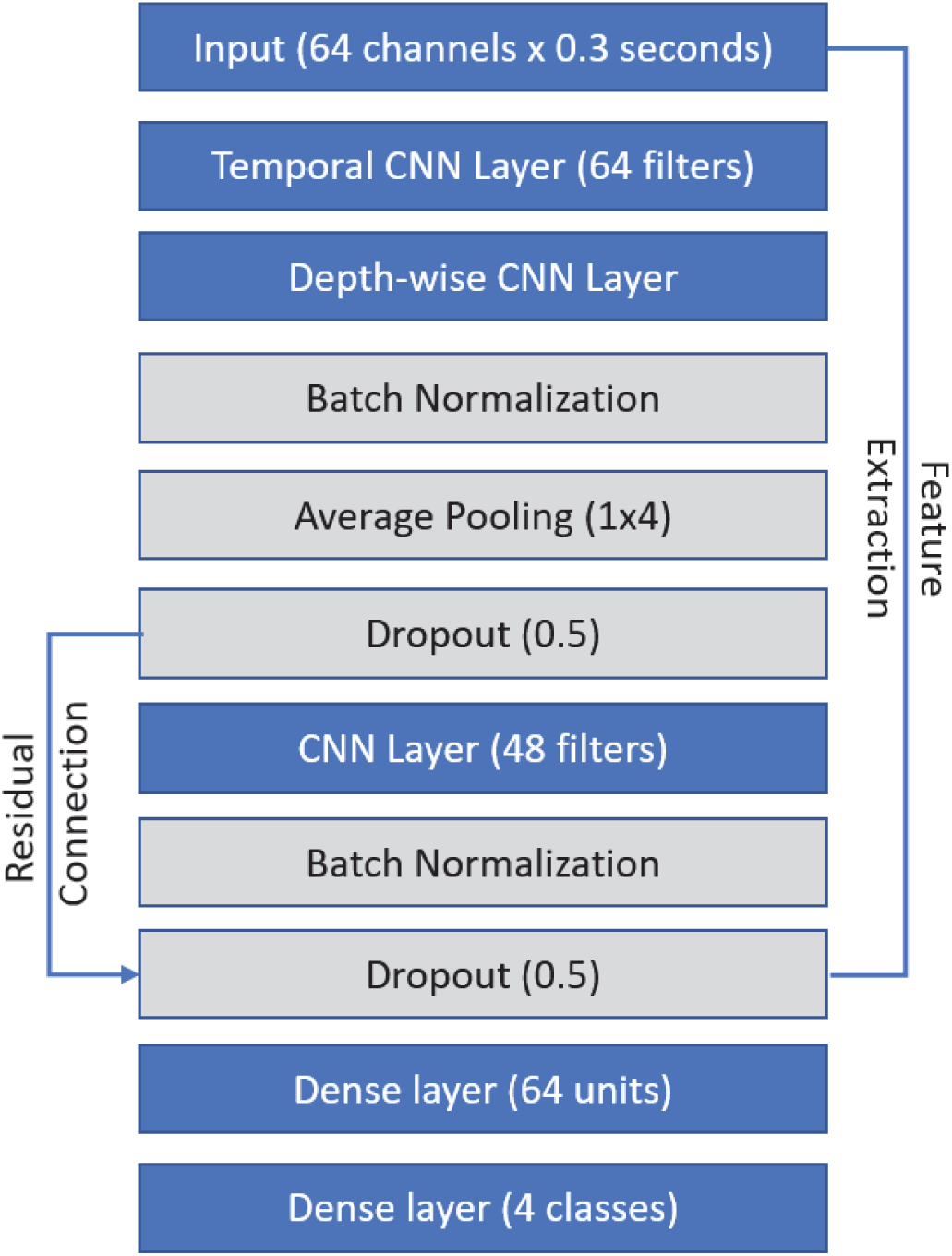
The optimized CNN architecture. Blue indicates architecture design choices and gray indicates regularization selections.

#### 4.1.2. Recurrent neural network

Table 2 presents the results from the optimization framework for the RNN architecture. Average performances across all participants are presented as performance above the chance model.

**Table 2:**
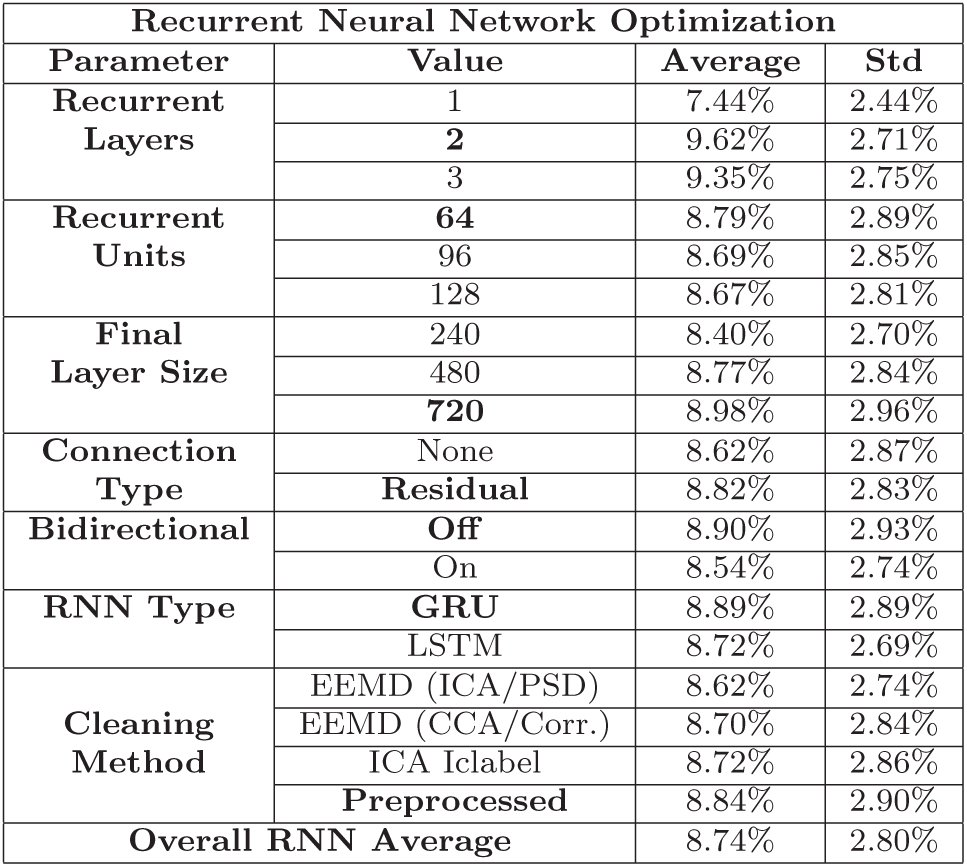
Optimization of the recurrent neural network models. Performance is presented as accuracy above the chance model.

| Recurrent Neural Network Optimization |  |  |  |
| --- | --- | --- | --- |
| Parameter | Value | Average | Std |
| Recurrent Layers | 1 | 7.44% | 2.44% |
|  | **2** | 9.62% | 2.71% |
|  | 3 | 9.35% | 2.75% |
| Recurrent Units | **64** | 8.79% | 2.89% |
|  | 96 | 8.69% | 2.85% |
|  | 128 | 8.67% | 2.81% |
| Final Layer Size | 240 | 8.40% | 2.70% |
|  | 480 | 8.77% | 2.84% |
|  | **720** | 8.98% | 2.96% |
| Connection Type | None | 8.62% | 2.87% |
|  | **Residual** | 8.82% | 2.83% |
| Bidirectional | **Off** | 8.90% | 2.93% |
|  | On | 8.54% | 2.74% |
| RNN Type | **GRU** | 8.89% | 2.89% |
|  | LSTM | 8.72% | 2.69% |
| Cleaning Method | EEMD (ICA/PSD) | 8.62% | 2.74% |
|  | EEMD (CCA/Corr.) | 8.70% | 2.84% |
|  | ICA ICLabel | 8.72% | 2.86% |
|  | **Preprocessed** | 8.84% | 2.90% |
| Overall RNN Average |  | 8.74% | 2.80% |

The optimization framework resulted in a shallow RNN with only two recurrent layers, 64 recurrent GRU units, and a final layer size of 720. The network does include a residual connection between the first and second recurrent layers. The recurrent layers do not include the bi-directional functionality. The cleaning method that resulted in the highest model performance is, similarly to the optimized CNN model, the ICA method with ICLabel EMG removal, though the difference in performances between datasets was not as pronounced as with the CNN architectures. All cleaning methods underperformed against the solely pre-processed data. The final RNN architecture is presented with Figure 10.

**Figure 10:**
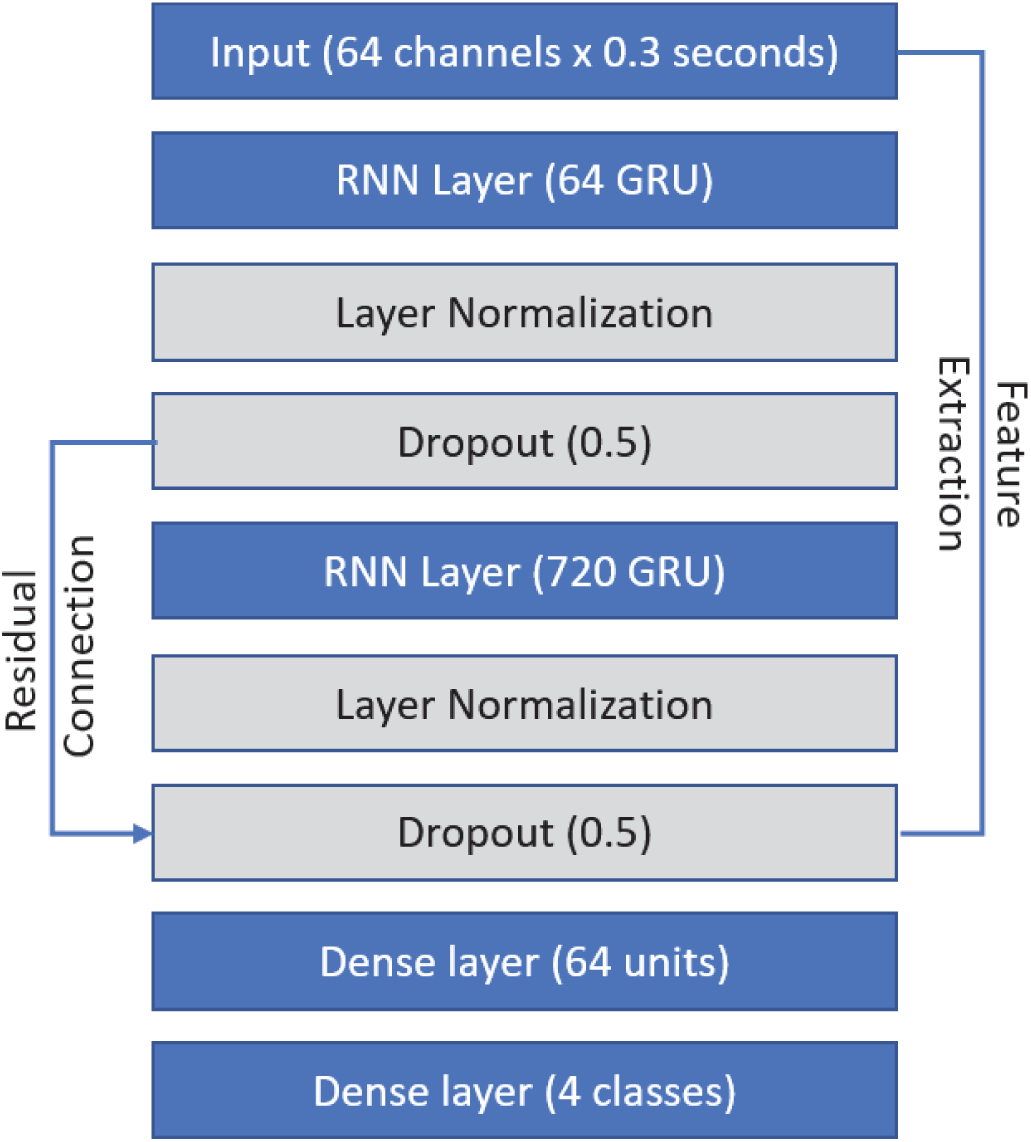
The optimized RNN architecture. Blue indicates architecture design choices and gray indicates regularization selections.

#### 4.1.3. Attention model

Table 3 presents the results from the optimization framework for the attention network. For the feature extraction layers, the optimum CNN and RNN architectures are included along with the optimum single-layer shallow CNN and RNN architectures. Average performances across all participants are presented as performance above the chance model.

**Table 3:**
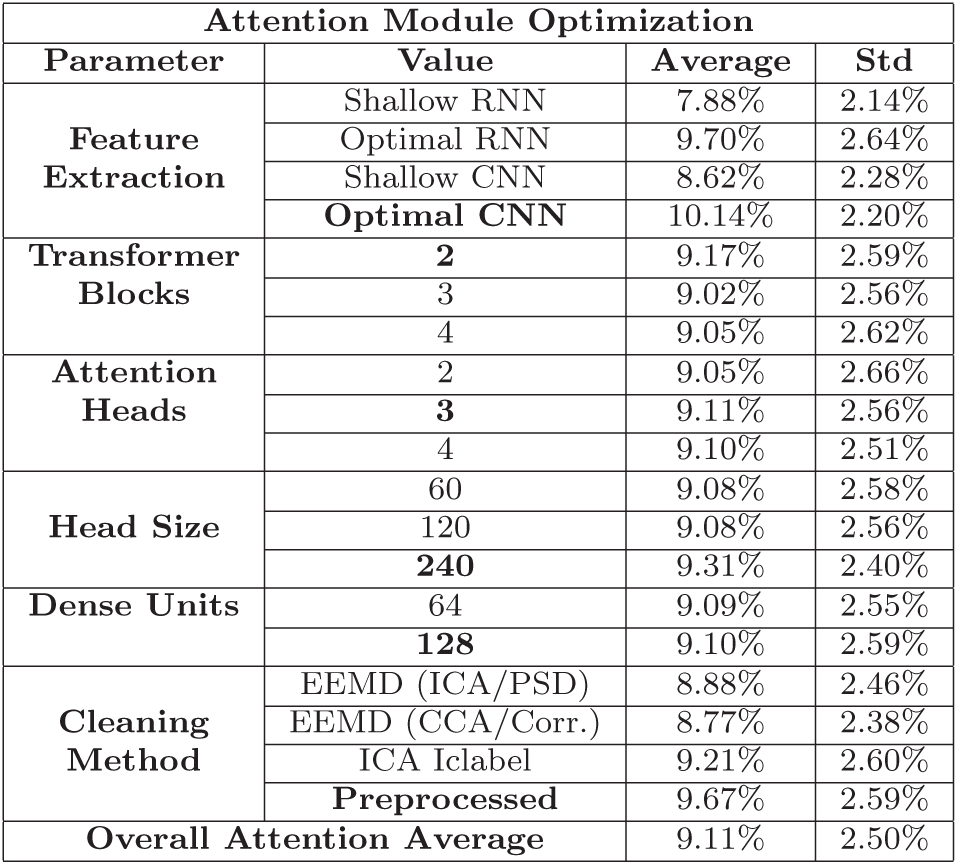
Optimization of the attention models. Performance is presented as accuracy above the chance model.

The most effective feature extraction section of the attention module was based on the optimized CNN. The optimized attention module included two transformer blocks with three attention heads each (head size of 240). The final dense layer employed 128 dense units. Like the CNN and RNN optimal architectures, the ICA dataset that employs ICLabel achieved the highest model performance when compared against the other two cleaning methods, but still underperformed against the solely pre-processed data. The optimal attention network (with the optimal CNN as the feature extraction layers) is presented with Figure 11.

**Figure 11:**
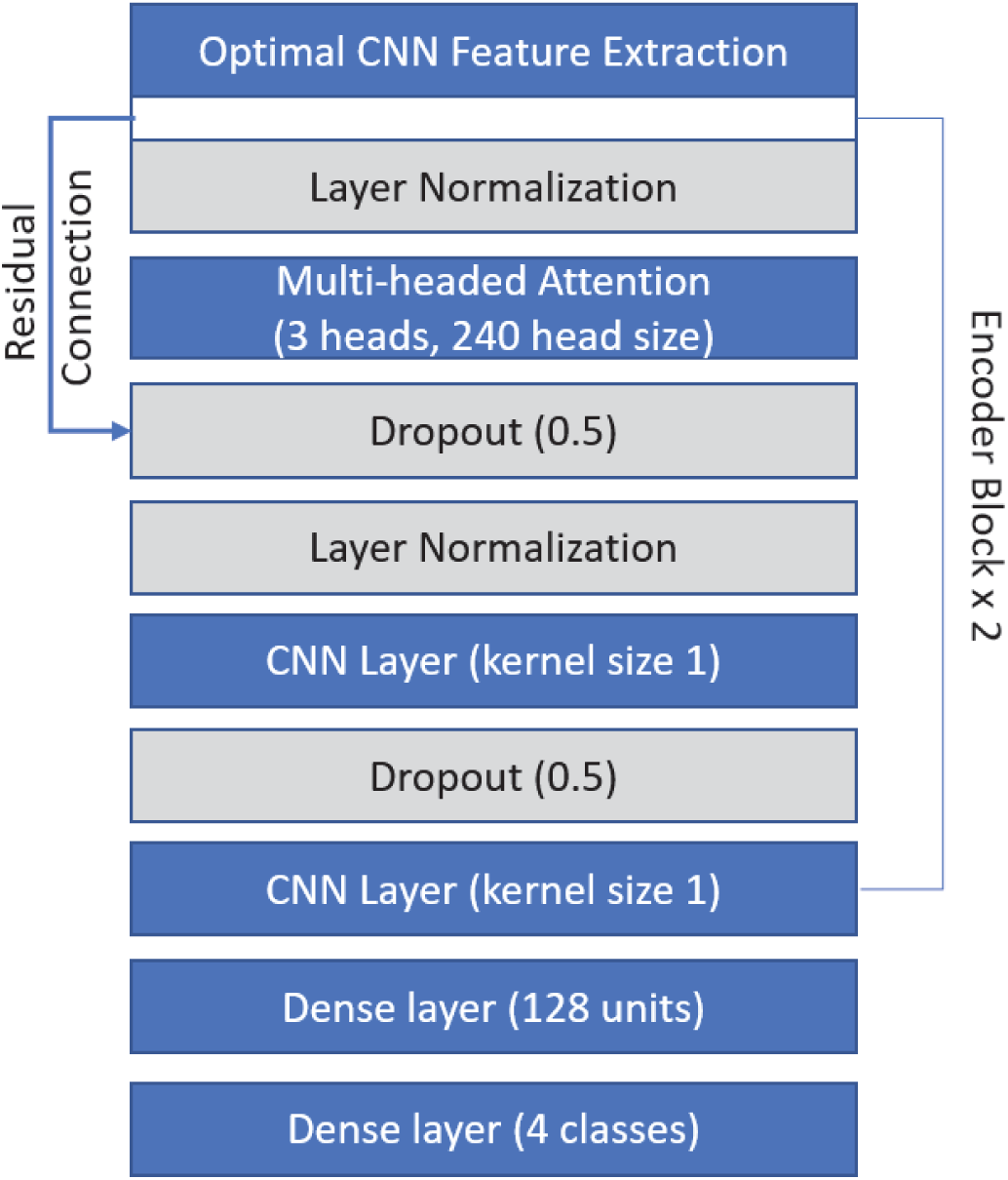
The optimized attention model. Blue indicates architecture design choices and gray indicates regularization selections.

These optimal architectures were then used as deep learning models for decoding of all discrete classes and for the decoding of continuous acoustic signal properties.

### 4.2. Discrete classification results

#### 4.2.1. Model performances with optimized architectures

The CNN outperformed the RNN and the CNN with attention for four of the six tested discrete class formations (Figure 12). For the Presence of Voice, the optimal RNN outperformed both the CNN and attention network. Models trained on the Presence of Voice class exhibited the least differentiability between the binary classes, evidenced by the relatively lower increase over the chancel model as compared to all other class formations, so the slight improvements in performance for the RNN here may be more indicative of the difficulty in decoding this specific class than on differences in model architectures. For the Place of Articulation for Vowels and Vowel vs Consonant classes, the CNN and the attention network achieved similar performance averages. However, in terms of training time and computational demands, the attention network can take up to 1000% more time, even with the use of a high performing computing cluster. For this reason, all other discrete decoding tests employed the optimized CNN architecture.

**Figure 12:**
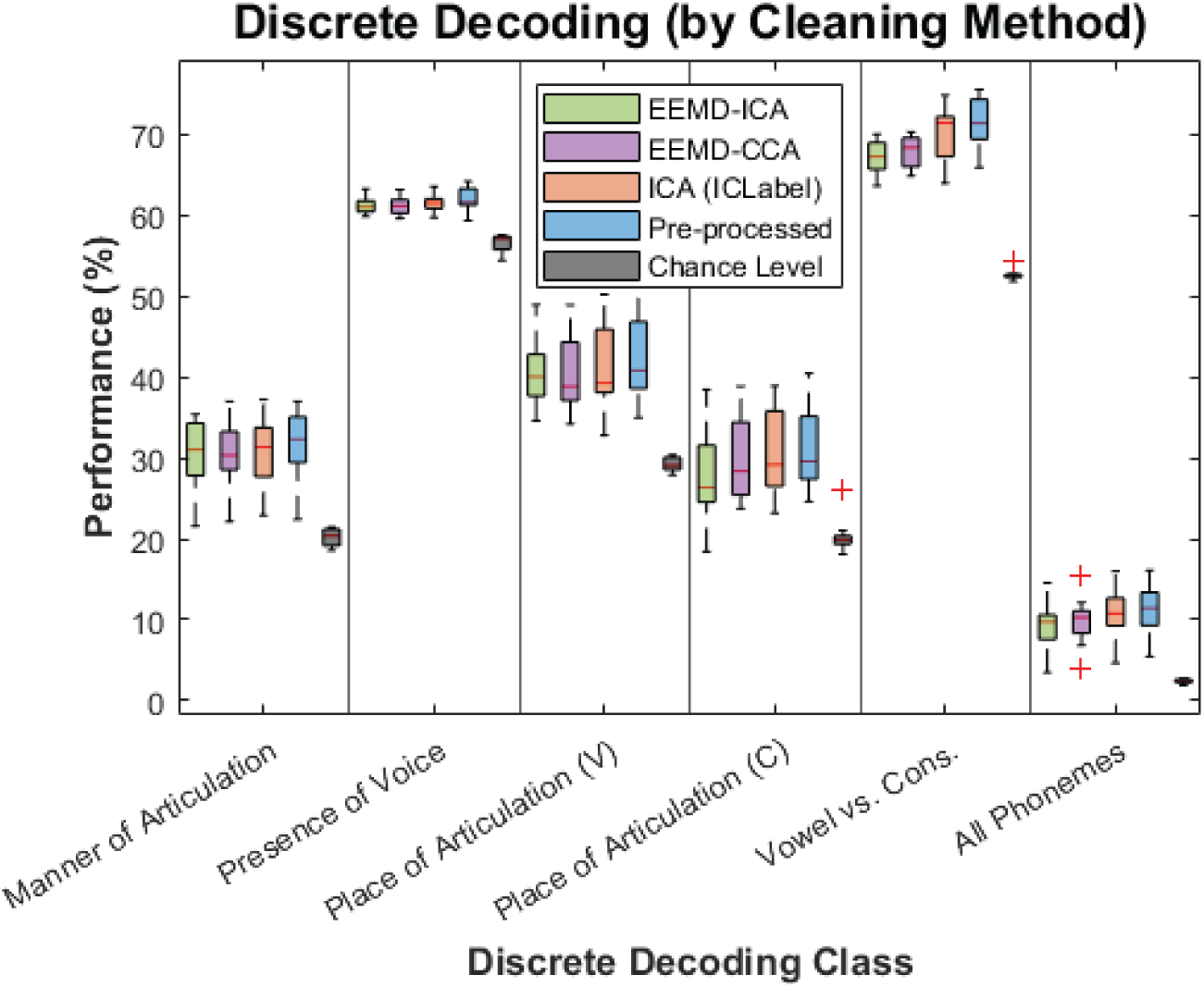
Discrete performances by optimized architecture averaged across cleaning methods as compared to the chance level.

The ICA implementation with ICLabel achieved higher model performance as compared to EEMD-ICA and EEMD-CCA for all but the Place of Articulation for Vowels class (Figure 13).

**Figure 13:**
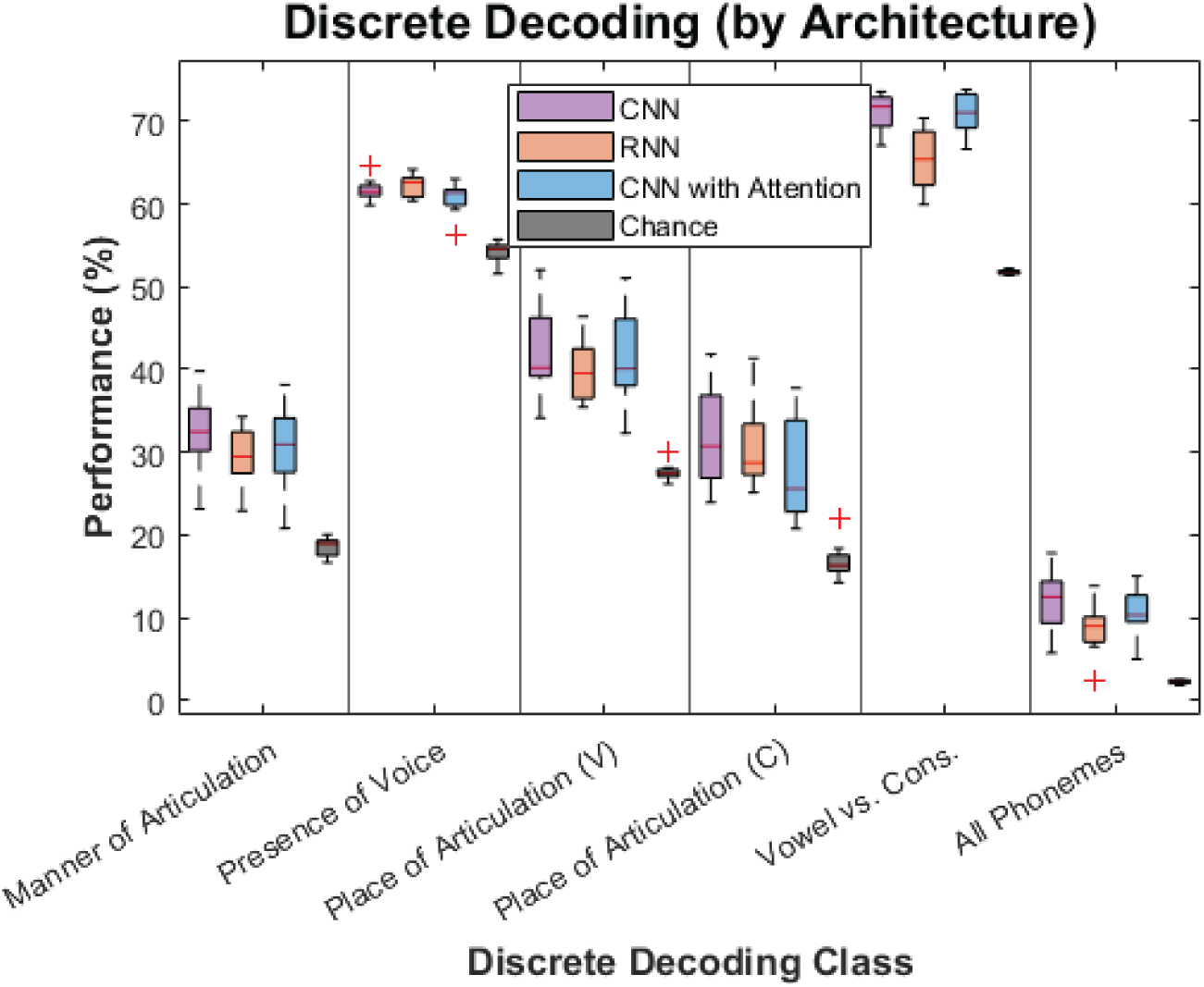
Discrete performances by cleaning method averaged across architectures as compared to the chance level.

Though the average performance of models with solely pre-processed data always outperformed the EMG-cleaned datasets, all datasets outperformed when compared against the performance of the chance model. The following confusion matrices are presented for a single participant with the ICA-ICLabel dataset for additional analysis. Since the focus of the sentence selection was on achieving a phonetic balance similar to the English language, the findings detailed below on relative levels of differentiability between phonemes or features are most appropriate for similarly designed protocols. Future research may find benefits from modifying the experimental protocol with a focus on even phonetic representation, or on selecting different neural modalities to help improve the SNR.

From Participant 1’s confusion matrix for the Manner of Articulation class (Figure 14), both Fricatives and Liquid consonants are good candidates for future speech BCI paradigms using EEG. There were not many samples of Affricates or Glide consonants, as the emphasis was on producing speech with a phonetic representation that closely matches the English language, so further data is needed to draw conclusions on those two classes. Both Nasal and Plosive consonants were misclassified significantly and were incorrectly classified as Fricatives and Liquid consonants, so the inclusion of these types of consonants with Fricatives and Liquid consonants is likely not an ideal paradigm formation for continuous production tasks. With these two relatively discriminable classes, a speech BCI that involved sequences of two phonemes could obtain up to four control signals.

**Figure 14:**
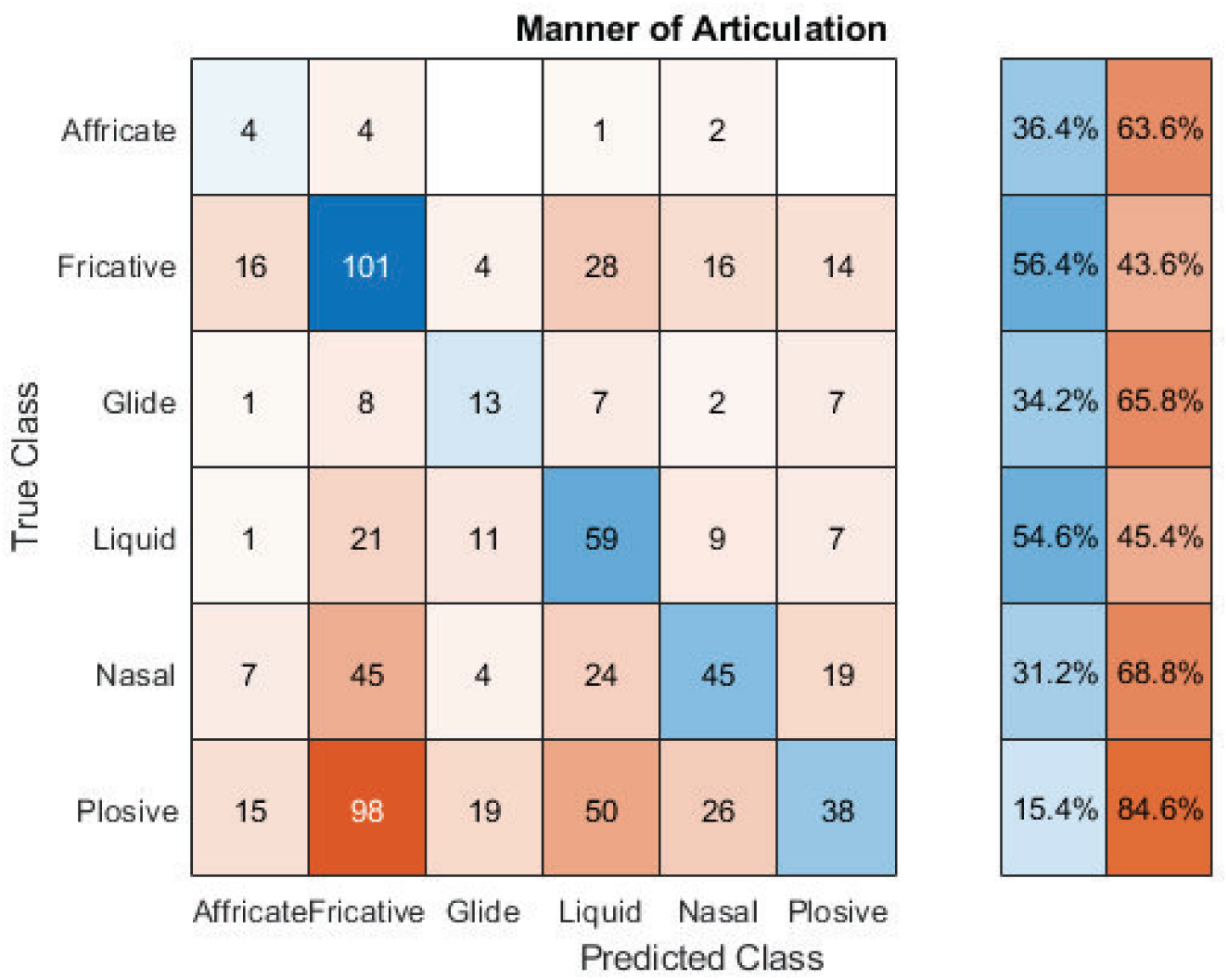
Confusion matrix for participant 1 on the ”Manner of Articulation” class for the testing data set cleaned with ICA and ICLabel.

In the resulting confusion matrix for the Place of Articulation for Vowels decoding task (Figure 15), vowels produced in the central and front locations in the mouth, along with Diphthongs, are good candidates for a speech BCI using EEG. Vowels produced in the back of the mouth are likely not good candidates, as these were misclassified 80% of the time. Sequences of three vowels would allow for nine potential classes for a speech BCI.

**Figure 15:**
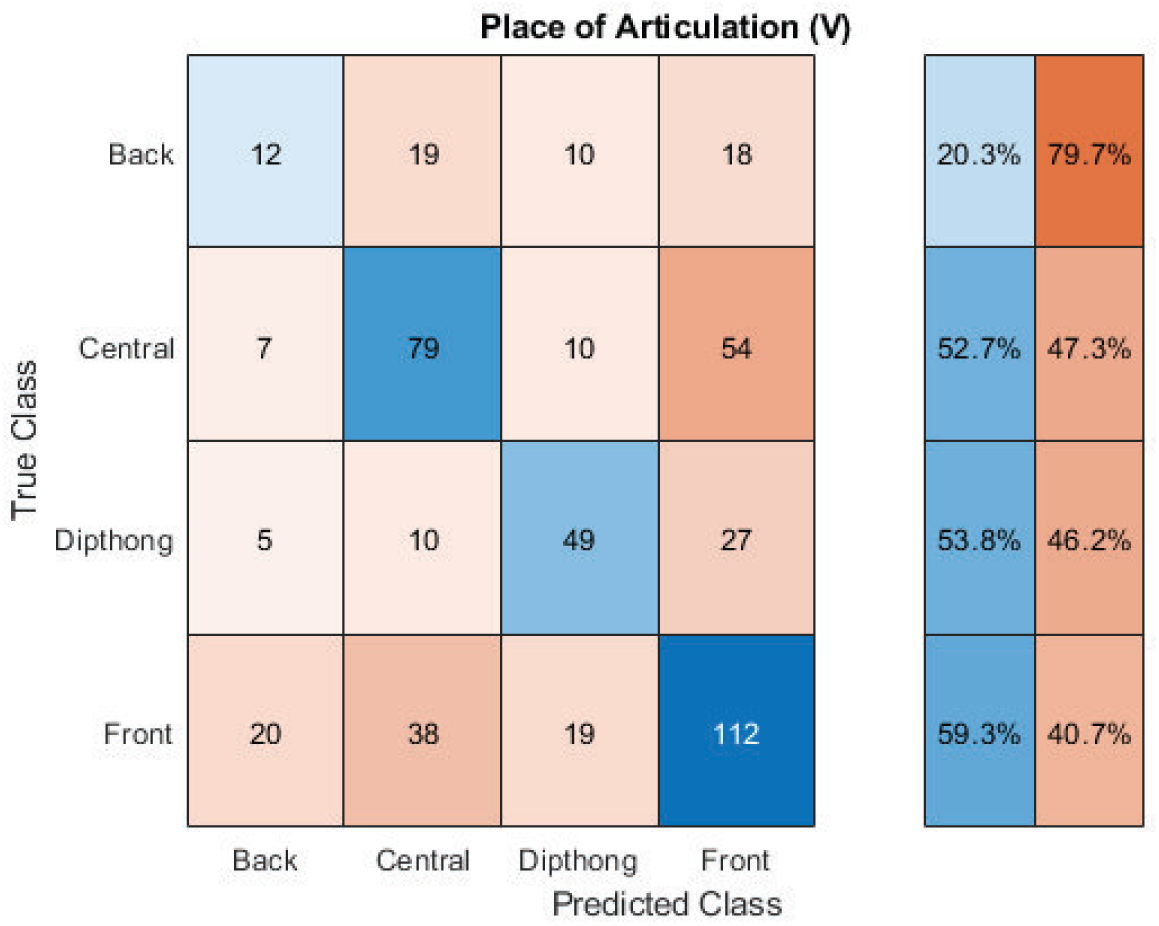
Confusion matrix for participant 1 on the ”Place of Articulation for Vowels” class for the testing data set cleaned with ICA and ICLabel.

In the resulting confusion matrix for the Place of Articulation for Consonants class (Figure 16), the CNN model correctly predicted Bilabial, Labiodental, Linguaalveolar, and Linguapalatal consonants more so than any other class, with Bilabial and Lingua-alveolar consonants showing the highest discriminative power.

**Figure 16:**
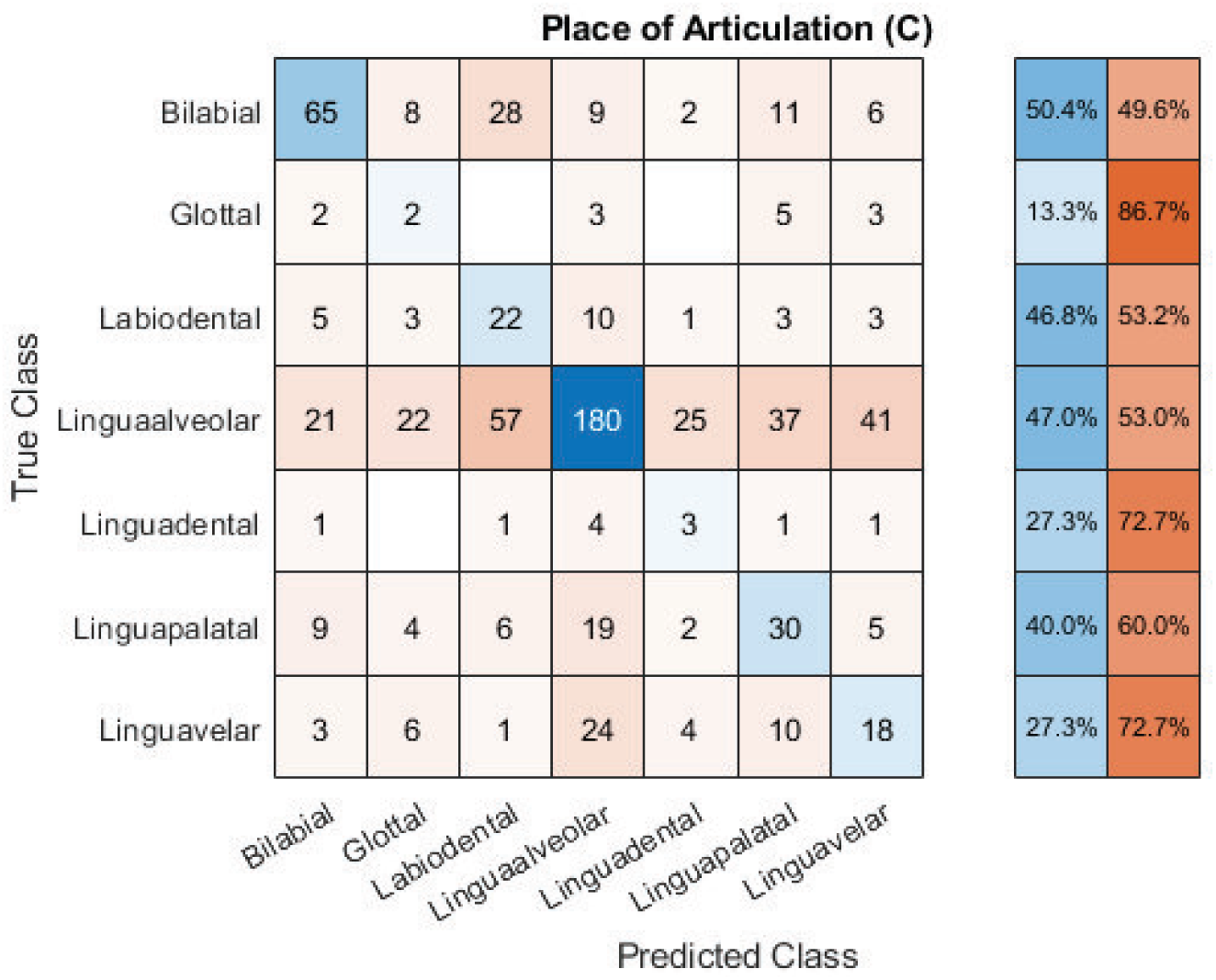
Confusion matrix for participant 1 on the ”Place of Articulation for Consonants” class for the testing data set cleaned with ICA and ICLabel.

Based on the confusion matrix (Figure 17), the CNN model was not very effective in decoding the binary presence of voice class. While the overall performance was above the chance model, the model misclassified the lack of voice more often than correctly classifying the lack of voice. The presence of voice is likely not a good candidate for an EEG-based speech BCI.

**Figure 17:**
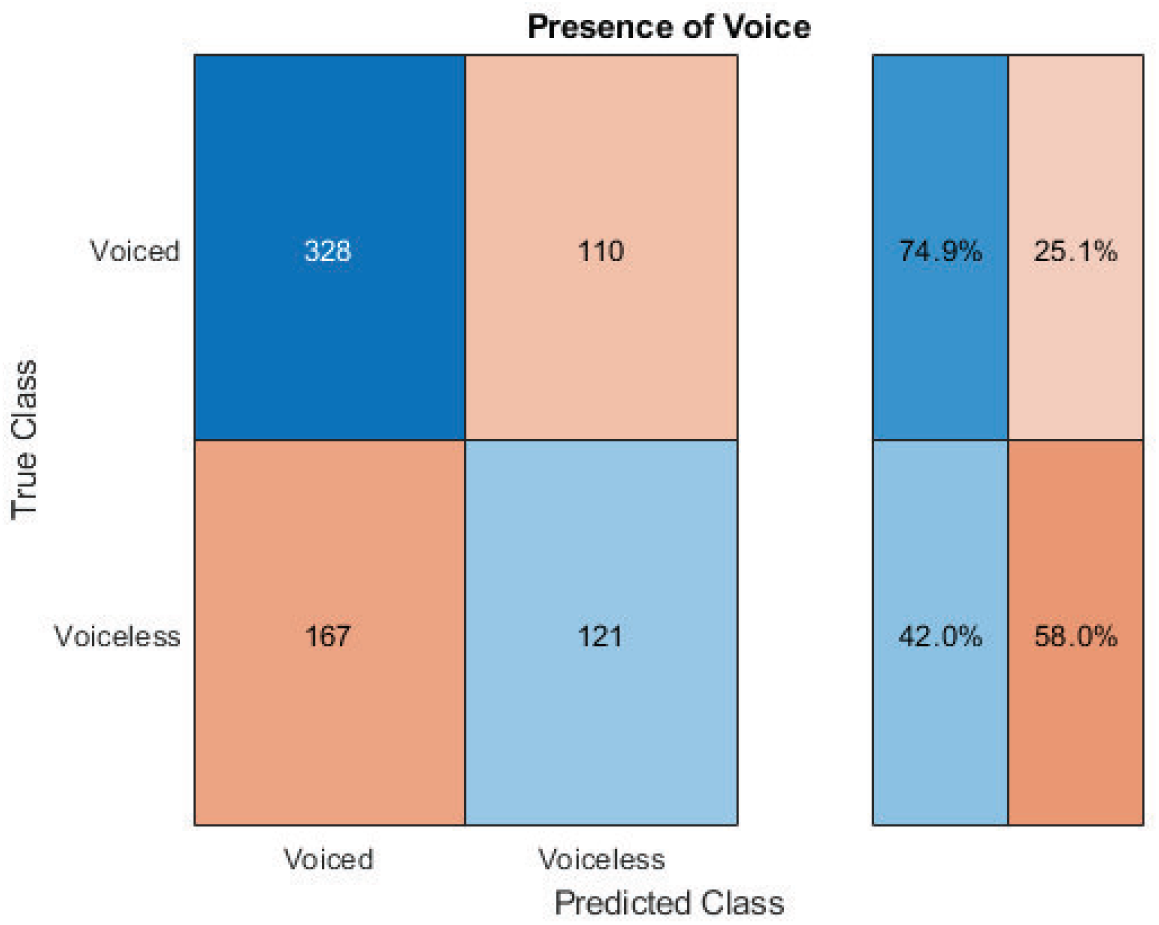
Confusion matrix for participant 1 on the ”Presence of Voice” class for the testing data set cleaned with ICA and ICLabel.

Decoding the binary vowels vs consonants class was significantly more effective than decoding the binary presence of voice class (Figure 18), with the model predicting 80% of the consonants correctly and 73.6% of the vowels correctly. These findings, along with the discriminative power for the vowels and consonants, open up the potential for a two-stage decoding mechanism, where the network first predicts whether an incoming phoneme is a consonant or vowel, before further decoding on the type of consonant or vowel.

**Figure 18:**
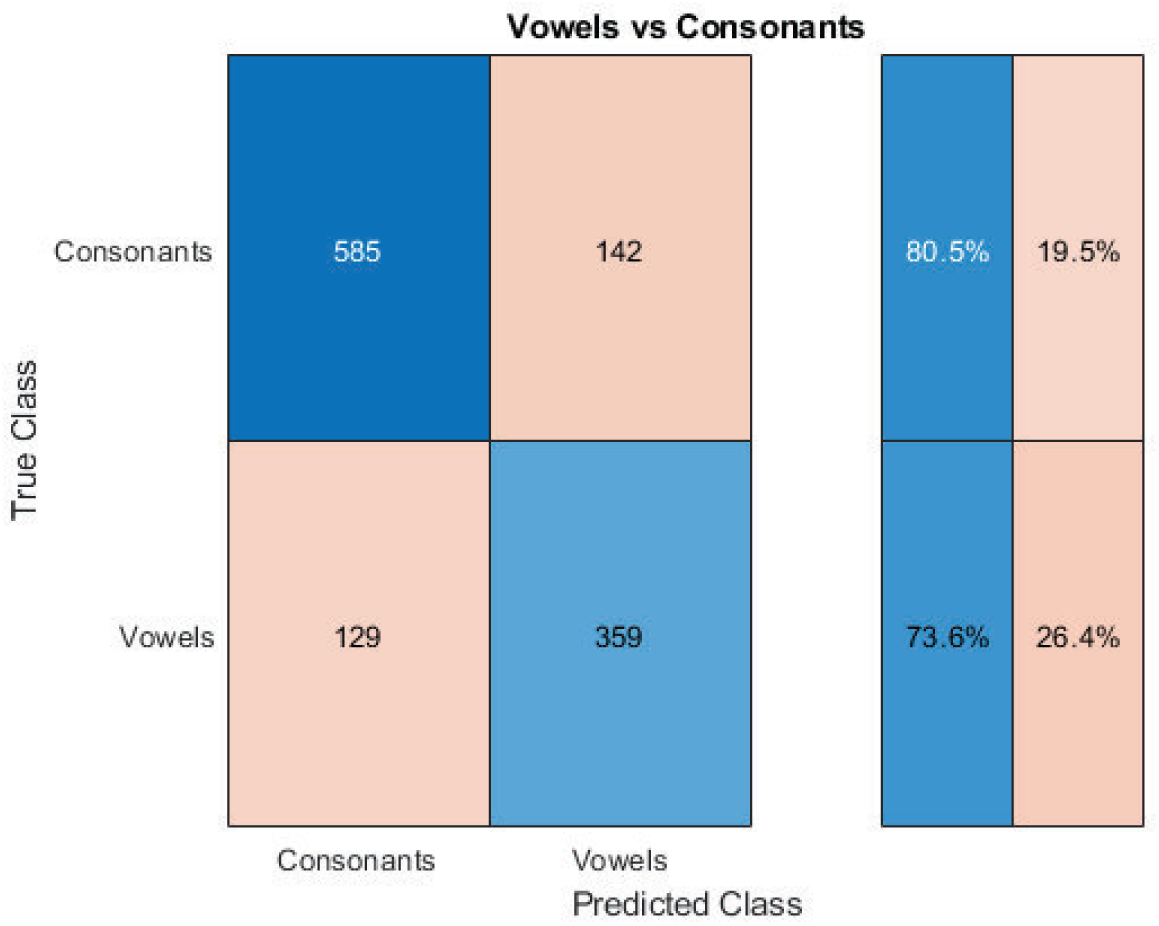
Confusion matrix for participant 1 on the ”Vowels versus Consonants” class for the testing data set cleaned with ICA and ICLabel.

While the overall model performance for the decoding of all individual phonemes was significantly above chance (19% accuracy for Figure 19 as compared to a chance level of 2.5%), the lack of representation for some phonemes makes direct comparisons of performance between phonemes difficult. The top two most distinguishable phonemes that have adequate representation are /ae/ and /ey/ (Figure ??). This is counter-intuitive as both of these vowels are produced at the front of the mouth, but promising since each of these vowels was rarely misclassified as the other. These two vowels could be used in conjunction with a consonant such as /s/, as /s/ had a relatively high correct classification rate and, more importantly, was never misclassified as either /ae/ or /ey/.

**Figure 19:**
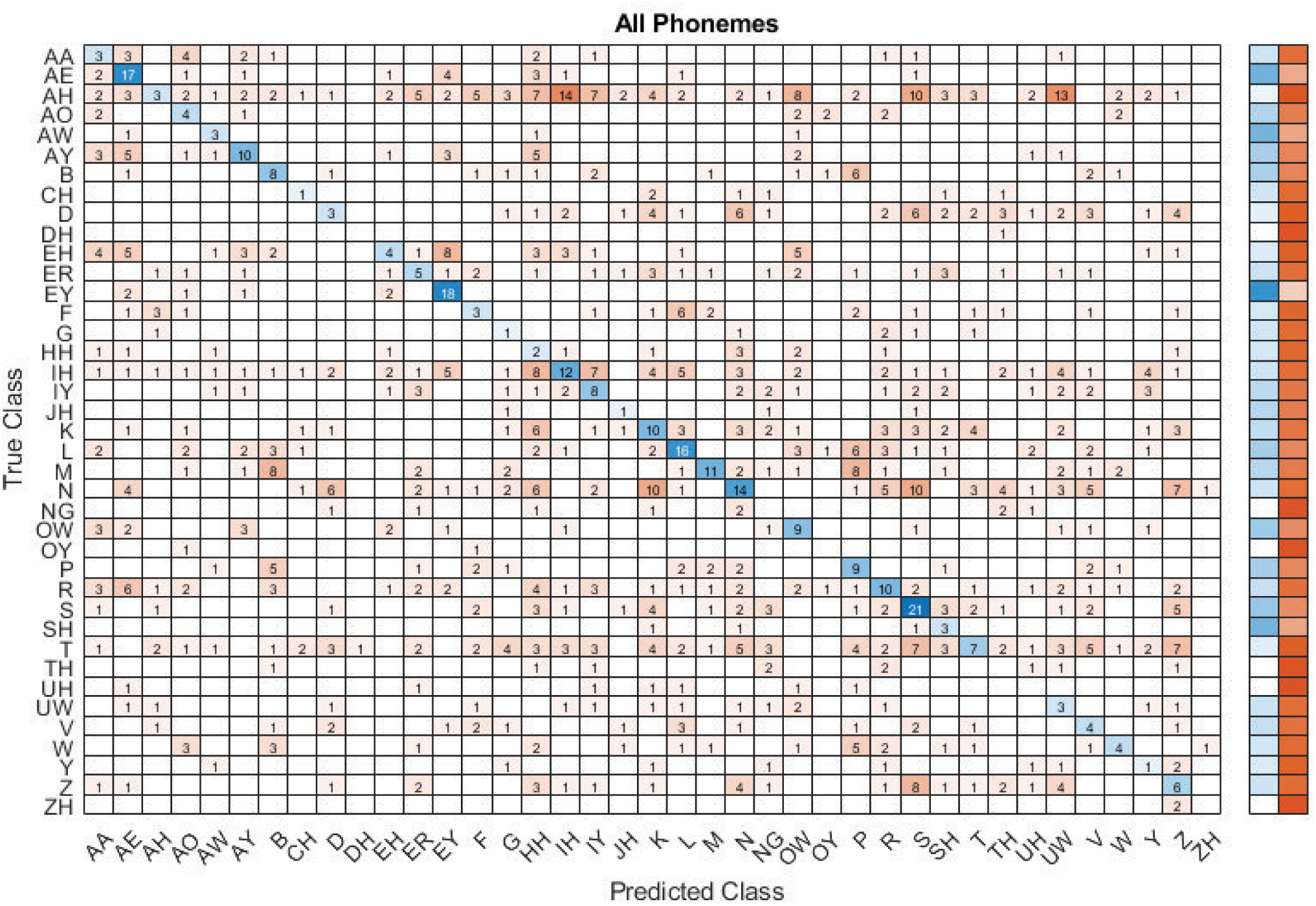
Confusion matrix for participant 1 on all phonemes for the testing data set cleaned with ICA and ICLabel.

#### 4.2.2. Frequency analysis

In the analysis of frequency sub-bands of importance, there were consistent trends across all discrete decoding tasks. The delta, theta, lower gamma, and upper gamma all exhibited higher performances when isolated as compared to the alpha and beta bands, with these bands barely outperforming the chance model (Figure 20). For the delta and theta bands, the performance averages were very similar when comparing the EMG-cleaned and solely pre-processed data, which was expected as the goal for EMG removal processes was on removing high frequency EMG content while preserving low frequency EEG content. The significance of low frequency information to speech processing with EEG is an understood phenomenon, with research pointing to both the Delta [76] and Theta bands [77] as being important.

**Figure 20:**
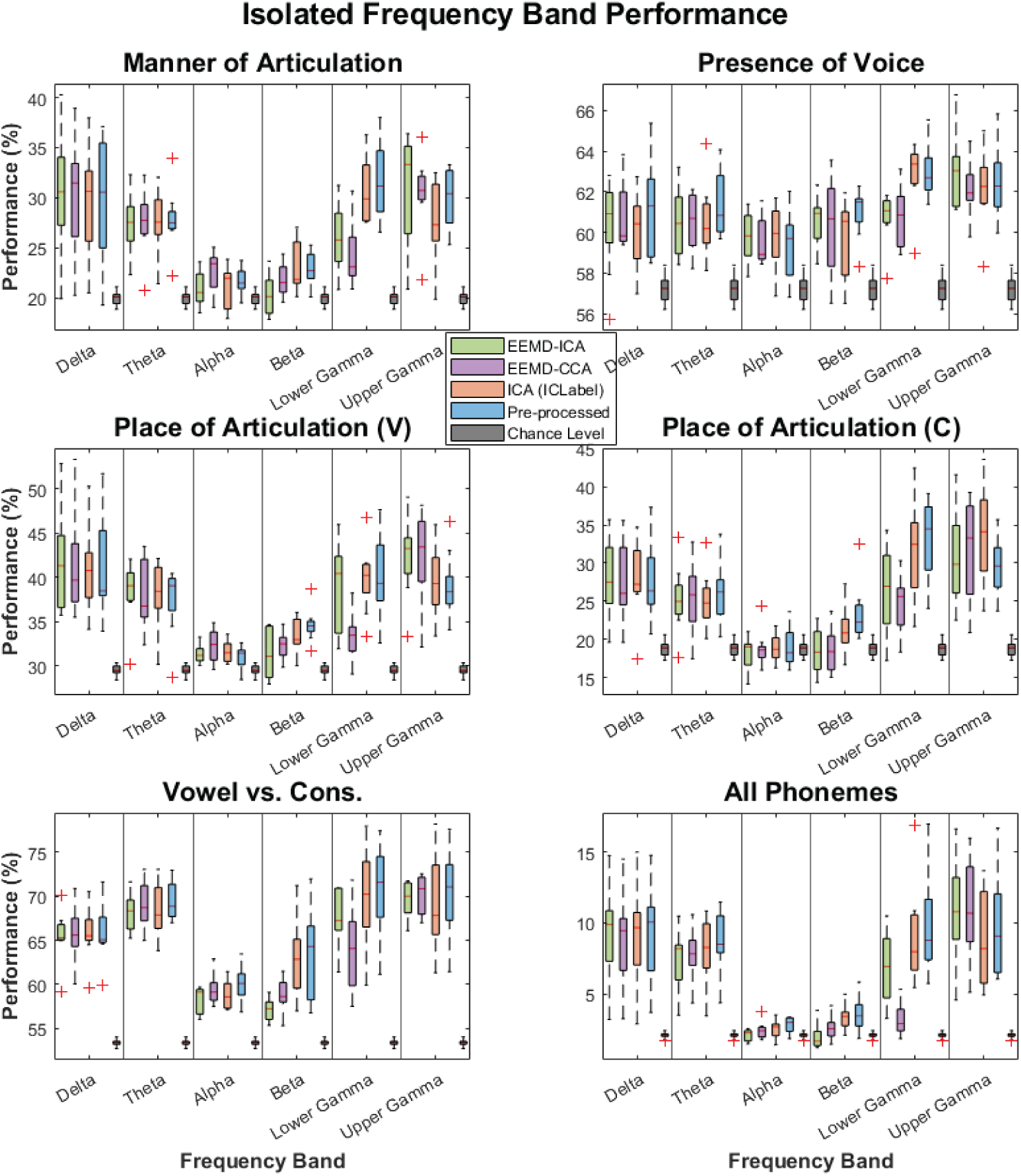
Discrete performances by isolated frequency band.

There were significant differences between cleaning methods when isolating the gamma bands. For lower gamma, the two-stage EMG removal methods both saw significant reduction in performance as compared to the ICA-ICLabel and solely pre-processed data, whereas for the upper gamma band this trend is not as evident. For all classes in the lower gamma band, the EEMD-CCA cleaning method performed worse than the EEMD-ICA cleaning method. This points to the likelihood that EEMD-CCA was more effective in removing lower gamma content related to EMG noise and that, for the remaining datasets, the trained models were potentially focusing on the information in the lower gamma bands due to the higher relative retention of EMG noise. Surprisingly, this trend did not hold for the upper gamma band, potentially indicating that, even though the EEMD-CCA and EEMD-ICA methods removed up to 99% of the power in the upper gamma band, the model was still able to use this remaining frequency information for effective decoding.

#### 4.2.3. Channel of importance analysis

In the perturbation analysis (Figure 21), there were some consistent trends across cleaning methods and discrete decoding tasks. On average, for the decoding of All Phonemes and Place of Articulation for Consonants tasks, all datasets saw decreased performance when removing the FP2 channel. For the Manner of Articulation for Consonants class, the three cleaned datasets highlighted FP1 and/or FP2 as important for performance, but the solely pre-processed data did not, which is counterintuitive since these locations were found to have the highest amount of power in the reconstructed artifact, which implies a greater level of contamination had initially been present at these locations. An expectation here was that the solely pre-processed data would gain more information important for decoding from the heavily contaminated electrodes, but this finding suggests the opposite. When discriminating between vowels and consonants, none of the three cleaning methods focused on the frontal channels, instead highlighting channels that overlay the temporal lobe. As this class had the highest difference between model performance and chance level, this could indicate that the discrimination of vowels and consonants does not rely on the EMG contamination as compared to the other class formations. For the class that attempts to predict the presence of voice, channels that led to the largest decreases in performance were located over the occipital and parietal lobes, a trend not observed with any other class formation. Since the presence of voice indicates whether the vocal folds are vibrating, it’s possible that this vibration led to frequency contamination present in the neck. Further analysis is needed to fully understand why the occipital and parietal lobes were relatively more important for decoding the presence of voice as compared to the other class formations.

**Figure 21:**
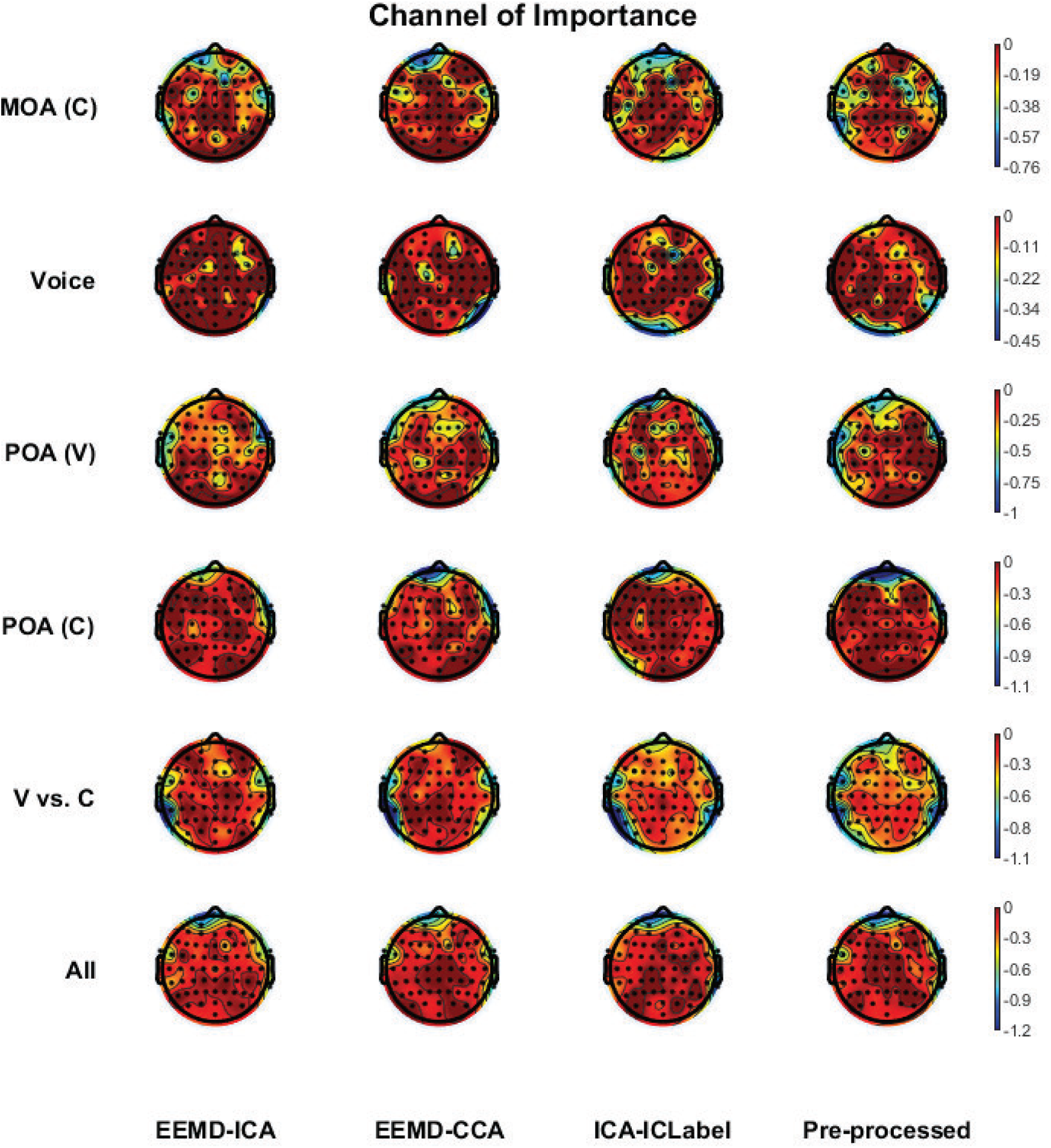
In this perturbation analysis, areas of the scalp that are blue indicate channels that, when removed, led to larger decreases in model performance as compared to the model trained with all channels active. The color scales represent percentage decrease in accuracy following the removal of a single channel.

The above perturbation analysis (Figure 21 indicated that FP1 and FP2 were still considered important channels for the decoding of four classes: Manner of Articulation, Place of Articulation for Vowels and Consonants, and the classification of individual phonemes. To assess the effects of removing these channels, models were again trained for all classes for all participants with these channels excluded. The assumption here was that, for classes that identified either FP1 or FP2 as important for decoding, the resulting models would exhibit poorer performance as compared to full-channel performances. This analysis is presented with Figure 22.

**Figure 22:**
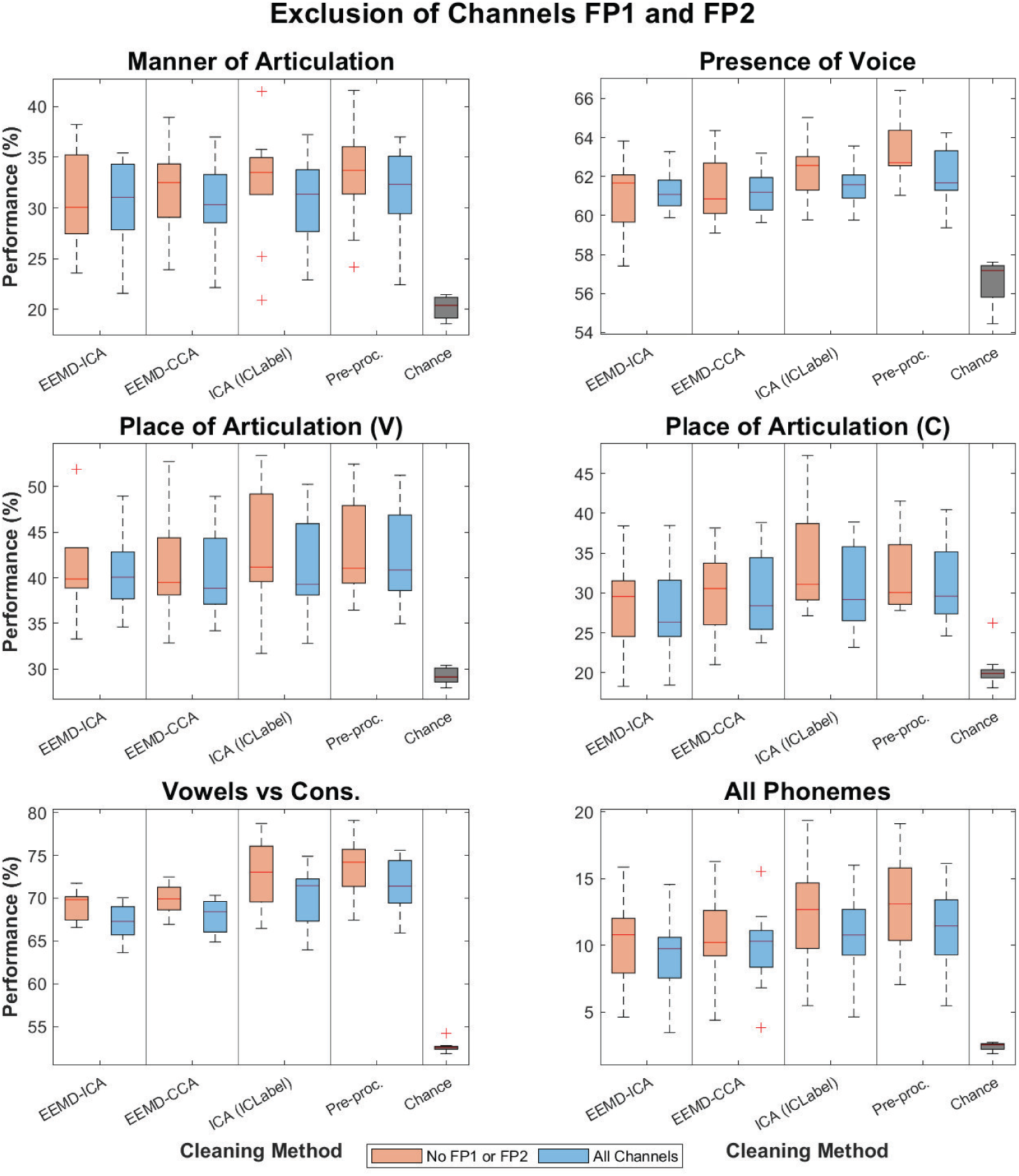
Models trained with the exclusion of channels FP1 and FP2 appear to have improved the performance of models against All Channel performances.

The results detailed in Figure 22 were contrary to our initial hypothesis. The exclusion of FP1 and FP2 from the training set actually led to relatively consistent improvements in performance as compared to the full-channel performances, both for the classes that identified FP1 or FP2 as important, but also for the other two classes.

The top performance among all participants for the classification of individual phonemes was 15.9% (chance level 2.8%), but, in this case with channel exclusion, this performance rose to 18.5%. These improvements may indicate that the models trained with all channels focused on FP1 and FP2, likely due to the remaining EMG contamination in these frontal channels, at the detriment of searching for more informative patterns in other channels. Since only two channels were removed, it is likely not due to the smaller size of the input data, though the removal of a larger number of channels was investigated in Section 4.2.4.

#### 4.2.4. Channel Selection Results

This analysis of channel selection methods included five different types of channel selections: manual selection based on past research, class and cleaning method independent CSP-guided channel selection, CSP-guided channel selection averaged over discrete decoding class, CSP-guided channel selection averaged over cleaning method type, and the overall average CSP-guided channel selection.

The resulting channel selection averaged over cleaning methods (Figure 23) led to a surprising finding. In Sectionof the Appendix, the reconstructed artifact resulting from each cleaning method found the highest power was removed from either FP1 and FP2, indicating more EMG noise was originally present at these sensors. However, FP2 was selected through this channel selection approach for each of the cleaning methods, whereas the solely pre-processed data did not choose this sensor with this CSP channel average. This finding matched the perturbation analysis for pre-processed data, and may further support the idea that EMG contamination was too high for the model to effectively decode, and that the significant reduction in high frequency power at this electrode for the assessed cleaning methods allowed for effective decoding.

**Figure 23:**
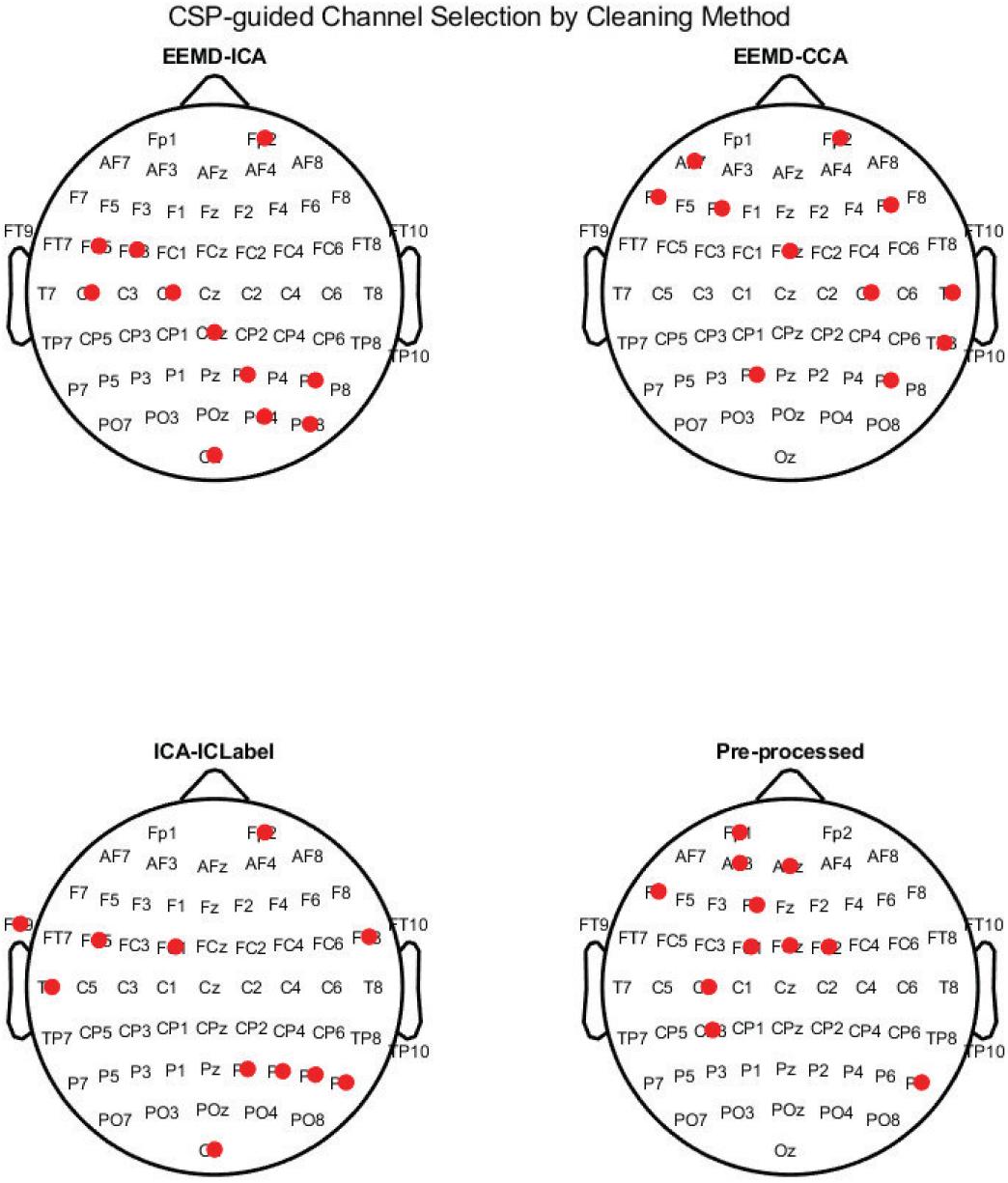
CSP-guided channel selection averaged across cleaning methods.

There were not many similarities between the CSP-guided channel selection method averaged over discrete decoding class (Figure 24). Channel FP1 was selected through this method for the Presence of Voice, Place of Articulation for Vowels, and Vowels vs. Consonants classes, whereas in the perturbation test for these three classes, only the Place of Articulation for Vowels highlighted either FP1 or FP2 as important channels for decoding performance. For the Vowels vs. Consonants class, the perturbation analysis pointed towards the temporal and parietal channels on the left hemisphere as being important, but the class-average CSP-guided selection for this class did not. This may indicate CSP-guided channel selection methods averaged over class type was not as effective for channel selection as compared to individually based channel selection or channel selection averaged over cleaning methods.

**Figure 24:**
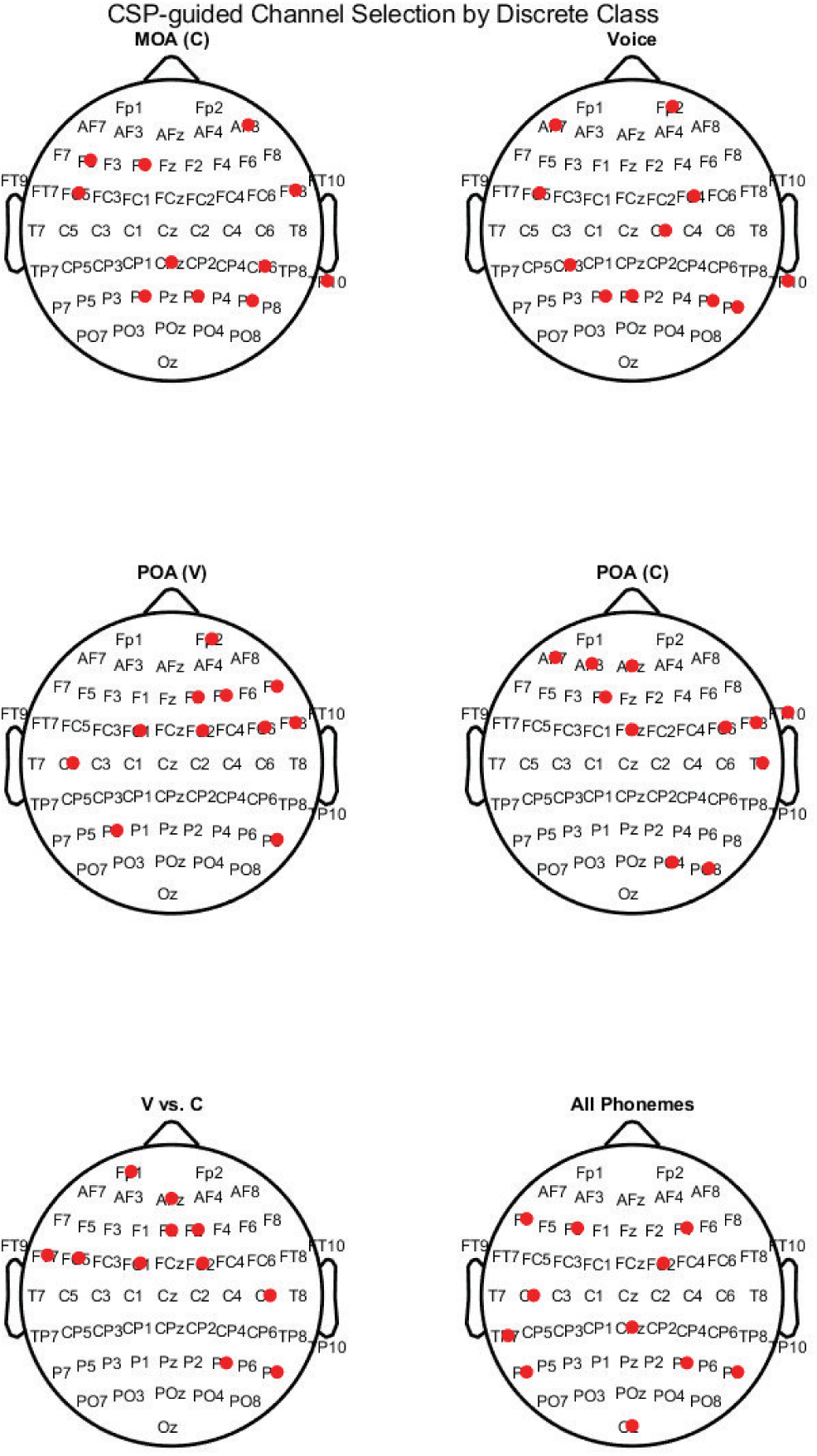
CSP-guided channel selection averaged by discrete decoding task.

The overall average CSP-guided channel selection (Figure 26) has only three channels that matched the literature-based manual selection (Figure 25) (T7, C5, and CZ), which corresponds to the middle temporal gyrus, secondary auditory cortex, Wernicke’s area, primary auditory cortex, and post-central gyrus. F3 was also selected by the average CSP-guided channel selection process, which is near F5 and F7, the two electrodes originally selected for local proximity to Broca’s area. While promising, this channel selection also included FP2 and AFz, channels in close proximity to the face, and this inclusion may have been due to the remaining EMG contamination in the data. The resulting model performance comparison supported the notion that the class average and overall average may not be effective channel selection methods, as the class-averaged CSP-guided selection and overall CSP-guided selection underperformed when compared against the individual or cleaning method-average CSP channel selection. This is counter-intuitive as the CSP approach finds important channels based on the output decoding classes, so the expectation was that an average over discrete decoding task would result in an effective selection, regardless of the type of employed EMG cleaning method. Overall, with very few exceptions, the use of all channels outperformed the channel selection methods (Figure 27). The potential reason for this is in the design of the CNN architectures, which employed a depth-wise convolutional layer to help reduce the parameter count. While effective in reducing parameter count, the lack of parameters in this spatial convolution may have reduced the capacity of the network to focus on spatial relationships between electrodes. However, all channel selection methods outperformed the chance model, indicating that, for channel-restricted clinical or commercial applications, speech BCI systems can still achieve higher than chance performance, even with a reduced subset of electrodes, and the reliance on high-density EEG systems may not be warranted.

**Figure 25:**
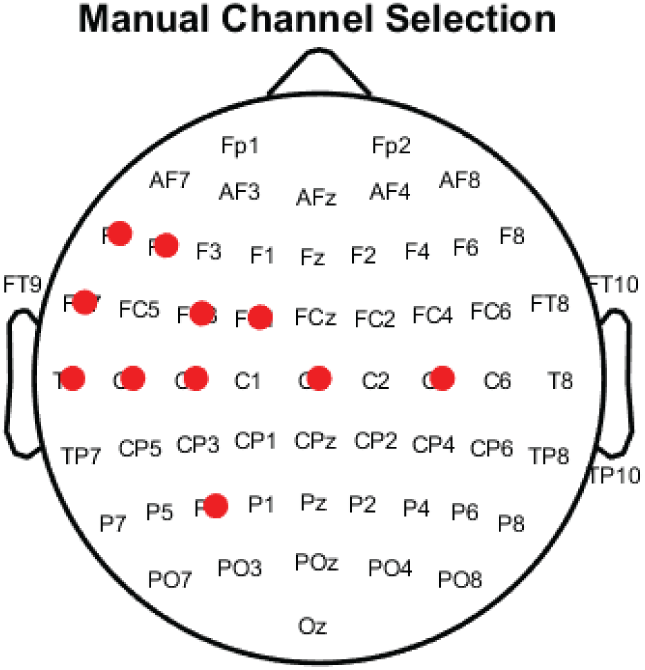
Manual selection of speech-related channels.

**Figure 26:**
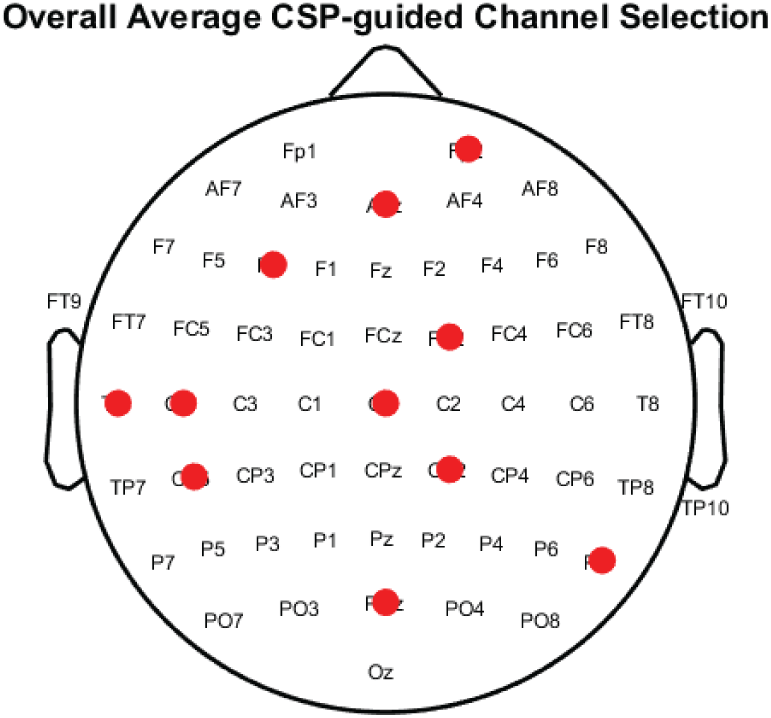
CSP-guided channel selection overall average.

**Figure 27:**
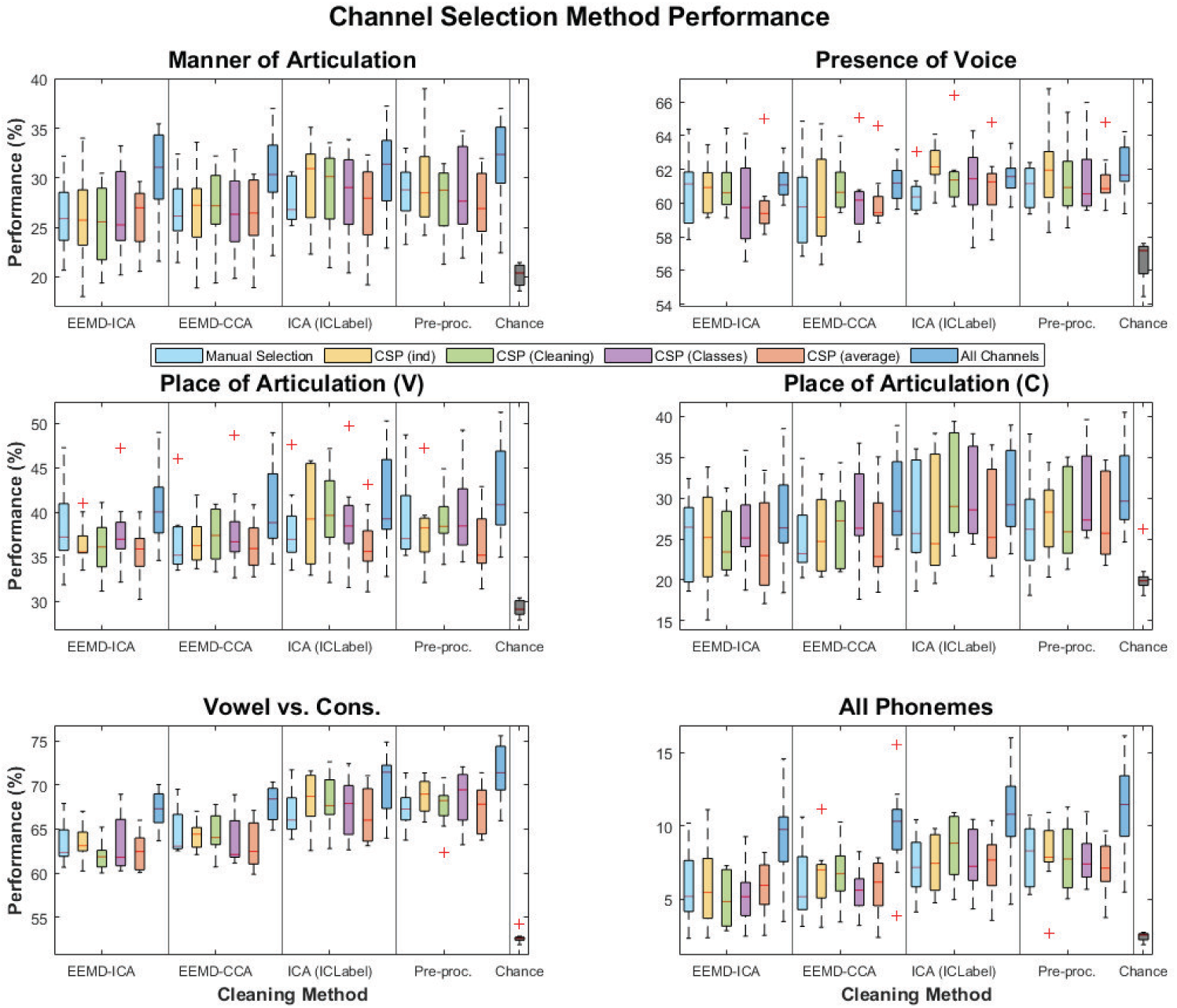
Model performances by channel selection method.

#### 4.2.5. Leave-One-Out testing results

The Leave-One-Out (LOO) testing strategy results are presented with Figure 28. In this analysis, model performances for strategies that did not use the Participant of Interest’s (POI’s) data in the training dataset (POI Not Included and POI Included (Val)), only outperformed chance level for the discrete classes ”Place of Articulation” and ”Vowels vs. Consonants”. When the first collected block of the POI’s data was included in the training set and the second block was included in the validation training set (strategy POI Included (Train)), the resulting trained models were able to outperform the chance level for all but the Presence of Voice class. In the Presence of Voice class, only the pre-processed data and the data cleaned with ICLabel were able to outperform against chance for the strategy that included some data from the POI in the training dataset. This LOO result makes sense as the Presence of Voice found the highest performance differential as compared to chance when the upper gamma frequency band was isolated. The relatively high gamma content for the pre-processed data and the ICLabel data, as compared to the two two-stage cleaning methods, may indicate that this particular class relied more so on higher frequency data than other classes. For the LOO strategy that employed the first POI block of data for the training set and the second POI block of data for the validation set, model performances for all classes and cleaning method types outperformed chance level, but did not exceed the single participant model average. This is promising as this may help to reduce the reliance on collecting large amounts of data for every participant. The final LOO strategy involved using all the POI’s data with a train/val/test split identical to the participant-independent training. With this strategy, the resulting performance exceeded the participant-independent model performance for all discrete classes and cleaning method types. This is particularly promising, as this result indicates there are common speech neural correlates that can be utilized to improve the performance over the participant-independent model, even without manual grouping of participants based on shared EEG data similarities.

**Figure 28:**
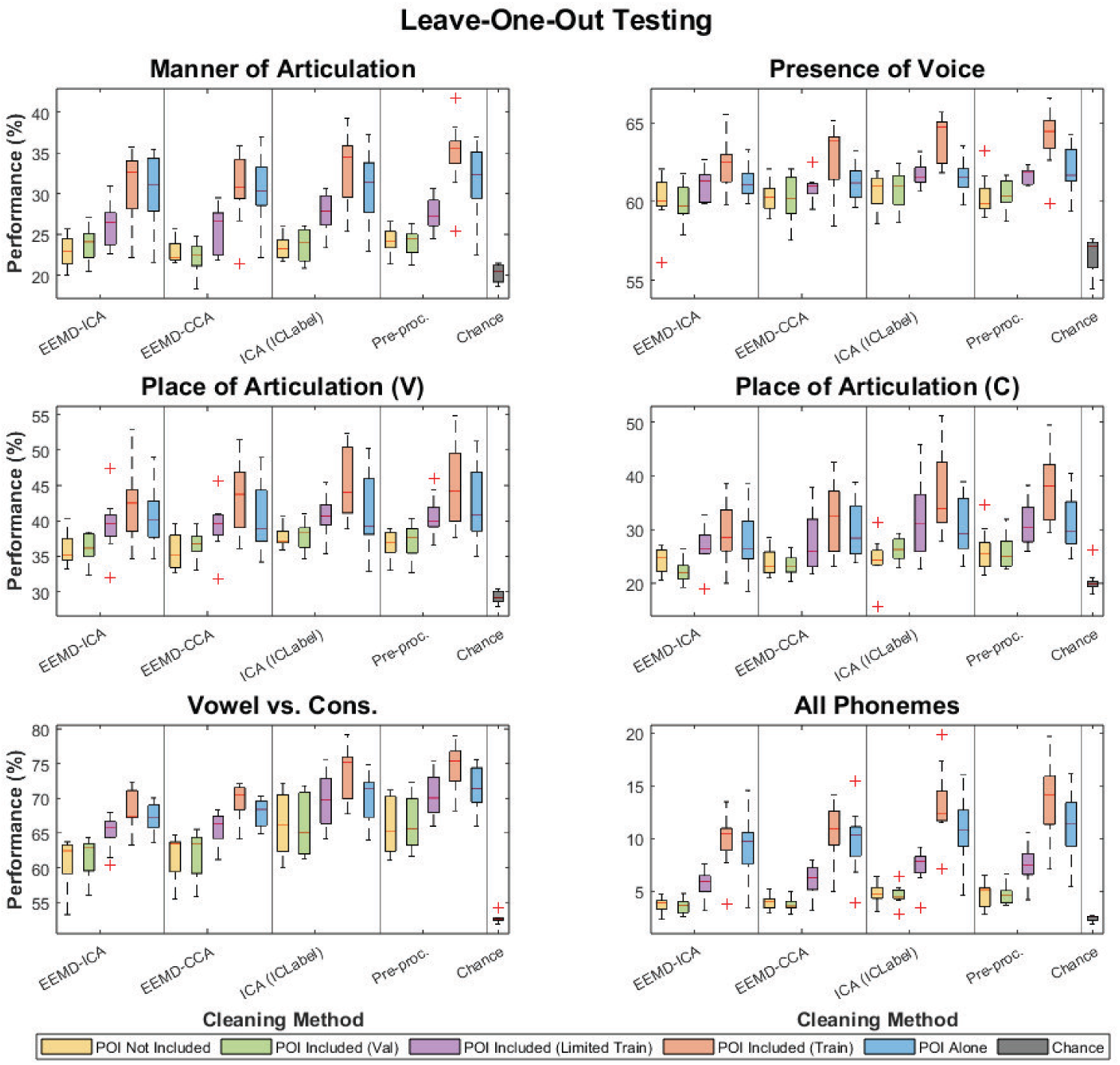
Model performances by Leave-One-Out (LOO) testing strategy.

#### 4.2.6. Continuous decoding results

The continuous decoding results are presented with Table 4. Resulting MCD metrics were compared against a chance model as created with SKlearn’s Dummy Regressor with a mean strategy. For this strategy, the chance model predicted the mean MFCC data of the training data. The below table presents the average MCD following three training sessions. As compared to [3], where the authors achieved between 5.5 and 7.1 MCD with the direct neural-to-MFCC decoder, the MCD values achieved in this research are significantly higher, indicating poorer speech synthesis. However, decoding performance for every participant was above chance, indicating acoustic characteristics of the produced audio signal can be decoded from non-invasive EEG.

**Table 4:** Decoding performance for the prediction of Mel-frequency Cepstral Coefficients (MFCCs). Performance is presented with Mel-Cepstral Distortion (MCD).

| Ptpt. | Mel-Cepstral Distortion per Participant |  |  |  |  |
| --- | --- | --- | --- | --- | --- |
|  | EEMD-ICA | EEMD-CCA | ICA (ICLabel) | Pre-processed | Chance |
| **1** | 30.79 | 30.48 | 28.97 | 28.76 | 32.38 |
| **2** | 27.19 | 27.49 | 26.76 | 26.66 | 29.06 |
| **3** | 22.67 | 22.69 | 22.14 | 22.36 | 24.23 |
| **4** | 26.03 | 25.83 | 25.83 | 25.61 | 26.88 |
| **5** | 27.64 | 27.39 | 25.78 | 26.18 | 29.11 |
| **6** | 27.90 | 27.81 | 27.05 | 27.11 | 29.59 |
| **7** | 25.04 | 24.93 | 24.60 | 24.59 | 26.10 |
| **8** | 23.52 | 23.54 | 23.03 | 23.40 | 25.49 |
| **9** | 30.91 | 30.41 | 29.06 | 29.15 | 32.47 |
| **Avg.** | **26.85** | **26.73** | **25.91** | **25.98** | **28.37** |

As with the discrete decoding, the performance of the model in predicting the continuous MFCC features was lowest (lower MCD is better) for the CNN and Attention networks (Figure 29). The average MCD for all model types outperforms the chance level, and, in fact, MCD’s for every participant were better than chance.

**Figure 29:**
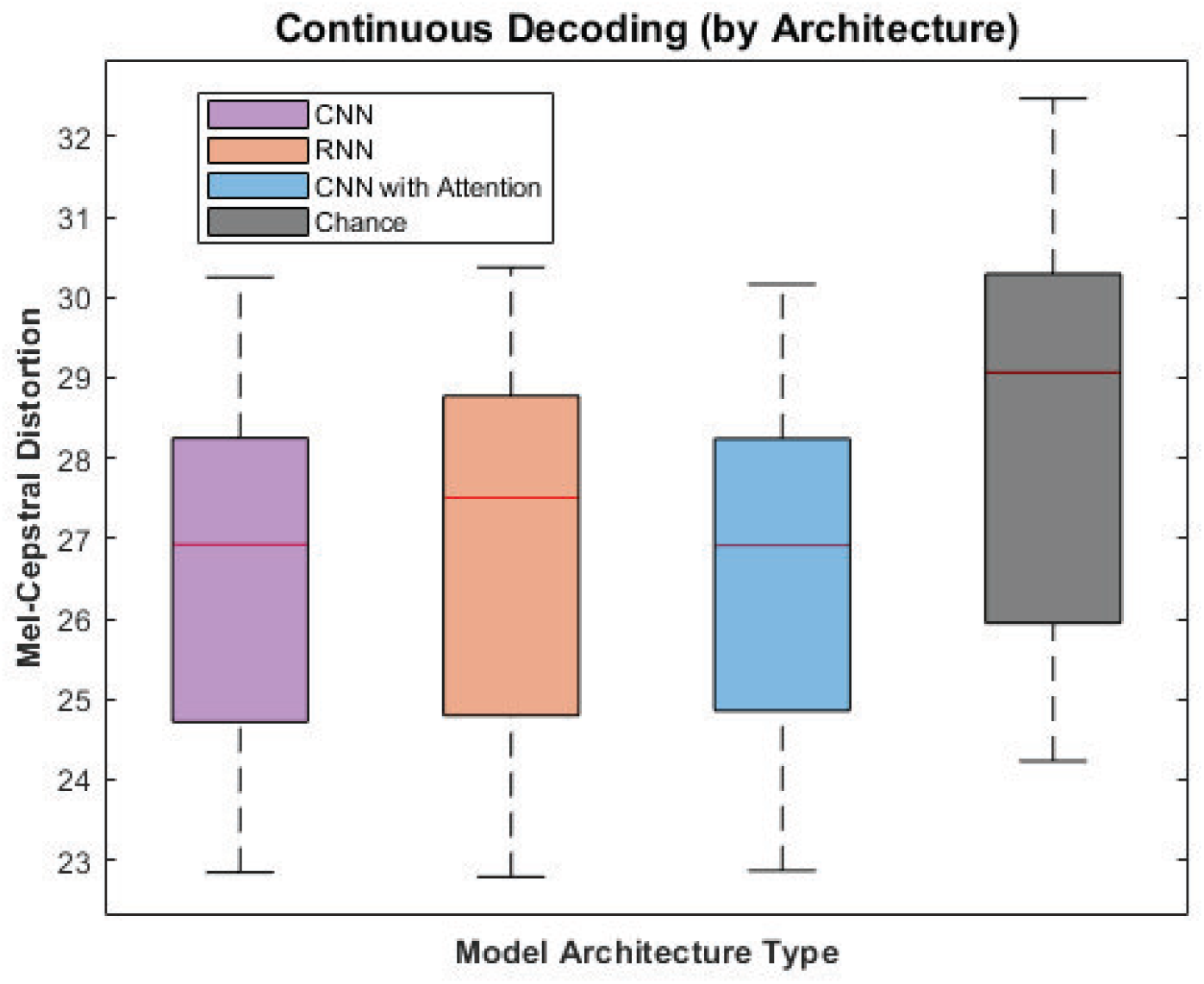
Continuous decoding performances by optimized architecture, averaged across cleaning methods. Note that lower MCD’s indicate better decoding performance.

When comparing the continuous prediction performance across cleaning methods (Figure 30), the ICA ICLabel method outperformed against models trained with the EEMD-CCA and EEMD-ICA cleaned datasets. Unlike the discrete decoding performances, ICA-ICLabel also outperformed against even the solely pre-processed data on average. While ICA-ICLabel reduced higher gamma activity to a lesser degree than EEMD-ICA and EEMD-CCA methods, it appears that decoding MFCCs benefits from some level of cleaning.

**Figure 30:**
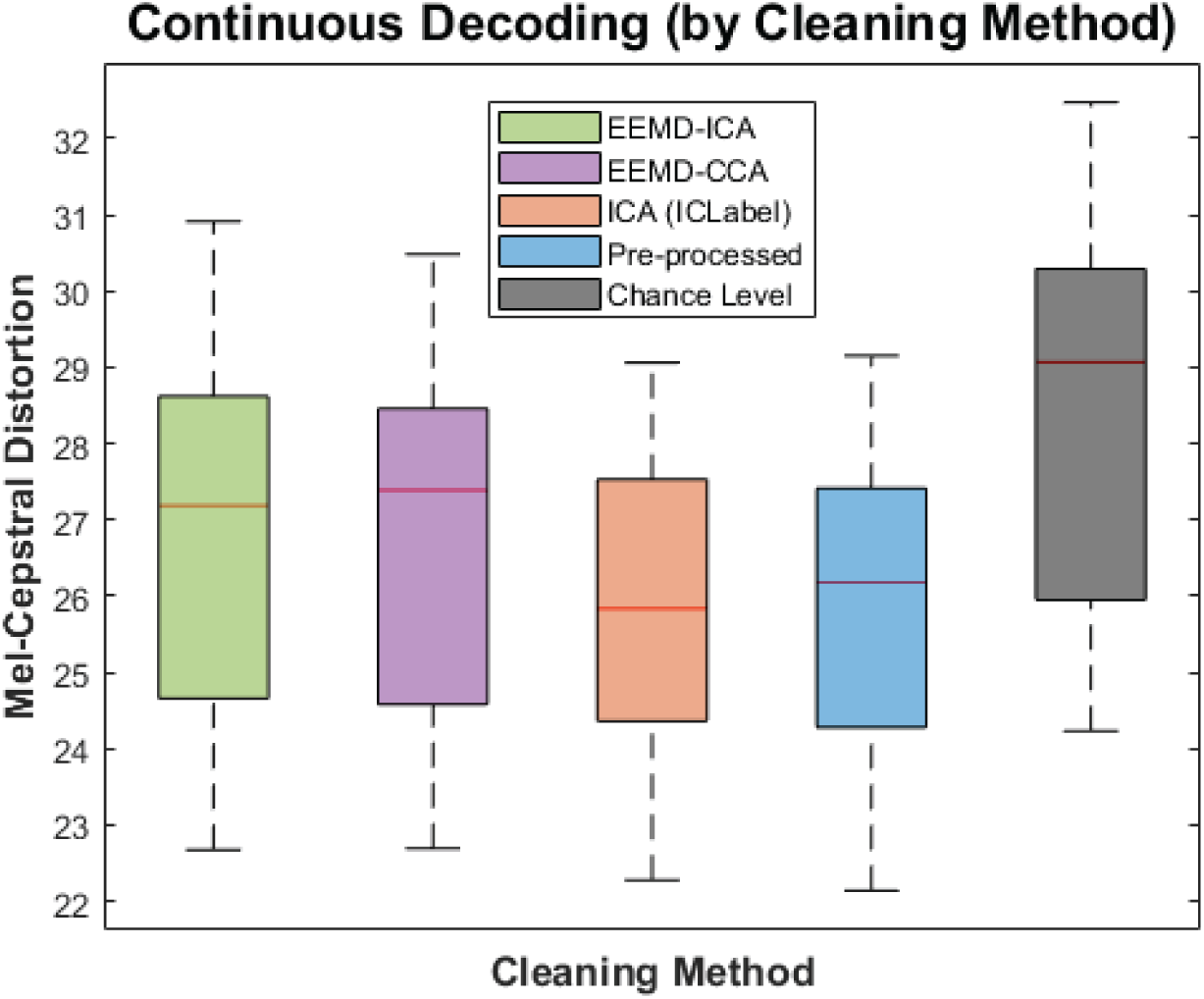
Continuous decoding performances by cleaning method, averaged across architecture. Note that lower MCD’s indicate better decoding performance.

Figure 31 presents an example reconstruction of the audio signal and resulting time-frequency plots derived from the True MFCCs (the first 25 MFCCs of the audio signal) and the Predicted MFCCs (as predicted from the continuous model), demonstrating clear temporal and frequency similarities between these two signals. The focus of this research was not on reconstructing intelligible audio signals, so advanced audio reconstruction models were not employed. Future research could find improvements in audio reconstruction by employing advanced reconstruction models, such as Wavenet [78] or LPCNet [79].

**Figure 31:**
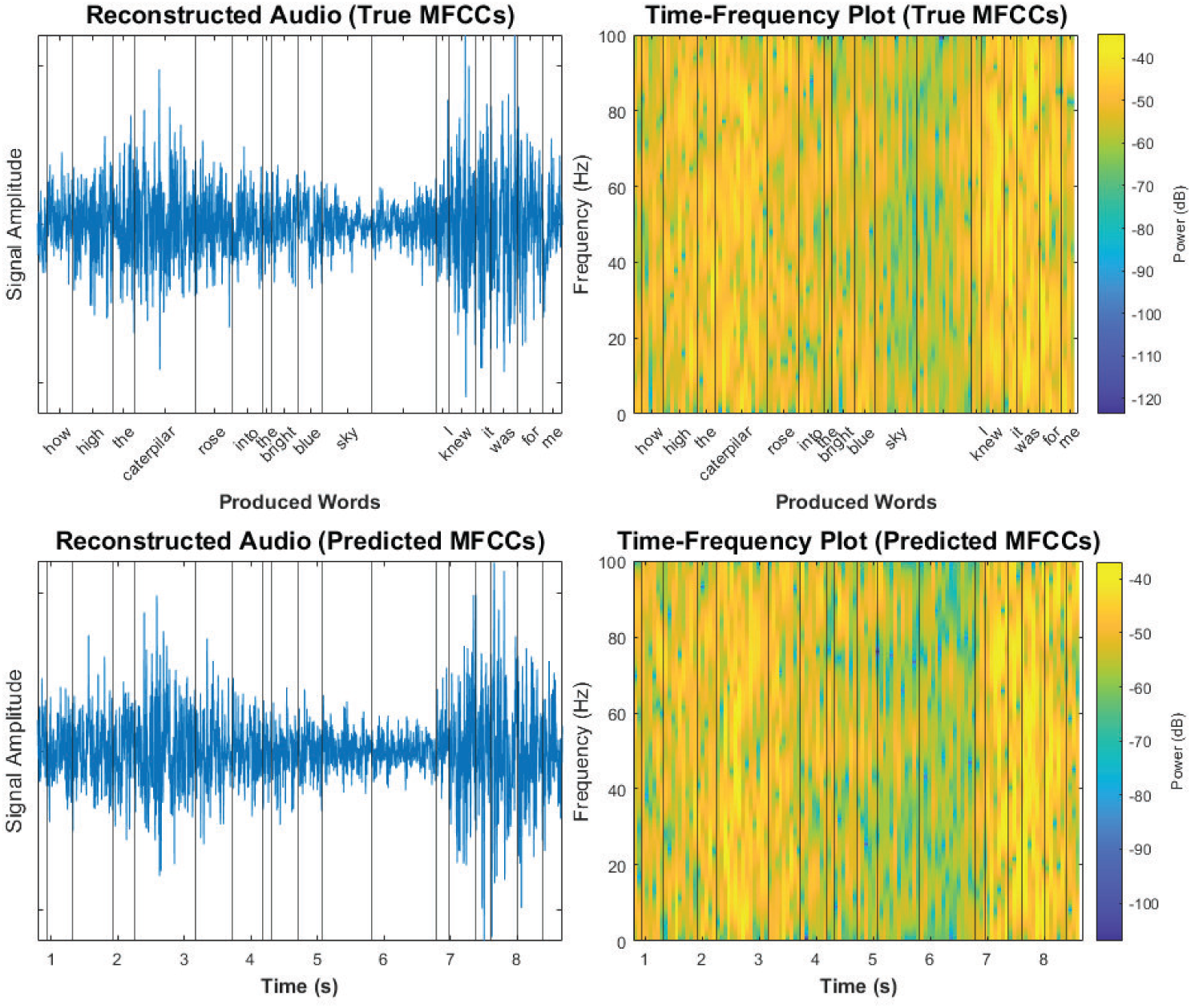
Example audio reconstruction and the resulting time-frequency plots as derived from the true and predicted MFCCs.

## 5. Discussion and Conclusion

In this research, Convolutional Neural Networks (CNN) and Recurrent Neural Networks (RNN), with and without the inclusion of an attention module, were optimized for the decoding of EEG signals in a speech BCI application, specifically for the decoding of discrete articulatory kinematic and phonetic classes, and continuous characteristics of the audio signal. Additionally, we assessed the channels and frequency bands of importance, explored different channel selection methods, and implemented Leave-One-Out (LOO) testing strategies. The results and limitations of our study are discussed below.

First, deep learning models were optimized by assessing multiple architectural modifications for CNN, RNN, and attention module architectures. Surprisingly, the CNN models outperformed the RNN models, despite RNNs being traditionally thought of as more appropriate for time-series data. Regarding the attention module, incorporating it with the optimized CNN model resulted in the best performance for all discrete decoding tasks. However, the improvement in performance was minimal compared to the optimized CNN model alone, and, considering the substantial increase in training time due to the attention module, its inclusion may not be a worthwhile addition.

For discrete decoding by class, models trained for all classes, except the Presence of Voice class, reliably achieved performance greater than chance. This suggests that specific phonemes or articulatory characteristics can be effectively differentiated for non-invasive speech BCI applications. While this decoding framework relied on knowing when the onset of a phoneme occurred, speech BCI applications can instead frame data collection by training participants to produce phoneme sequences at set intervals of time. With this phoneme sequence mechanism, a larger number of control signals could be obtained as compared to other traditional non-speech paradigms. For instance, phonemes /ae/, /ey/, and /s/ were highly distinguishable between each other. This would theoretically provide nine potential sequences of these phonemes, opening up speech-BCI enabled applications that involve up to nine classes. Alternatively, as these nine classes would not be actual words, future research could select a set of sentences, phrases, or words and incorporate MFA phoneme alignment into the processing pipeline to first isolate phonemes prior to decoding, which could potentially eliminate the need for unnatural timing in the above phoneme production method.

A perturbation analysis was implemented to assess the importance of different channels with respect to decoding performance. Interestingly, many of the discrete classes identified FP1 and FP2, electrodes closest to the face, as being important. This indicates that the deep learning models may still be focusing on channels that could be contaminated with EMG noise, even though several of the cleaning methods (EEMD-CCA, EEMD-ICA) removed up to 99% of the power in the upper gamma band. Training models when excluding these channels, surprisingly, led to improvements in performance, which was contrary to our initial hypothesis. This may be explained by the fact that the many facial muscles activated during speech are mixed as they propagate to the frontal EEG channels, making phoneme differentiation based on EMG contamination difficult. This improvement in performance could support the idea of simply removing channels with high EMG contamination, rather than relying on extensive EMG cleaning processes, which further opens up the possibility for real-time applicable EEG-based speech BCIs.

The analysis of frequency bands of importance revealed that Delta and Theta sub-bands were critical for discrete decoding, which aligns with existing literature on EEG frequency bands relevant to speech BCI [77, 76]. Additionally, the upper and lower Gamma bands were found to be important for decoding performance. Notably, when using EEMD-CCA cleaning, isolating the low gamma band resulted in significantly lower performance compared to other cleaning bands. However, isolating the upper gamma band still achieved high performance, despite the resulting low relative power in this sub-band for this cleaning method.

Different channel selection methods were assessed to better understand their impact on decoding performance for the possibility of channel-restricted real-time applications. While the use of all channels outperformed the assessed channel selection methods, models trained with reduced channel datasets still achieved performance above chance, indicating that channel-restricted applications can still achieve better than chance decoding performance. This may help to reduce the reliance on high-density EEG systems, improving prospects for clinical and commercial applications.

The LOO testing analysis was approached in four ways: training without any of the data from the participant of interest (POI), training with a single POI block used in the validation set, training with a single POI block in the training set and a single POI block in the validation set, and training with all data from the POI. For the three approaches that did not use all the data from the POI, models trained for most classes achieved performance greater than chance. In the final approach, where all the data from the POI was employed, virtually all discrete classes and cleaning methods experienced improvements to performance as compared to the participant-independent training. This implies that there are common neural correlates between participants and that data from other participants can be effectively used to improve decoding performance. As data from more participants are collected, this LOO testing framework could help to reduce the amount of data needed from each individual participant.

For the continuous decoding of acoustic characteristics, the resulting Continuous Mel-Cepstral Distortion (MCD) scores were significantly higher than invasive speech decoding MCDs, indicating worse overall decoding performance. However, all participants achieved MCD scores better than the chance model, indicating that there is promise in decoding continuous acoustic characteristics from EEG. Future research could improve upon these results by collecting longer periods of continuous speech production from a larger number of participants.

Several limitations were encountered in this study. Computational and time restrictions limited the selection of parameters for optimization. The time-intensive nature of continuous predictions, especially with the attention module, further compounded these limitations. Additionally, the lack of balance in the discrete decoding dataset hindered a more thorough analysis of the most discriminable set of phonemes. EEG data collection for this overt speech task proved to be energy-intensive for participants, so improving protocol efficiency by reducing inter-trial interval or including more words/sentences could facilitate improved decoding by allowing for more data collection. Additionally, the promising LOO results may indicate that the focus should be on collecting single sessions from more participants, rather than on collecting multiple sessions from individuals, as the inclusion of data from all participants led to consistent increases in performance as compared to the participant-independent models.

Future research could explore the use of vision transformers as an alternative to the 1-D multi-headed attention modules employed in this study. A focus would need to be given to modifying the attention window to avoid introducing artificial spatial relationships between channels. In conclusion, this research presented an optimization framework for EEG signal decoding in a speech BCI application. The performances of CNN, RNN, and attention models were evaluated, and the importance of channels and frequency bands was analyzed. Discrete decoding by class and continuous decoding of acoustic characteristics were performed, with model performances generally exceeding chance level. These research findings make a substantial contribution to the development of EEG-enabled speech synthesis frameworks. By effectively tackling the issue of EMG interference and employing optimized deep learning models for speech decoding, this study establishes a strong foundation for future advancements in the field of EEG-based speech brain-computer interfaces.

## 6. Acknowledgements

This research is supported in part by NSF IUCRC BRAIN award # 2137255, the University of Houston CLASS Early Career Research Grant, and the Research Computing Data Core at the University of Houston.

## 7. Author Contributions

A.C.: Conceptualization, Methodology, Software, Validation, Formal analysis, Investigation, Data Curation, Writing - Original Draft, Visualization; H.D.: Resources, Investigation, Writing - Review & Editing, Project administration, Funding acquisition; J.L.C.V.: Resources, Writing - Review & Editing, Project administration, Supervision, Funding acquisition

## 8. References

[1] N. Team, “Ipums nhis online data analysis system.” [Online]. Available: https://nhis.ipums.org/nhis/sda.shtml

[2] Q. Rabbani, G. Milsap, and N. E. Crone, “The potential for a speech brain–computer interface using chronic electrocorticography,” Neurotherapeutics, vol. 16, no. 1, pp. 144–165, 2019.

[3] G. K. Anumanchipalli, J. Chartier, and E. F. Chang, “Speech synthesis from neural decoding of spoken sentences,” Nature, vol. 568, no. 7753, pp. 493–498, 2019.

[4] J. A. Perge, M. L. Homer, W. Q. Malik, S. Cash, E. Eskandar, G. Friehs, J. P. Donoghue, and L. R. Hochberg, “Intra-day signal instabilities affect decoding performance in an intracortical neural interface system,” Journal of Neural Engineering, vol. 10, no. 3, p. 036004, 2013.

[5] X. Chen, X. Xu, A. Liu, S. Lee, X. Chen, X. Zhang, M. J. McKeown, and Z. J. Wang, “Removal of muscle artifacts from the eeg: a review and recommendations,” IEEE Sensors Journal, vol. 19, no. 14, pp. 5353–5368, 2019.

[6] C. Herff and T. Schultz, “Automatic speech recognition from neural signals: a focused review,” Frontiers in Neuroscience, vol. 10, p. 429, 2016.

[7] S. Chakrabarti, H. M. Sandberg, J. S. Brumberg, and D. J. Krusienski, “Progress in speech decoding from the electrocorticogram,” Biomedical Engineering Letters, vol. 5, no. 1, pp. 10–21, 2015.

[8] S. Martin, I. Iturrate, J. d. R. Millán, R. T. Knight, and B. N. Pasley, “Decoding inner speech using electrocorticography: Progress and challenges toward a speech prosthesis,” Frontiers in Neuroscience, vol. 12, p. 422, 2018.

[9] O. Woolnough, K. J. Forseth, P. S. Rollo, and N. Tandon, “Uncovering the functional anatomy of the human insula during speech,” Elife, vol. 8, p. e53086, 2019.

[10] G. Krishna, C. Tran, J. Yu, and A. H. Tewfik, “Speech recognition with no speech or with noisy speech,” in *ICASSP 2019-2019 IEEE International Conference on Acoustics, Speech and Signal Processing (ICASSP)*. IEEE, 2019, pp. 1090–1094.

[11] G. Krishna, C. Tran, Y. Han, M. Carnahan, and A. H. Tewfik, “Speech synthesis using eeg,” in *ICASSP 2020-2020 IEEE International Conference on Acoustics, Speech and Signal Processing (ICASSP)*. IEEE, 2020, pp. 1235–1238.

[12] J. Willis, A. Nelson, J. Rice, and F. W. Black, “The topography of muscle activity in quantitative eeg,” Clinical Electroencephalography, vol. 24, no. 3, pp. 123–126, 1993.

[13] J. Gotman, J. Ives, and P. Gloor, “Frequency content of eeg and emg at seizure onset: possibility of removal of emg artefact by digital filtering,” Electroencephalography and Clinical Neurophysiology, vol. 52, no. 6, pp. 626–639, 1981.

[14] R. J. Davidson, “Eeg measures of cerebral asymmetry: Conceptual and methodological issues,” International Journal of Neuroscience, vol. 39, no. 1-2, pp. 71–89, 1988.

[15] B. H. Friedman and J. F. Thayer, “Facial muscle activity and eeg recordings: redundancy analysis,” Electroencephalography and Clinical Neurophysiology, vol. 79, no. 5, pp. 358–360, 1991.

[16] S. Waldert, “Invasive vs. non-invasive neuronal signals for brain-machine interfaces: will one prevail?” Frontiers in Neuroscience, vol. 10, p. 295, 2016.

[17] A. Craik, Y. He, and J. L. Contreras-Vidal, “Deep learning for electroencephalogram (eeg) classification tasks: a review,” Journal of Neural Engineering, vol. 16, no. 3, p. 031001, 2019.

[18] N. Yoshimura, A. Nishimoto, A. N. Belkacem, D. Shin, H. Kambara, T. Hanakawa, and Y. Koike, “Decoding of covert vowel articulation using electroencephalography cortical currents,” Frontiers in Neuroscience, vol. 10, p. 175, 2016.

[19] R. D. Kent, “Research on speech motor control and its disorders: A review and prospective,” Journal of Communication Disorders, vol. 33, no. 5, pp. 391–428, 2000.

[20] S. P. Fitzgibbon, D. M. Powers, K. J. Pope, and C. R. Clark, “Removal of eeg noise and artifact using blind source separation,” Journal of Clinical Neurophysiology, vol. 24, no. 3, pp. 232–243, 2007.

[21] C. J. James and C. W. Hesse, “Independent component analysis for biomedical signals,” Physiological Measurement, vol. 26, no. 1, p. R15, 2004.

[22] L. Sun, Y. Liu, and P. J. Beadle, “Independent component analysis of eeg signals,” in *Proceedings of 2005 IEEE International Workshop on VLSI Design and Video Technology*, *2005*. IEEE, 2005, pp. 219–222.

[23] M. Scherg and J. Ebersole, “Models of brain sources,” Brain topography, vol. 5, pp. 419–423, 1993.

[24] K. Nazarpour, A. H. Al-Timemy, G. Bugmann, and A. Jackson, “A note on the probability distribution function of the surface electromyogram signal,” Brain Research Bulletin, vol. 90, pp. 88–91, 2013.

[25] M. B. I. Reaz, M. S. Hussain, and F. Mohd-Yasin, “Techniques of emg signal analysis: detection, processing, classification and applications,” Biological Procedures Online, vol. 8, pp. 11–35, 2006.

[26] Z. Wu and N. E. Huang, “Ensemble empirical mode decomposition: a noise-assisted data analysis method,” Advances in Adaptive Data Analysis, vol. 1, no. 01, pp. 1–41, 2009.

[27] G. Rilling, P. Flandrin, and P. Goncalves, “On empirical mode decomposition and its algorithms,” in *IEEE-EURASIP Workshop on Nonlinear Signal and Image Processing*, vol. 3, no. 3. Grado: IEEE, 2003, pp. 8–11.

[28] D. Moher, A. Liberati, J. Tetzlaff, D. G. Altman, and P. Group*, “Preferred reporting items for systematic reviews and meta-analyses: the prisma statement,” Annals of Internal Medicine, vol. 151, no. 4, pp. 264–269, 2009.

[29] S. Zhao and F. Rudzicz, “Classifying phonological categories in imagined and articulated speech,” in 2015 *IEEE International Conference on Acoustics, Speech and Signal Processing (ICASSP)*. IEEE, 2015, pp. 992–996.

[30] N. Nieto, V. Peterson, H. L. Rufiner, J. E. Kamienkowski, and R. Spies, “Thinking out loud, an open-access eeg-based bci dataset for inner speech recognition,” Scientific Data, vol. 9, no. 1, pp. 1–17, 2022.

[31] Y.-E. Lee and S.-H. Lee, “Eeg-transformer: Self-attention from transformer architecture for decoding eeg of imagined speech,” in 2022 *10th International Winter Conference on Brain-Computer Interface (BCI)*. IEEE, 2022, pp. 1–4.

[32] G. Krishna, C. Tran, M. Carnahan, Y. Han, and A. H. Tewfik, “Improving eeg based continuous speech recognition,” arXiv preprint arXiv*:1911.11610*, 2019.

[33] G. Krishna, C. Tran, M. Carnahan, and A. Tewfik, “Advancing speech recognition with no speech or with noisy speech,” in 2019 *27th European Signal Processing Conference (EUSIPCO)*. IEEE, 2019, pp. 1–5.

[34] G. Krishna, C. Tran, M. Carnahan, and A. H. Tewfik, “Eeg based continuous speech recognition using transformers,” arXiv preprint arXiv:2001.00501, 2019.

[35] B. Products, “Actichamp series: Brain products gmbh gt; solutions,” May 2023. [Online]. Available: https://www.brainproducts.com/solutions/actichamp/

[36] M. Zhu, H. Zhang, X. Wang, X. Wang, Z. Yang, C. Wang, O. W. Samuel, S. Chen, and G. Li, “Towards optimizing electrode configurations for silent speech recognition based on high-density surface electromyography,” Journal of Neural Engineering, vol. 18, no. 1, p. 016005, 2021.

[37] O. Abbasi, N. Steingräber, and J. Gross, “Correcting meg artifacts caused by overt speech,” Frontiers in Neuroscience, vol. 15, p. 691, 2021.

[38] WhisperRoom, Inc., “Sound isolation enclosures,” April 2024. [Online]. Available: https://whisperroom.com/

[39] D. H. Brainard and S. Vision, “The psychophysics toolbox,” Spatial Vision, vol. 10, no. 4, pp. 433–436, 1997.

[40] A. Wrench, “A multichannel articulatory speech database and its application for automatic speech recognition,” in Proc. 5th Seminar on Speech Rroduction: Models and Data, 2000, 2000.

[41] A. C. Lammert, J. Melot, D. E. Sturim, D. J. Hannon, R. DeLaura, J. R. Williamson, G. Ciccarelli, and T. F. Quatieri, “Analysis of phonetic balance in standard english passages,” Journal of Speech, Language, and Hearing Research, vol. 63, no. 4, pp. 917–930, 2020.

[42] M. McAuliffe, M. Socolof, S. Mihuc, M. Wagner, and M. Sonderegger, “Montreal forced aligner: Trainable text-speech alignment using kaldi.” in Interspeech, vol. 2017, 2017, pp. 498–502.

[43] L. Rabiner and B.-H. Juang, Fundamentals of speech recognition. Prentice-Hall, Inc., 1993.

[44] SCCN, “Makoto’s preprocessing pipeline,” n.d. [Online]. Available: https://sccn.ucsd.edu/wiki/Makoto’spreprocessingpipeline

[45] A. de Cheveigné, “Zapline: A simple and effective method to remove power line artifacts,” NeuroImage, vol. 207, p. 116356, 2020.

[46] A. Kilicarslan and J. L. Contreras-Vidal, “Towards a unified framework for de-noising neural signals,” in 2019 *41st Annual International Conference of the IEEE Engineering in Medicine and Biology Society (EMBC)*. IEEE, 2019, pp. 620–623.

[47] Á. Martínez-Ballester, M. Ortiz, E. Iáñez, and J. M. Azorín, “Optimización de parámetros para un algoritmo de eliminación de artefactos oculares,” LOS CONFERENCIANTES, p. 18, 2021.

[48] C.-Y. Chang, S.-H. Hsu, L. Pion-Tonachini, and T.-P. Jung, “Evaluation of artifact subspace reconstruction for automatic eeg artifact removal,” in 2018 *40th Annual International Conference of the IEEE Engineering in Medicine and Biology Society (EMBC)*. IEEE, 2018, pp. 1242–1245.

[49] X. Chen, Q. Liu, W. Tao, L. Li, S. Lee, A. Liu, Q. Chen, J. Cheng, M. J. McKeown, and Z. J. Wang, “Remae: User-friendly toolbox for removing muscle artifacts from eeg,” IEEE Transactions on Instrumentation and Measurement, vol. 69, no. 5, pp. 2105–2119, 2019.

[50] T.-W. Lee, M. Girolami, and T. J. Sejnowski, “Independent component analysis using an extended infomax algorithm for mixed subgaussian and supergaussian sources,” Neural Computation, vol. 11, no. 2, pp. 417–441, 1999.

[51] A. C. Tang, M. T. Sutherland, and C. J. McKinney, “Validation of sobi components from high-density eeg,” NeuroImage, vol. 25, no. 2, pp. 539–553, 2005.

[52] G. Sahonero-Alvarez and H. Calderon, “A comparison of sobi, fastica, jade and infomax algorithms,” in Proceedings of the 8th international multi-conference on complexity, informatics and cybernetics, 2017, pp. 17–22.

[53] K. T. Sweeney, S. F. McLoone, and T. E. Ward, “The use of ensemble empirical mode decomposition with canonical correlation analysis as a novel artifact removal technique,” IEEE transactions on Biomedical Engineering, vol. 60, no. 1, pp. 97–105, 2012.

[54] X. Chen, Q. Chen, Y. Zhang, and Z. J. Wang, “A novel eemd-cca approach to removing muscle artifacts for pervasive eeg,” IEEE Sensors Journal, vol. 19, no. 19, pp. 8420–8431, 2018.

[55] D. M. Vos, S. Riès, K. Vanderperren, B. Vanrumste, F.-X. Alario, V. S. Huffel, and B. Burle, “Removal of muscle artifacts from eeg recordings of spoken language production,” Neuroinformatics, vol. 8, no. 2, pp. 135–150, 2010.

[56] Q. Liu, A. Liu, X. Zhang, X. Chen, R. Qian, and X. Chen, “Removal of emg artifacts from multichannel eeg signals using combined singular spectrum analysis and canonical correlation analysis,” Journal of Healthcare Engineering, vol. 2019, 2019.

[57] A. Likas, N. Vlassis, and J. J. Verbeek, “The global k-means clustering algorithm,” Pattern Recognition, vol. 36, no. 2, pp. 451–461, 2003.

[58] N. Chomsky and M. Halle, “The sound pattern of english.” 1968.

[59] C. P. Browman and L. Goldstein, “Articulatory phonology: An overview,” Phonetica, vol. 49, no. 3-4, pp. 155–180, 1992.

[60] L. E. Rolston, “An independent assessment of phonetic distinctive feature sets used to model pronunciation variation,” Ph.D. dissertation, 2014.

[61] V. Peter, G. McArthur, and S. Crain, “Using event-related potentials to measure phrase boundary perception in english,” BMC neuroscience, vol. 15, no. 1, pp. 1–11, 2014.

[62] K. O’Shea and R. Nash, “An introduction to convolutional neural networks,” arXiv preprint arXiv:1511.08458, 2015.

[63] S. Santurkar, D. Tsipras, A. Ilyas, and A. Madry, “How does batch normalization help optimization?” Advances in Neural Information Processing Systems, vol. 31, 2018.

[64] P. Baldi and P. J. Sadowski, “Understanding dropout,” Advances in Neural Information Processing Systems, vol. 26, 2013.

[65] V. J. Lawhern, A. J. Solon, N. R. Waytowich, S. M. Gordon, C. P. Hung, and B. J. Lance, “Eegnet: a compact convolutional neural network for eeg-based brain–computer interfaces,” Journal of Neural Engineering, vol. 15, no. 5, p. 056013, 2018.

[66] L. R. Medsker and L. Jain, “Recurrent neural networks,” Design and Applications, vol. 5, no. 64-67, p. 2, 2001.

[67] A. Sherstinsky, “Fundamentals of recurrent neural network (rnn) and long short-term memory (lstm) network,” Physica D: Nonlinear Phenomena, vol. 404, p. 132306, 2020.

[68] R. Dey and F. M. Salem, “Gate-variants of gated recurrent unit (gru) neural networks,” in 2017 *IEEE 60th International Midwest Symposium on Circuits and Systems (MWSCAS)*. IEEE, 2017, pp. 1597–1600.

[69] A. Vaswani, N. Shazeer, N. Parmar, J. Uszkoreit, L. Jones, A. N. Gomez, L . Kaiser, and I. Polosukhin, “Attention is all you need,” Advances in Neural Information Processing Systems, vol. 30, 2017.

[70] J. T. Panachakel and A. G. Ramakrishnan, “Decoding covert speech from eeg-a comprehensive review,” Frontiers in Neuroscience, vol. 15, p. 392, 2021.

[71] A. Gramfort, M. Luessi, E. Larson, D. A. Engemann, D. Strohmeier, C. Brodbeck, R. Goj, M. Jas, T. Brooks, L. Parkkonen, and M. Hämäläinen, “Meg and eeg data analysis with mne-python,” Frontiers in Neuroscience, p. 267, 2013.

[72] M. R. Hasan, M. Jamil, and M. Rahman, “Speaker identification using mel frequency cepstral coefficients,” Variations, vol. 1, no. 4, pp. 565–568, 2004.

[73] R. Kubichek, “Mel-cepstral distance measure for objective speech quality assessment,” in Proceedings of IEEE Pacific Rim Conference on Communications Computers and Signal processing, vol. 1. IEEE, 1993, pp. 125–128.

[74] M. Abadi, A. Agarwal, P. Barham, E. Brevdo, Z. Chen, C. Citro, G. S. Corrado, A. Davis, J. Dean, M. Devin, S. Ghemawat, I. Goodfellow, A. Harp, G. Irving, M. Isard, Y. Jia, R. Jozefowicz, L. Kaiser, M. Kudlur, J. Levenberg, D. Mané, R. Monga, S. Moore, D. Murray, C. Olah, M. Schuster, J. Shlens, B. Steiner, I. Sutskever, K. Talwar, P. Tucker, V. Vanhoucke, V. Vasudevan, F. Viégas, O. Vinyals, P. Warden, M. Wattenberg, M. Wicke, Y. Yu, and X. Zheng, “TensorFlow: Large-scale machine learning on heterogeneous systems,” 2015, software available from tensorflow.org. [Online]. Available: https://www.tensorflow.org/

[75] F. Pedregosa, G. Varoquaux, A. Gramfort, V. Michel, B. Thirion, O. Grisel, M. Blondel, P. Prettenhofer, R. Weiss, V. Dubourg, J. Vanderplas, A. Passos, D. Cournapeau, M. Brucher, M. Perrot, and E. Duchesnay, “Scikit-learn: Machine learning in Python,” Journal of Machine Learning Research, vol. 12, pp. 2825–2830, 2011.

[76] F. Bröhl and C. Kayser, “Delta/theta band eeg differentially tracks low and high frequency speech-derived envelopes,” Neuroimage, vol. 233, p. 117958, 2021.

[77] E. S. Teoh, M. S. Cappelloni, and E. C. Lalor, “Prosodic pitch processing is represented in delta-band eeg and is dissociable from the cortical tracking of other acoustic and phonetic features,” European Journal of Neuroscience, vol. 50, no. 11, pp. 3831–3842, 2019.

[78] A. v. d. Oord, S. Dieleman, H. Zen, K. Simonyan, O. Vinyals, A. Graves, N. Kalchbrenner, A. Senior, and K. Kavukcuoglu, “Wavenet: A generative model for raw audio,” arXiv preprint arXiv:1609.03499, 2016.

[79] J.-M. Valin and J. Skoglund, “Lpcnet: Improving neural speech synthesis through linear prediction,” in *ICASSP 2019-2019 IEEE International Conference on Acoustics, Speech and Signal Processing (ICASSP)*. IEEE, 2019, pp. 5891–5895.

[80] G. Krishna, C. Tran, M. Carnahan, and A. H. Tewfik, “Advancing speech synthesis using eeg,” in 2021 *10th International IEEE/EMBS Conference on Neural Engineering (NER)*. IEEE, 2021, pp. 199–204.

[81] G. Krishna, Y. Han, C. Tran, M. Carnahan, and A. H. Tewfik, “State-of-the-art speech recognition using eeg and towards decoding of speech spectrum from eeg,” arXiv preprint arXiv:1908.05743, 2019.

[82] S.-H. Lee, M. Lee, and S.-W. Lee, “Eeg representations of spatial and temporal features in imagined speech and overt speech,” in Asian Conference on Pattern Recognition. Springer, 2019, pp. 387–400.

[83] S.-H. Lee, Y.-E. Lee, and S.-W. Lee, “Voice of your brain: Cognitive representations of imagined speech, overt speech, and speech perception based on eeg,” arXiv preprint arXiv:2105.14787, 2021.

[84] D.-H. Lee, S.-J. Kim, and S.-W. Lee, “Dal: Feature learning from overt speech to decode imagined speech-based eeg signals with convolutional autoencoder,” arXiv preprint arXiv:2107.07064, 2021.

[85] B. McMurray, M. E. Sarrett, S. Chiu, A. K. Black, A. Wang, R. Canale, and R. N. Aslin, “Decoding the temporal dynamics of spoken word and nonword processing from eeg,” NeuroImage, p. 119457, 2022.

[86] N. Janssen, M. v. d. Meij, P. J. López-Pérez, and H. A. Barber, “Exploring the temporal dynamics of speech production with eeg and group ica,” Scientific Reports, vol. 10, no. 1, pp. 1–14, 2020.

[87] L. Pion-Tonachini, K. Kreutz-Delgado, and S. Makeig, “Iclabel: An automated electroencephalographic independent component classifier, dataset, and website,” NeuroImage, vol. 198, pp. 181–197, 2019.

[88] R. Tibshirani, G. Walther, and T. Hastie, “Estimating the number of clusters in a data set via the gap statistic,” Journal of the Royal Statistical Society: Series B (Statistical Methodology), vol. 63, no. 2, pp. 411–423, 2001.

[89] T. Caliński and J. Harabasz, “A dendrite method for cluster analysis,” Communications in Statistics-theory and Methods, vol. 3, no. 1, pp. 1–27, 1974.

[90] M. Scherg, “Fundamentals of dipole source potential analysis,” Advances in Audiology, vol. 6, no. 40-69, p. 25, 1990.

[91] C. M. Michel and B. He, “Eeg source localization,” Handbook of Clinical Neurology, vol. 160, pp. 85–101, 2019.

[92] M. Strotzer, “One century of brain mapping using brodmann areas,” Clinical Neuroradiology, vol. 19, no. 3, p. 179, 2009.

[93] G. Toyoda, E. C. Brown, N. Matsuzaki, K. Kojima, M. Nishida, and E. Asano, “Electrocorticographic correlates of overt articulation of 44 english phonemes: intracranial recording in children with focal epilepsy,” Clinical Neurophysiology, vol. 125, no. 6, pp. 1129– 1137, 2014.

[94] R. Pijnenburg, L. H. Scholtens, D. J. Ardesch, S. C. de Lange, Y. Wei, and M. P. van den Heuvel, “Myelo-and cytoarchitectonic microstructural and functional human cortical atlases reconstructed in common mri space,” NeuroImage, vol. 239, p. 118274, 2021.

[95] A. Ardila, B. Bernal, and M. Rosselli, “How localized are language brain areas? a review of brodmann areas involvement in oral language,” Archives of Clinical Neuropsychology, vol. 31, no. 1, pp. 112–122, 2016.

[96] I. Cohen, Y. Huang, J. Chen, J. Benesty, J. Benesty, J. Chen, Y. Huang, and I. Cohen, “Pearson correlation coefficient,” Noise Reduction in Speech Processing, pp. 1–4, 2009.

