## Appendices for "Continuous and discrete decoding of overt speech with electroencephalography"

### 9. Appendix

#### *9.1. Phoneme and class sample counts*

Table 5: Resulting phoneme counts from 1, 4, and 8 blocks of collection.

| Phoneme | 1 Block | 4 Blocks | 8 Blocks |
|---|---|---|---|
| AA | 36 | 144 | 288 |
| AE | 61 | 244 | 488 |
| AH | 236 | 944 | 1888 |
| AO | 32 | 128 | 256 |
| AW | 10 | 40 | 80 |
| AY | 61 | 244 | 488 |
| B | 54 | 216 | 432 |
| CH | 9 | 36 | 72 |
| D | 87 | 348 | 696 |
| DH | 2 | 8 | 16 |
| EH | 69 | 276 | 552 |
| ER | 59 | 236 | 472 |
| EY | 37 | 148 | 296 |
| F | 46 | 184 | 368 |
| G | 17 | 68 | 136 |
| HH | 27 | 108 | 216 |
| IH | 138 | 552 | 1104 |
| IY | 69 | 276 | 552 |
| JH | 9 | 36 | 72 |
| K | 87 | 348 | 696 |

Table 6: Resulting phoneme counts from 1, 4, and 8 blocks of collection.

| Phoneme | 1 Block | 4 Blocks | 8 Blocks |
|---|---|---|---|
| L | 96 | 384 | 768 |
| M | 71 | 284 | 568 |
| N | 158 | 632 | 1264 |
| NG | 21 | 84 | 168 |
| OW | 47 | 188 | 376 |
| OY | 5 | 20 | 40 |
| P | 51 | 204 | 408 |
| R | 109 | 436 | 872 |
| S | 103 | 412 | 824 |
| SH | 11 | 44 | 88 |
| T | 149 | 596 | 1192 |
| TH | 17 | 68 | 136 |
| UH | 8 | 32 | 64 |
| UW | 24 | 96 | 192 |
| V | 43 | 172 | 344 |
| W | 53 | 212 | 424 |
| Y | 15 | 60 | 120 |
| Z | 84 | 336 | 672 |
| ZH | 18 | 72 | 144 |

#### *9.2. Overt Speech Decoding Literature Review*

Dictionary: Partic.: Participants, Chan.: Channels, Aud.: Audio availability, sent.: sentence, mus.: music, V.: Vowels, Ph.: Phonemes, ICA: Independent Component Analysis, RNN: Recurrent Neural Network, Tran.: Transformer, GRU: Gated Recurrent Unit, art.: Articulatory stage, LSTM: Long Short-Term Memory, LDA: Linear

Table 7: Discrete class sample counts for 1, 4, and 8 blocks of collection.

| Group Name | Class | 1 Block | 4 Blocks | 8 Blocks |
|---|---|---|---|---|
| **Vowel vs Consonant** | **Vowel Counts** | 887 | 3548 | 7096 |
| | **Consonant Counts** | 1426 | 5704 | 11408 |
| **Manner of Articulation (Consonants)** | **Affricate** | 22 | 88 | 176 |
| | **Approximant** | 284 | 1136 | 2272 |
| | **Fricative** | 391 | 1564 | 3128 |
| | **Nasal** | 252 | 1008 | 2016 |
| | **Unvoiced Stop** | 286 | 1144 | 2288 |
| | **Voiced Stop** | 157 | 628 | 1256 |
| **Place of Articulation (Consonants)** | **Alveolar** | 427 | 1708 | 3416 |
| | **Dental** | 94 | 376 | 752 |
| | **Labial (Bi)** | 175 | 700 | 1400 |
| | **Labial (Dental)** | 93 | 372 | 744 |
| | **Palatal** | 46 | 184 | 368 |
| | **Velar** | 123 | 492 | 984 |
| **Place of Articulation (Vowels)** | **Front** | 314 | 1256 | 2512 |
| | **Central** | 277 | 1108 | 2216 |
| | **Back** | 132 | 528 | 1056 |
| | **Dipthong** | 164 | 656 | 1312 |
| **Presence of Voice** | **Voiced** | 1175 | 4700 | 9400 |
| | **Voiced (No vowels)** | 288 | 1152 | 2304 |
| | **Unvoiced** | 439 | 1756 | 3512 |

Table 8: Literature review of research involving overt speech decoding.

| Research | Task | Partic. | Chan. | Aud. | EMG | Model | Output |
|---|---|---|---|---|---|---|---|
| [33] | A: 30 sent.<br>B: A + mus.) | A: 10<br>B: 8 | 32 | Yes | ICA | RNN | Words |
| [34] | A: 30 sent.<br>B: A + mus.) | A: 10<br>B: 8 | 32 | Yes | ICA | Tran. | Words |
| [32] | A: 30 sent.<br>B: A + mus.) | A: 10<br>B: 8 | 32 | Yes | ICA | RNN (GRU) | Words |
| [10] | 5 V.,<br>4 words | 4 | 32 | Yes | ICA | RNN (GRU) | Words/V. |
| [80] | 4 sent. | 4 | 32 | Yes | ICA | RNN (w/wo art.) | MFCCs |
| [11] | 4 sent. | 4 | 32 | Yes | ICA | RNN (GRU) | MFCCs |
| [81] | 4 sent. | 4 | 32 | Yes | ICA | RNN (LSTM) | MFCCs |
| [82] | 12 words | 9 | 64 | No | None | LDA | Words |
| [31] | 12 words | 9 | 64 | No | ICA | Tran. | Words |
| [83] | 12 words | 9 | 64 | No | None | CNN | Words |
| [29] | 7 Ph./4 words | 14 | 64 | Yes | None | DBN | Words/Ph. |
| [84] | 4 words | 8 | 58 | No | None | N/A* | Words |
| [85] | 8 word pairs | 16 | 128 | Yes | None | N/A** | N/A |
| [86] | 100 pictures | 30 | 32 | No | ICA | N/A*** | N/A |
| [30] | 4 words | 10 | 128 | No | ICA | Data Only | N/A |

Discriminant Analysis, DBN: Deep Belief Network, MFCCs: Mel-Frequency Cepstral Coefficients
* Focused on Overt / Covert correlation
** Focused on Perception / Production correlation
*** Focused on ICA Group Analysis

##### *9.3. Passages for overt production*

*9.3.1. Extended grandfather passage* You wish to know all about my grandfather. Well, he is nearly ninety three years old, yet he still thinks as swiftly as ever. He dresses himself in an old black frock coat with a thick hood, usually several buttons missing. A long beard clings to his chin, giving those who observe him a pronounced feeling of the utmost respect. When he speaks, his voice is just a bit cracked and quivers a bit, but it pulls the listener in. Twice each day he plays skillfully and with zest upon a small organ. Except in the winter when the snow or ice prevents, he slowly takes a short walk in the open air each day. We have often urged him to walk more and smoke less, but he always answers, Banana oil! Grandfather likes to be modern in his language.

*9.3.2. Rainbow passage* When the sunlight strikes raindrops in the air, they act as a prism and form a rainbow. The rainbow is a division of white light into many beautiful colors. These take the shape of a long round arch, with its path high above, and its two ends apparently beyond the horizon. There is , according to legend, a boiling pot of gold at one end. People look, but no one ever finds it. When a man looks for something beyond his reach, his friends say he is looking for the pot of gold at the end of the rainbow. Throughout the centuries people have explained the rainbow in various ways. Some have accepted it as a miracle without physical explanation. To the Hebrews it was a token that there would be no more universal floods. The Greeks used to imagine that it was a sign from the gods to foretell war or heavy rain. The Norsemen considered the rainbow as a bridge over which the gods passed from earth to their home in the sky. Others have tried to explain the phenomenon physically. Aristotle thought that the rainbow was caused by reflection of the sun's rays by the rain. Since then physicists have found that it is not reflection, but refraction by the raindrops which causes the rainbows. Many complicated ideas about the rainbow have been formed. The difference in the rainbow depends considerably upon the size of the drops, and the width of the colored band increases as the size of the drops increases. The actual primary rainbow observed is said to be the effect of super- imposition of a number of bows. If the red of the second bow falls upon the green of the first, the result is to give a bow with an abnormally wide yellow band, since red and green light when mixed form yellow. This is a very common type of bow, one showing mainly red and yellow, with little or no green or blue.

*9.3.3. Caterpillar passage* Do you like amusement parks? Well, I sure do. To amuse myself, I went twice last spring. My most memorable moment was riding on the Caterpillar, which is a gigantic roller coaster high above the ground. When I saw how high the Caterpillar rose into the bright blue sky I knew it was for me. After waiting in line for thirty minutes, I made it to the front where the man measured my height to see if I was tall enough. I gave the man my coins, asked for change, and jumped on the cart. Tick, tick, tick, the Caterpillar climbed slowly up the tracks. It went SO high I could see the parking lot. Boy was I scared! I thought to myself, "There's no turning back now." People were so scared they screamed as we swiftly zoomed fast, fast, and faster along the tracks. As quickly as it started, the Caterpillar came to a stop. Unfortunately, it was time to pack the car and drive home. That night I dreamt of the wild ride on the Caterpillar. Taking a trip to the amusement park and riding on the

Caterpillar was my most memorable moment ever!

##### *9.4. Power line noise removal*

To remove power line noise, EEGLab's Zapline function [45] was employed. Initially, this was the only employed power line noise removal method, but an analysis of the resulting Power Spectral Density (PSD) plots revealed the continued contamination of the EEG data with power line noise (Figure 32). An investigation found that Zapline was not effective in removing all the power line noise from the passive EOG sensors. Since the EOG sensors were used solely for the creation of noise representations for the following H-inf Adaptive Noise Cancellation, which adaptively removes eye artifacts from EEG, a notch filter (60Hz) was applied to the EOG data. This modification to the pipeline was effective in removing power line noise from the EEG signals (Figure 33).

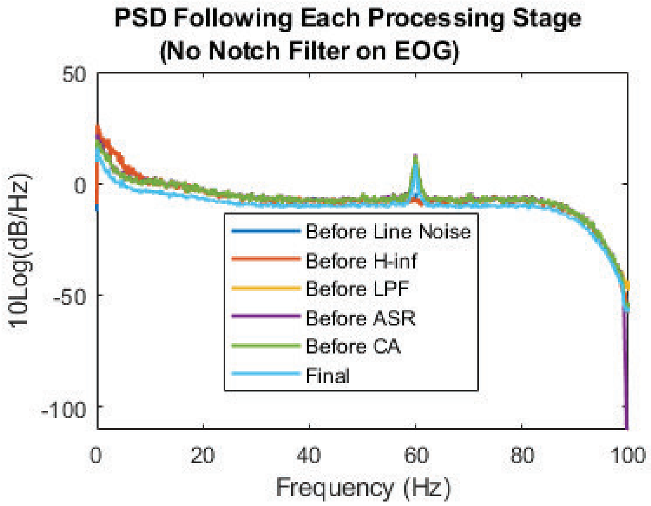


Figure 32: Power Spectral Density (PSD) following each processing step when the notch filter is not included, revealing continued power line contamination.

##### *9.5. EMG Cleaning Methodology*

*9.5.1. Single-Stage Blind Source Separation (BSS)* For ICA, one major drawback is that it is time-consuming and computationally demanding. The specific implementation of ICA for all single-stage ICA cleaning methods is SOBI. This was selected based on research that suggests SOBI is both faster and more accurate than other ICA implementations (JADE, Infomax, and FastICA) [52], while retaining the ability to accurately isolate EEG sources [51]. CCA blind source separation involves comparing a signal with a time lagged version of itself. CCA implementations in this research incorporate a time lag of one point, as previous studies have indicated that this lag facilitates better discrimination between EEG and EMG components [54]. Components resulting from ICA and CCA are then assessed based on criteria discussed in Section ??.

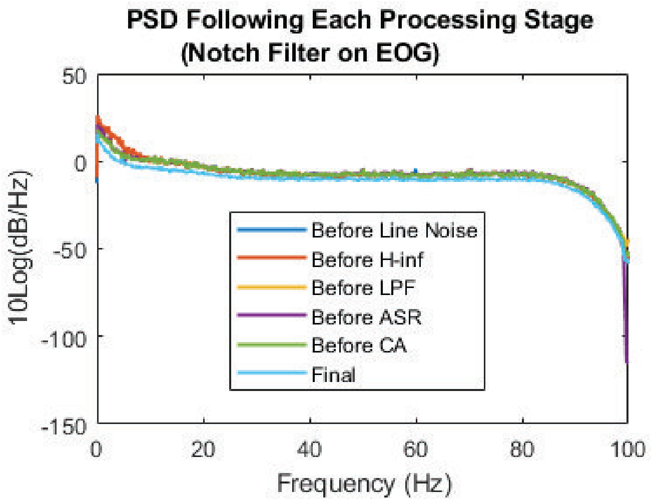


Figure 33: Power Spectral Density (PSD) for each processing step when the notch filter is included. The inclusion of a notch filter on the passive EOG sensors led to effective removal of power line noise.

*9.5.2. EEMD Parameter Selections* EEMD required the selection of three primary parameters: the noise level (expressed as a portion of the EEG data standard deviation), the number of ensembles, and the number IMFs to decompose each channel into. Previous studies have provided insights into these parameter selections. For the noise level, research [26] suggests that values over 0.1 times the standard deviation of the input data effectively stabilize the EEMD decomposition. Another study [54] empirically found that a noise level of 0.2 was effective. This research employed a noise level of 0.2 times the standard deviation of the input data.

Regarding the number of ensembles, [26] compared 10, 50, and 100 ensembles but found minimal differences in the resulting IMFs. In [53], the authors found that anything over five ensembles effectively extracted IMFs from EEG data. Due to the significant impact on processing speed, this study adopted 10 ensembles for all EEMD implementations.

The final parameter is the number of IMFs to extract from each channel. Based on [53]), where the authors considered different neural modalities and the effects of IMF count per modality, six IMFs were chosen as a suitable number of distinct IMFs for the EEMD process.

After EEMD decomposition, a subset of IMF's is selected for further analysis in the BSS stage. To isolate potentially artifactual components, an autocorrelation value of 0.95 was chosen, which followed the recommendations of [56, 54]. This selection does not significantly affect the subsequent cleaning process, as this threshold only isolates potentially artifactual components and does not directly remove any data. The resulting group of potentially artifactual components were then passed to the artifactual component assessment methods.

*9.5.3. Artifactual Component Assessment* The EMG cleaning methods utilized in this research yield a set of EEG components that required assessment for the identification and removal of artifactual components. To select and remove these artifactual components, specific criteria based on the characteristics of EMG data were incorporated, namely autocorrelation values [53] and spectral content [55]. While autocorrelation has been used as a component selection criteria in previous research, there is a lack of guidance on selecting specific threshold values. Additionally, due to the unique spectral content of neural signals during overt speech, it was not appropriate to set correlation thresholds solely based on past research findings.

To address these challenges, the presented analysis explored two component rejection methods: spectral content and autocorrelation thresholds. These rejection methods were applied to components obtained from all four types of EMG removal methods (ICA, CCA, EEMD-ICA, EEMD-CCA), which enabled a comprehensive and comparative analysis.

The spectral thresholds were created following the suggestions outlined in [55]. According to the authors, a component was considered to be of EMG origin if the average power within the 15 – 30 Hz frequency range constitutes at least 12.5% of the total power within the 0.1 – 30 Hz frequency range. The choice of 12.5% as a threshold was empirically determined to be effective in removing simulated EMG data from EEG, but the authors cautioned against solely relying on this threshold without additional analysis.

In addition to spectral content thresholds, autocorrelation thresholds have been employed for component rejection in the two-stage EEMD-CCA EMG removal method detailed in [54]. The authors of that study identified an effective range for the autocorrelation thresholds in the BSS stage, ranging from 0.8 to 0.95, with higher values indicating a higher likelihood of the component being EEG data. However, they also emphasized the need to assess multiple threshold levels to determine the appropriate threshold for specific applications.

By investigating both spectral content and autocorrelation thresholds as component rejection methods, this analysis provided a comprehensive evaluation of the effectiveness of these approaches across various EMG removal methods. This allowed for a more informed selection and removal of artifactual components, ultimately improving the quality of the EEG data and enhancing the reliability of subsequent analysis and interpretation.

To evaluate the performance of the proposed component assessment methods alongside a standard EMG removal algorithm, the ICLabel module provided by EEGLab was utilized for the single-stage Independent Component Analysis (ICA) removal [87]. EEGLab is a widely used software toolbox for processing and analyzing electroencephalography (EEG) data, offering various algorithms and tools for artifact removal and signal processing. The ICLabel module within EEGLab is specifically designed for the automatic identification and labeling of independent components obtained through ICA decomposition. By applying ICLabel to the single-stage ICA removal, the objective was to compare the performance of the proposed methods with a well-established and commonly used algorithm for EMG removal.

*9.5.4. Artifactual Component Rejection Threshold Selection* To ensure a more accurate and reliable analysis, component rejection methods, including autocorrelation and spectral content thresholds, were applied to all four types of EMG removal methods. However, blindly using previously suggested threshold ranges from past research was deemed inappropriate. Instead, Kmeans clustering [57] was employed to determine threshold levels that would yield reliably distinct cleaned datasets.

The single-stage EMG removal methods produce fewer components compared to the two-stage methods. Consequently, iterating locally around the provided threshold ranges resulted in minimal differences in the resulting signals as the limited ranges often identified the exact same components as artifactual. To overcome this issue and ensure unique cleaned datasets, all components for each participant across all sessions were collected, and Kmeans clustering was applied to the autocorrelation and spectral content threshold levels.

Ideally, the number of optimum groups in the Kmeans clustering would be determined by a measure of grouping, such as with a gap evaluation [88] or a Calinski-Harabasz criterion [89]. However, calculating optimal groups using such measures proved to be unstable in this context. For instance, hinge loss calculations suggested an optimal grouping of two groups, while the Calinski-Harabasz criterion resulted in up to 10 groups. Additionally, considering the entire range of $0 - 1$ for threshold levels would lead to groupings that include components with characteristics significantly different from the expected properties of EEG and EMG components. For instance, including threshold levels that covered components introduced by the Gaussian noise addition mechanism of the two-stage EEMD-CCA removal process (e.g. spectral content thresholds around 0.5 or very low autocorrelation thresholds) would guarantee that the cleaning process was retaining artifactual components.

To address these challenges, the Kmeans clustering search was restricted to specific ranges. For spectral content thresholds, the Kmeans search range was set to 0-0.3 for autocorrelation values, 0.6-1 for spectral content thresholds, and $0.4 - 1$ for the ICLabel comparison (which represents the confidence that a component is EMG activity [87]). Furthermore, the search was restricted to three total groupings. These restrictions in the Kmeans clustering allowed for an analysis of low, medium, and high levels of cleaning across a range of theoretically viable threshold levels.

After conducting the Kmeans analysis, the average threshold levels across all participants were selected as the final threshold levels for the EMG removal methods. This choice was made to ensure fairness and comparability in the analysis, as using participant-specific thresholds would reduce the generalizability of this threshold selection framework.

The participant-averaged thresholds that resulted from the Kmeans clustering approach are of low variance, which demonstrates the similarities in component

Table 9: Cleaning method average threshold levels and variances.

| | Mean and Variance for Threshold Levels | | | | | |
|---|---|---|---|---|---|---|
| **Cleaning Method** | **Mean** | **Variance** | **Mean** | **Variance** | **Mean** | **Variance** |
| **EMD-CCA (corr)** | 0.703 | 0.023 | 0.799 | 0.033 | 0.891 | 0.023 |
| **EMD-ICA (corr)** | 0.696 | 0.022 | 0.786 | 0.039 | 0.882 | 0.027 |
| **CCA (corr)** | 0.711 | 0.011 | 0.807 | 0.012 | 0.887 | 0.013 |
| **ICA (corr)** | 0.711 | 0.013 | 0.806 | 0.012 | 0.885 | 0.011 |
| **EMD-CCA (PSD)** | 0.086 | 0.016 | 0.152 | 0.017 | 0.223 | 0.012 |
| **EMD-ICA (PSD)** | 0.087 | 0.017 | 0.151 | 0.013 | 0.224 | 0.007 |
| **CCA (PSD)** | 0.094 | 0.012 | 0.138 | 0.024 | 0.193 | 0.038 |
| **ICA (PSD)** | 0.068 | 0.012 | 0.126 | 0.014 | 0.202 | 0.014 |
| **ICA (ICLabel)** | 0.477 | 0.032 | 0.686 | 0.039 | 0.865 | 0.039 |

characteristics across participants (Table 9). Additionally, the resulting thresholds for the different component rejection methods (i.e. the use of auto-correlation or spectral content) were similar regardless of the number of stages or blind source separation selection. The thresholds presented here overlap the proposed thresholds provided by the previous research that motivated the component rejection methods, specifically that the thresholds for spectral-based rejection methods contained the 12.5% threshold proposed in [55] whereas the thresholds for auto-correlation on the blind source separation contained the lower bound of the proposed auto-correlation threshold of 0.80, as proposed in [5].

*9.5.5. Effectiveness of EMG Cleaning Methods* The performance of the different EMG removal methods was evaluated using multiple types of assessment metrics: signal Root-Mean-Square error (RMSE) by condition, spectral analysis by condition, artifactual component reconstruction, and source analysis.

To assess how the different EMG removal methods directly affected the signal, the cleaned EEG data was segmented into periods of overt speech production and rest. For each EMG removal method, the RMSE values between the solely pre-processed and cleaned EEG signals were calculated separately for speech and non-speech conditions. The hypothesis was that the RMSE would be lower for non-speech segments, as there should be minimal EMG activity to be removed, while a higher RMSE value was expected during overt speech production.

A spectral analysis was conducted by calculating the power spectral density (PSD) of all cleaned datasets and comparing it with the PSD calculated from the solely pre-processed data. An example of this PSD comparison is presented with Figure 34. In this comparison, the EMG power spectral density exhibits power in the lower frequencies. This is likely due to EEG contamination of the EMG data as no past research has indicated EMG contains significant power in these bands. This finding reinforces the idea that using adaptive filtering techniques that rely on facial EMG noise references might not be practical due to the low-frequency power from EEG contamination that could still be present. Consequently, applying an adaptive filter to remove this noise may inadvertently lead to the removal of potentially important EEG data.

Metrics were computed by comparing the RMSE of the PSD before and after

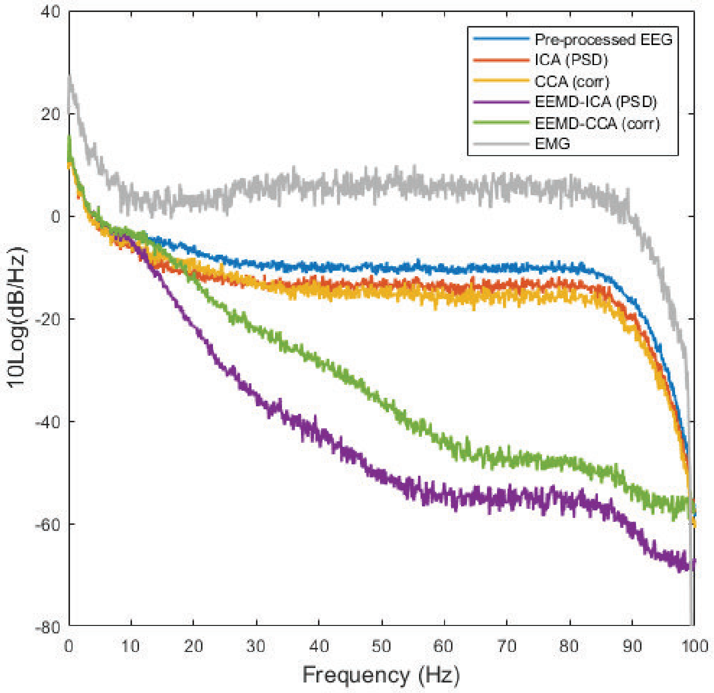


Figure 34: Comparison of the Power Spectral Densities for the four cleaning methods, the solely pre-processed data, and the EMG signal collected at the Orbicularis Oris. The EMG signal includes information in the lower frequency sub-bands, which is due to the contamination of EMG with EEG.

cleaning for different EEG sub-bands. Additionally, relative power changes between EEG sub-bands before and after EMG cleaning were analyzed. The hypothesis was that higher frequency bands (beta, gamma) will exhibit higher distortion levels compared to lower frequency bands (delta, theta, alpha) due to the overlap between the spectral content of EMG and EEG data in the higher bands.

Another method for comparing EMG removal methods involved analyzing the identified artifactual components. Instead of removing artifactual components, the EEG components were instead removed, and the theoretical artifact was reconstructed. The correlation between the reconstructed artifact and the signals collected by EMG sensors placed around the mouth (superior and inferior orbicularis) was assessed.

To provide additional context in how this reconstructed artifact propagates across all EEG channels, Figure 35 presents the topoplot of the reconstructed artifact, centered around one instance of the onset of speech. As expected, larger amplitudes for the reconstructed artifact were found in the channels closest to the face, with some additional high frequency content is evident at channel locations closest to the neck, which potentially represents muscle activity due to neck or head movement. This specific reconstruction was based on data cleaned with the EEMD-ICA method.

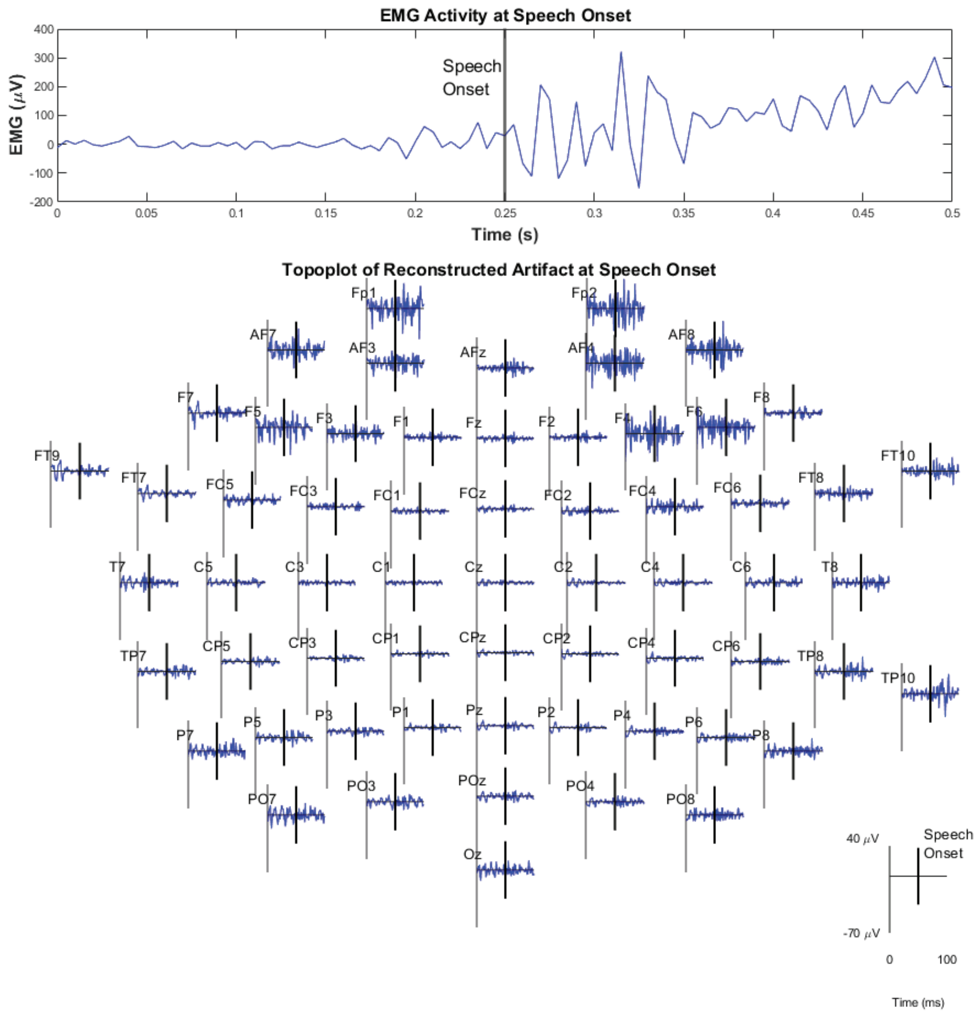


Figure 35: Propagation of the reconstructed artifact across all EEG channels. This is plotted alongside the collected EMG data. All plots are centered around one instance of the onset of speech.

To address the concern of removing speech-related EEG information, the solely pre-processed and cleaned EEG data were subjected to independent component analysis (ICA) separately for each condition. The resulting independent components were then linked to dipoles using EEGlab's DIPFIT module [90]. Dipole fitting is a technique that attempts to determine where activations are occurring in the brain based on the resulting scalp EEG.

To perform dipole fitting, a model is created to estimate how electrical signals from neural sources in the brain propagate to the scalp surface, which is the so-called forward model. The forward model considers the conductive properties of the head tissues, the geometry of the head, and the electrode positions. The inverse problem is the core concept behind dipole fitting. Given the recorded EEG data, the goal is to estimate the distribution of current dipoles (electrical sources) inside the brain that best explains the observed scalp potentials [91]. Fitted dipoles can then be clustered and the resulting centroid of each cluster can be used to gain insights into sources related to overt speech production. The resulting centroids can then be further localized to Brodmann areas based on a participant-independent brain model. While the lack of participant-specific MRI data or recorded electrode positions may reduce the accuracy of the resulting localization, careful application of the EEG headsets during collection was followed to allow for accurate and consistent electrode placement across participants and sessions.

Brodmann areas represent common neurological areas, segmented based on the cytoarchitectural properties of neural tissue [92]. This segmentation enables functional analysis of different brain regions. It is expected that neural activation during speech production will be localized to speech centers such as Broca area and Wernicke area [93]. From a limited standpoint, Broca area is typically only associated with BAs 44 and 45, whereas Wernicke area is typically only associated with BA 22. However, since MRI was not collected for any of the participants, this research instead employs a brain atlas developed by [94], where the authors mapped Brodmann areas to a participant's MRI scan. This means that localization of dipoles for the participants collected through this research won't be perfectly aligned to the MRI scan, as the MRI scan is not participant-specific. For this reason, this research expanded the number of BAs selected to represent the two primary speech areas based on research that examines larger sets of speech-related BAs [95]. The BAs corresponding to speech centers used for this research are presented in Table 10. The hypothesis for this source analysis was that an EMG removal method that effectively preserves speech-related EEG components would result in higher retention of components in clusters associated with these speech areas.

Table 10: Brodmann areas associated with Wernicke and Broca areas.

| Region | Primary Speech Function | Brodmann Area |
|---|---|---|
| Broca Area | Speech production | 44 / 45 / 46 |
| Wernicke Area | Speech perception | 21 / 22 / 41 / 42 |

To investigate the relationship between the defined metrics and decoding accuracy, a decoding task was conducted. The task involved using two types of models, a Recurrent Neural Network (RNN) and a Convolutional Neural Network (CNN), to classify the articulatory features of individual phonemes. For this model performance assessment, relatively simply variations of CNNs and RNNs were employed. The architectures for these models are presented with Figure 36. These models (eight variations of models for each dataset) were selected to give a relatively wide range of model capacities. All models were trained with a training / validation / testing dataset split of 70/20/10 to help prevent over fitting. Training employed ADAM optimization [?] and a batch size of 64 for 50 epochs with an early stopping criteria of 10 epochs. Performance of the models was evaluated based on accuracy of the resulting classifications. As an additional check on the effectiveness of the EMG cleaning metrics, each of the above defined metrics was correlated with the changes in model performance and the resulting correlation strengths and p-values were obtained. Correlation was obtained using Pearson's correlation analysis [96], a common correlation metric that compares the strength and direction of linear relationships between two variables.

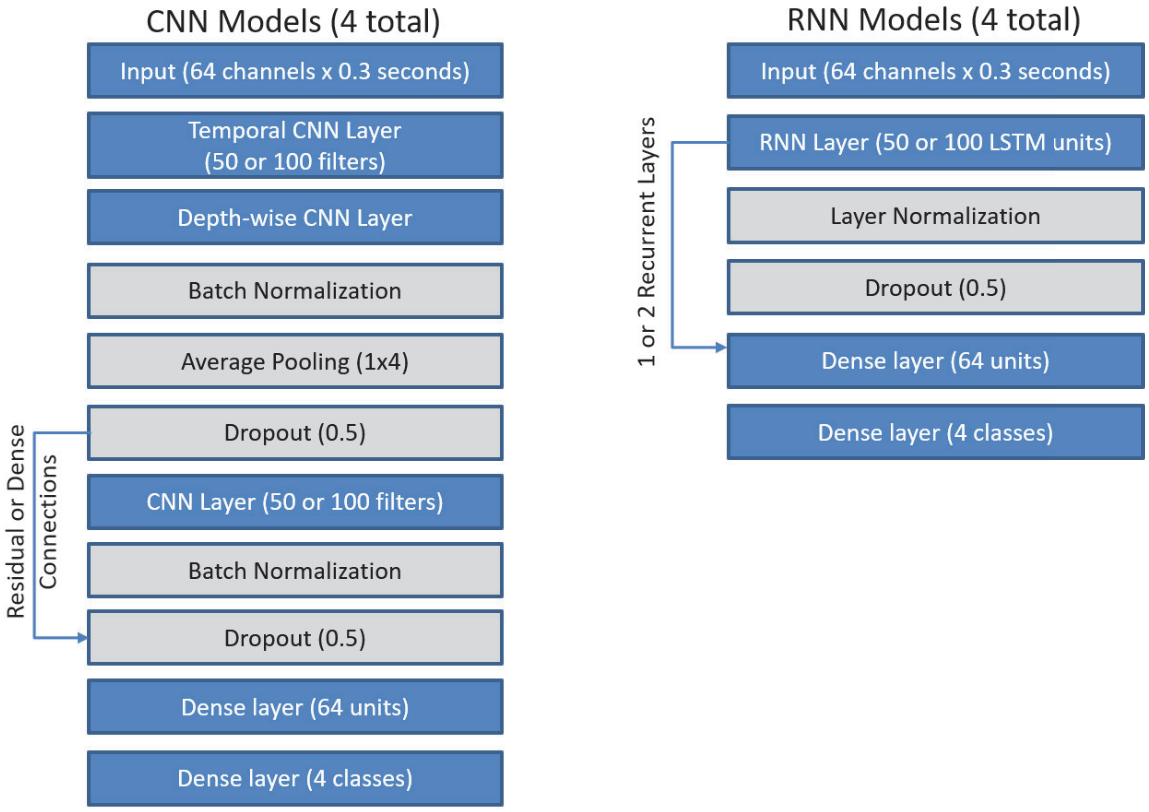


Figure 36: Model architectures employed for discrete decoding task. Blue indicates architecture design choices and gray indicates regularization selections.

*9.5.6. Signal RMSE analysis* Table 11 presents the findings from the signal distortion analysis. While the two-stage methods exhibited less distortion compared to one-stage methods, the RMSE difference between the two approaches was found to be similar. For component rejection methods, correlation-based rejection methods were observed to lead to high distortion during speech (but not in non-speech conditions), resulting in a higher RMSE difference, which may indicate correlation-based rejection methods are able to focus on speech-related artifacts more effectively than spectral methods. ICLabel demonstrated significantly lower distortion in both speech and non-speech periods and a lower difference, indicating less cleaning generally. Regarding threshold levels, higher distortion was expected for higher levels of cleaning, but the distortion ratio remained relatively even across the threshold levels. In the context of blind source separation selection, we found that using independent component analysis (ICA) resulted in more distortion during both speech and non-speech conditions.

Table 11: Signal RMSE metrics by cleaning method characteristic.

| | | RMSE Signal | | |
|---|---|---|---|---|
| | | **Speech** | **Nonspeech** | **Difference** |
| **Stages** | **Single Stage** | 6.35 | 5.02 | 1.34 |
| | **Two Stage** | 5.96 | 4.61 | 1.35 |
| **Comp. Rej.** | **ICLabel** | 2.91 | 2.42 | 0.49 |
| | **PSD** | 5.98 | 4.83 | 1.14 |
| | **Auto-correlation** | 6.34 | 4.84 | 1.50 |
| **Thresh. Level** | **Low** | 5.15 | 3.88 | 1.27 |
| | **Medium** | 5.90 | 4.61 | 1.29 |
| | **High** | 6.59 | 5.31 | 1.28 |
| **BSS** | **CCA** | 5.99 | 4.70 | 1.29 |
| | **ICA** | 6.38 | 4.94 | 1.44 |

The Pearson's correlation analysis is presented with Figure 37. In all following Pearson's correlation figures, each label on the x-axis for the bottom figure includes whether the correlation strength was consistently negative or positive for all participants, or whether the correlation direction was not consistent across participants ('mixed'). The analysis of the RMSE spectral metrics (Figure 37) revealed only two correlations, Signal RMSE for both the speech and non-speech conditions, that exhibited high confidence levels when compared against the null hypothesis. However, it is worth noting that the RMSE difference, despite displaying correlation, was deemed an unreliable indicator of model performance. While it does suggest a relationship between the cleaning methods and the level of signal distortion during both conditions, it is likely not a good metric for insights into model performance, but instead served primarily to inform on condition-dependent differences in cleaning methods.

*9.5.7. Spectral RMSE and relative spectral change analysis* Tables 12 and 13 present the findings from spectral RMSE analysis. Comparing the two-stage and one-stage approaches, we found significantly less distortion in the Delta, Theta, and Alpha bands for two-stage approaches, with the effect being less pronounced in non-speech conditions. This was expected as the two-stage methods allow for the deconstruction of the signal into more components, theoretically helping to avoid the removal of EEG data. Regarding component rejection methods, correlation-based rejection introduced more distortion in the Delta, Theta, and Alpha bands compared to power spectral density (PSD) rejection. However, ICLabel exhibited lower distortion in the Alpha and Beta bands. As for threshold levels, increasing levels of distortion were observed with higher thresholds, except in the upper gamma band. During non-speech conditions, minimal distortion was observed when employing low threshold levels, particularly when compared to the distortion observed during speech. Lastly, in blind source separation selection, the use of independent component analysis (ICA) resulted in more distortion in all bands during speech. However, during non-speech periods, ICA showed less distortion in the Delta, Theta, Alpha, and Beta bands.

The trends for effective metrics in the RMSE space (Figures 38 and 39) matched relatively well between speech and non-speech conditions across cleaning methods,

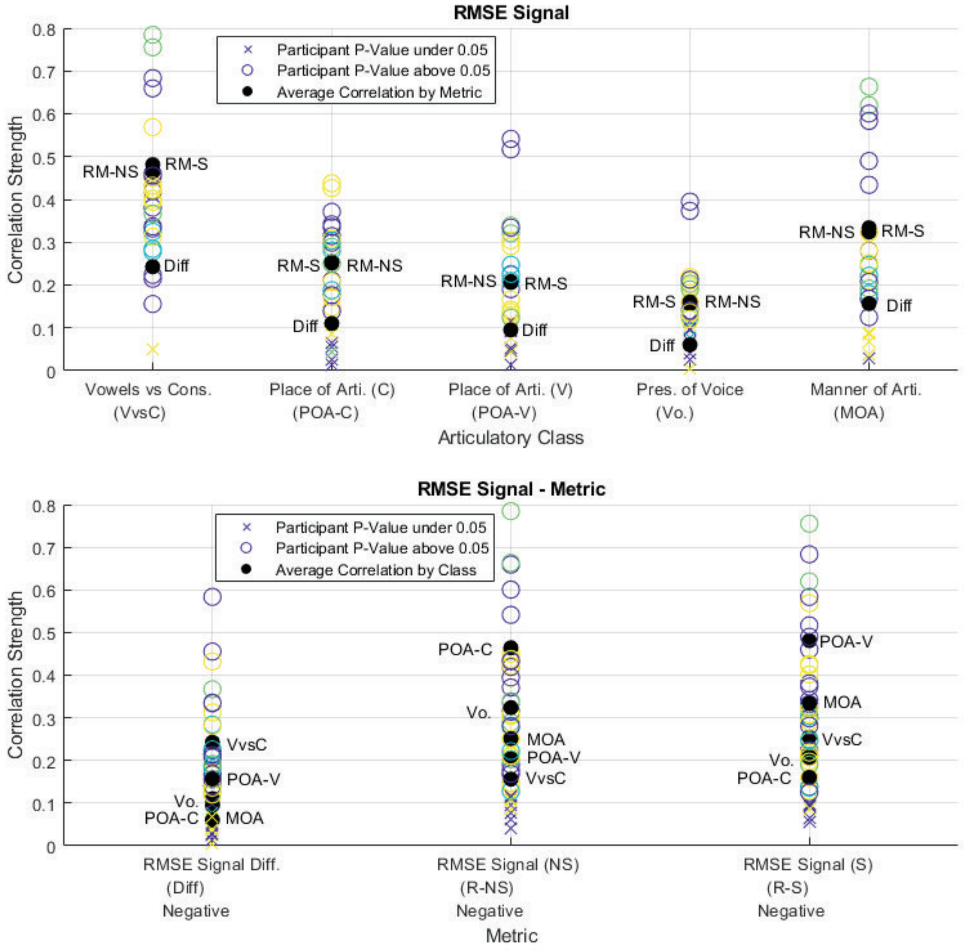


Figure 37: Pearson correlation analysis between the RMSE Signal metrics and decoding model performance for all classes, viewed as class and metric averages. Each label on the x-axis for the bottom figure includes whether the correlation strength was consistently negative or positive for all participants.

particularly for the Delta and Theta RMSE values. These metrics show the largest correlation strengths and highest confidence across all investigated metrics, particularly the "Vowels vs Cons." discrete decoding class.

Tables 14 and 15 present the findings from the changes in relative power analysis. Comparing the two-stage and one-stage approaches, larger changes in relative power across all bands were observed for two-stage approaches, with the lower and upper gamma bands being almost entirely removed. When considering component rejection methods, larger decreases in the gamma bands were observed for the auto-correlation rejection method as compared to power spectral density (PSD) rejection, providing further support that auto-correlation rejection methods are more effective in removing gamma activity. ICLabel demonstrated effective preservation of the beta band, but,

Table 12: Spectral RMSE metrics by cleaning method characteristic (low frequency bands).

| | | RMSE Spectral | | | | | |
|---|---|---|---|---|---|---|---|
| | | Low Frequency Bands | | | | | |
| | | Delta | | Theta | | Alpha | |
| | | S | NS | S | NS | S | NS |
| **Stages** | **Single Stage** | 5.29 | 3.08 | 5.94 | 1.02 | 2.31 | 1.45 |
| | **Two Stage** | 1.05 | 0.59 | 0.98 | 0.38 | 0.71 | 0.98 |
| **Comp. Rej.** | **ICLabel** | 1.14 | 1.86 | 0.91 | 0.42 | 0.32 | 0.42 |
| | **PSD** | 2.89 | 1.79 | 3.16 | 0.69 | 1.45 | 1.24 |
| | **Auto-correlation** | 3.86 | 2.15 | 4.25 | 0.77 | 1.73 | 1.24 |
| **Thresh. Level** | **Low** | 2.75 | 0.85 | 2.97 | 0.34 | 1.13 | 0.61 |
| | **Medium** | 3.76 | 2.26 | 3.98 | 0.73 | 1.52 | 1.14 |
| | **High** | 3.86 | 3.33 | 4.12 | 1.15 | 1.88 | 1.75 |
| **BSS** | **CCA** | 3.71 | 2.69 | 3.48 | 0.78 | 1.37 | 1.20 |
| | **ICA** | 4.49 | 2.05 | 4.31 | 0.72 | 1.62 | 1.18 |

Table 13: Spectral RMSE metrics by cleaning method characteristic (high frequency bands).

| | | RMSE Spectral | | | | | |
|---|---|---|---|---|---|---|---|
| | | High Frequency Bands | | | | | |
| | | Beta | | Low Gamma | | Upper Gamma | |
| | | S | NS | S | NS | S | NS |
| **Stages** | **Single Stage** | 1.29 | 0.49 | 0.92 | 0.31 | 0.56 | 0.18 |
| | **Two Stage** | 1.34 | 0.54 | 1.05 | 0.39 | 0.64 | 0.23 |
| **Comp. Rej.** | **ICLabel** | 0.37 | 0.29 | 0.46 | 0.20 | 0.28 | 0.16 |
| | **PSD** | 1.25 | 0.52 | 0.91 | 0.34 | 0.55 | 0.19 |
| | **Auto-correlation** | 1.36 | 0.51 | 1.04 | 0.36 | 0.63 | 0.21 |
| **Thresh. Level** | **Low** | 1.03 | 0.37 | 0.84 | 0.29 | 0.52 | 0.18 |
| | **Medium** | 1.25 | 0.49 | 0.93 | 0.33 | 0.57 | 0.20 |
| | **High** | 1.37 | 0.61 | 1.00 | 0.36 | 0.60 | 0.21 |
| **BSS** | **CCA** | 1.25 | 0.50 | 0.94 | 0.33 | 0.57 | 0.19 |
| | **ICA** | 1.32 | 0.49 | 0.98 | 0.34 | 0.59 | 0.20 |

as the beta band has not been thought to be informative for speech processes, this finding may have minimal importance. Regarding threshold levels, we found that a lower percentage of power in the upper bands was removed during speech conditions as cleaning levels increase when considering relative power, which matches intuitions for cleaning levels. Finally, in blind source separation selection, the differences between blind source separation (BSS) methods were less pronounced. However, the use of canonical correlation analysis (CCA) resulted in less distortion in the upper bands compared to independent component analysis (ICA). It was assumed the use of CCA would lead to higher levels of distortion in the upper bands, but, as both ICA and CCA investigated the use of both spectral and auto-correlation rejection methods, this assumption may not have been valid. Instead, ranking the resulting components by auto-correlation may be as effective for either type of blind source separation technique.

Relative changes in power (Figures 40 and 41) may be a less effective type of metric to deduce model performance than spectral RMSE, as very few participants achieve statistical significance (p ¡ 0.05). The ones that did achieve statistical significance have a correlation strength of around 0.2, which is considered a low strength of association.

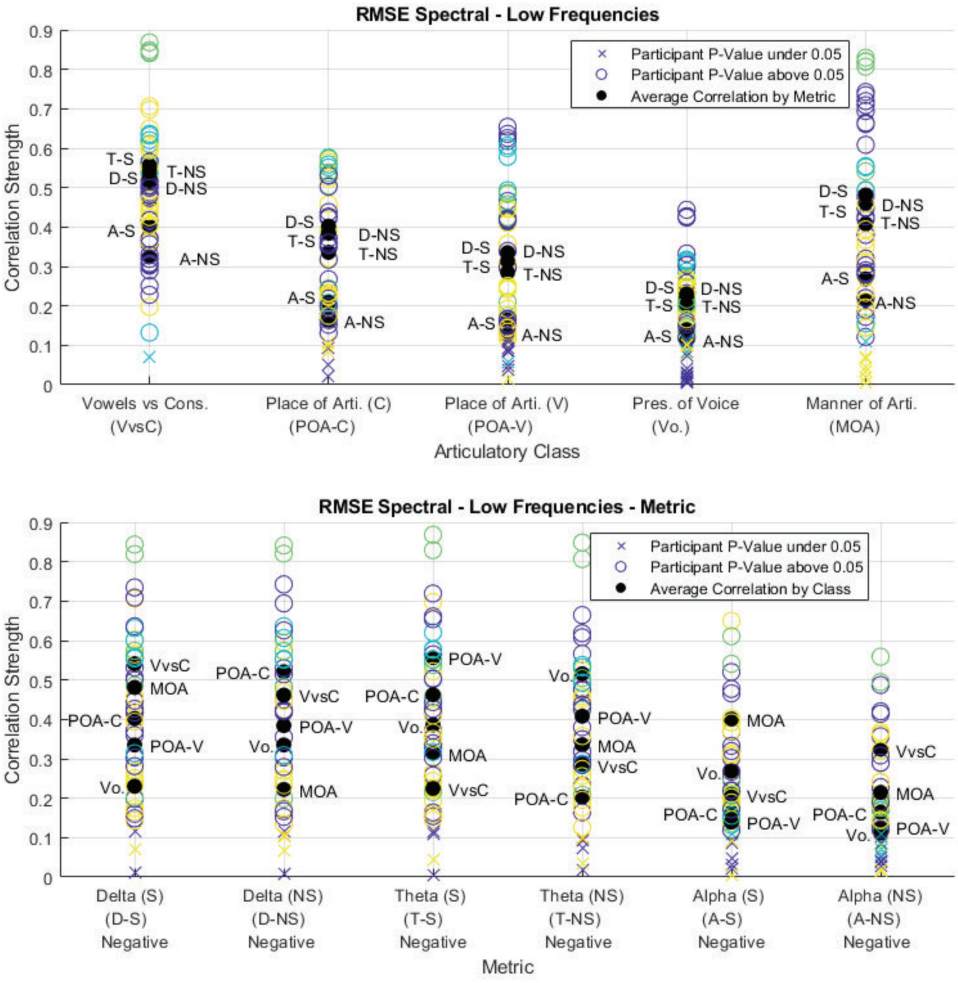


Figure 38: Pearson correlation analysis between the RMSE Spectral metrics (low frequencies) and decoding model performance for all classes, viewed as class and metric averages.

*9.5.8. Reconstructed artifact and source analysis* Table 16 presents the findings from the analysis of the reconstructed artifact. For all cleaning methods, the highest power of the reconstructed artifact was always either at electrode FP1 or FP2, indicating these methods' effectiveness in removing facial EMG contamination as these sensors are closest in proximity to the articulatory system. Comparing the two-stage and one-stage approaches, we found that the two-stage method generally removed less data, as evidenced by the lower values for average and highest power of the reconstructed artifact. Additionally, the correlation between the localized data and the original data was minimal, but slightly lower in the two-stage method. Moving on to component rejection methods, we observed similar R-squared and correlation values compared to power spectral density (PSD) rejection, with slightly less data removal. Notably, ICLabel

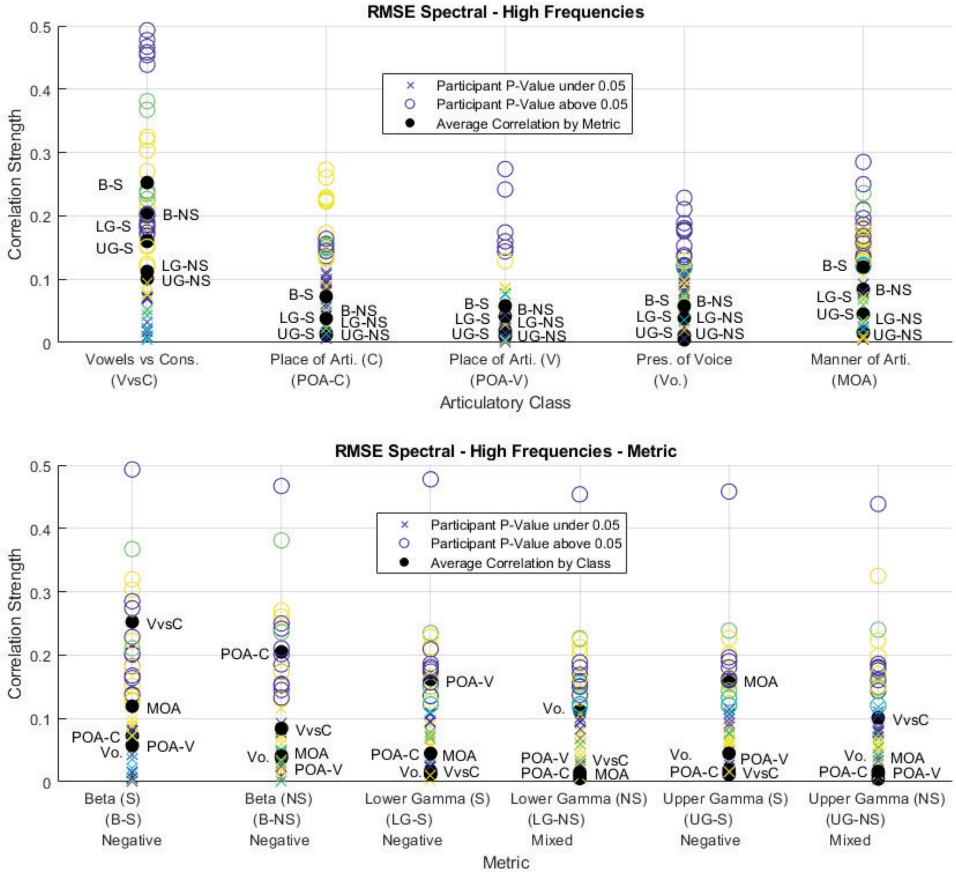


Figure 39: Pearson correlation analysis between the RMSE Spectral metrics (high frequencies) and decoding model performance for all classes, viewed as class and metric averages.

Table 14: Relative spectral metrics by cleaning method characteristic (low frequency bands).

| | | Relative Power Differences | | | | | |
|---|---|---|---|---|---|---|---|
| | | Low Frequency Bands | | | | | |
| | | Delta | | Theta | | Alpha | |
| | | S | NS | S | NS | S | NS |
| **Stages** | **Single Stage** | 0.57 | 0.52 | 0.46 | 0.36 | 0.20 | 0.02 |
| | **Two Stage** | 0.85 | 0.72 | 0.87 | 0.72 | 0.55 | 0.32 |
| **Comp. Rej.** | **ICLabel** | 0.19 | 0.18 | 0.21 | 0.20 | 0.20 | 0.22 |
| | **PSD** | 0.70 | 0.62 | 0.62 | 0.51 | 0.22 | 0.04 |
| | **Auto-correlation** | 0.69 | 0.60 | 0.66 | 0.52 | 0.46 | 0.23 |
| **Thresh. Level** | **Low** | 0.52 | 0.44 | 0.55 | 0.46 | 0.49 | 0.34 |
| | **Medium** | 0.57 | 0.49 | 0.59 | 0.49 | 0.48 | 0.29 |
| | **High** | 0.74 | 0.66 | 0.60 | 0.46 | 0.21 | -0.02 |
| **BSS** | **CCA** | 0.57 | 0.48 | 0.60 | 0.50 | 0.50 | 0.29 |
| | **ICA** | 0.63 | 0.54 | 0.61 | 0.50 | 0.41 | 0.20 |

exhibited the least correlation and the least removal of data. Regarding threshold levels, we observed a slight increase in correlation and power as the thresholds increased,

Table 15: Relative spectral metrics by cleaning method characteristic (high frequency bands).

| | | **Relative Power Differences** | | | | | |
|---|---|---|---|---|---|---|---|
| | | **High Frequency Bands** | | | | | |
| | | **Beta** | | **Low Gamma** | | **Upper Gamma** | |
| | | **S** | **NS** | **S** | **NS** | **S** | **NS** |
| **Stages** | **Single Stage** | -0.42 | -0.46 | -0.67 | -0.71 | -0.69 | -0.74 |
| | **Two Stage** | -0.59 | -0.66 | -0.97 | -0.98 | -1.00 | -1.00 |
| **Comp. Rej.** | **ICLabel** | -0.09 | -0.09 | -0.23 | -0.31 | -0.22 | -0.32 |
| | **PSD** | -0.58 | -0.63 | -0.75 | -0.77 | -0.75 | -0.77 |
| | **Auto-correlation** | -0.43 | -0.49 | -0.85 | -0.87 | -0.88 | -0.91 |
| **Thresh. Level** | **Low** | -0.29 | -0.34 | -0.66 | -0.70 | -0.69 | -0.73 |
| | **Medium** | -0.39 | -0.45 | -0.72 | -0.74 | -0.73 | -0.76 |
| | **High** | -0.60 | -0.65 | -0.83 | -0.86 | -0.85 | -0.87 |
| **BSS** | **CCA** | -0.39 | -0.45 | -0.73 | -0.74 | -0.75 | -0.77 |
| | **ICA** | -0.47 | -0.53 | -0.78 | -0.82 | -0.80 | -0.84 |

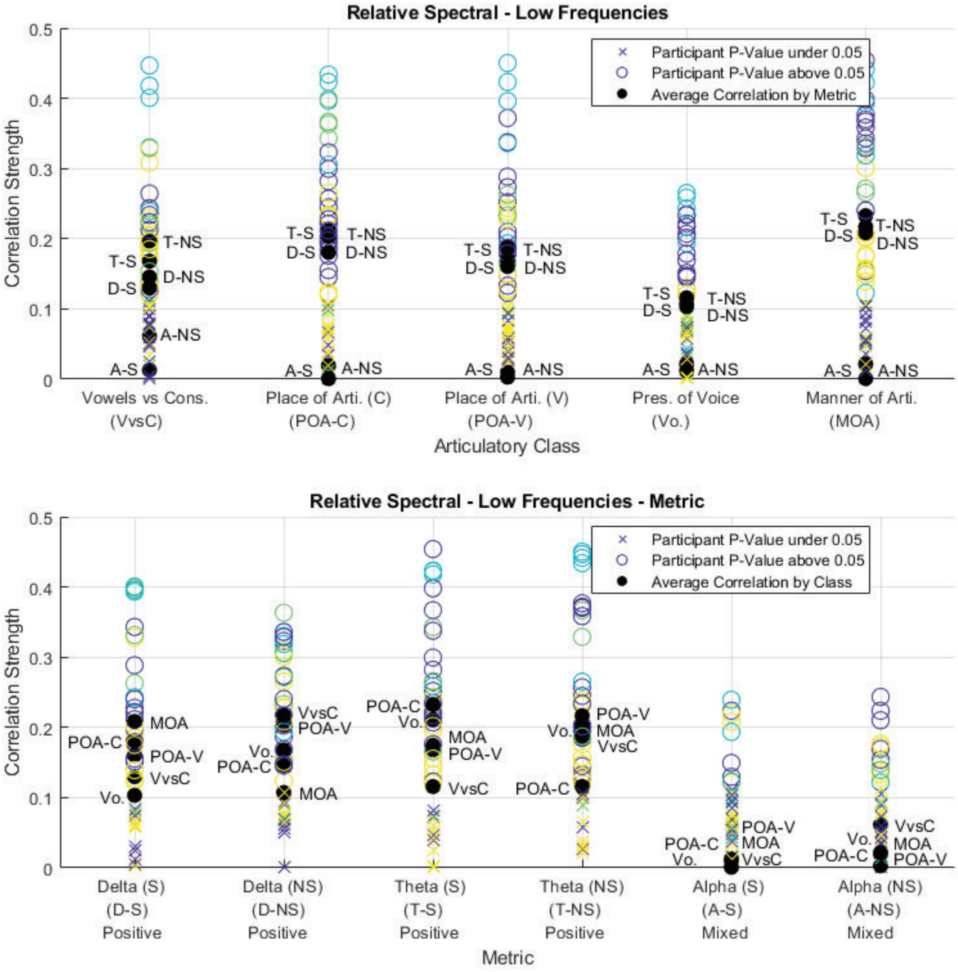


Figure 40: Pearson correlation analysis between the Relative Spectral metrics (low frequencies) and decoding model performance for all classes, viewed as class and metric averages.

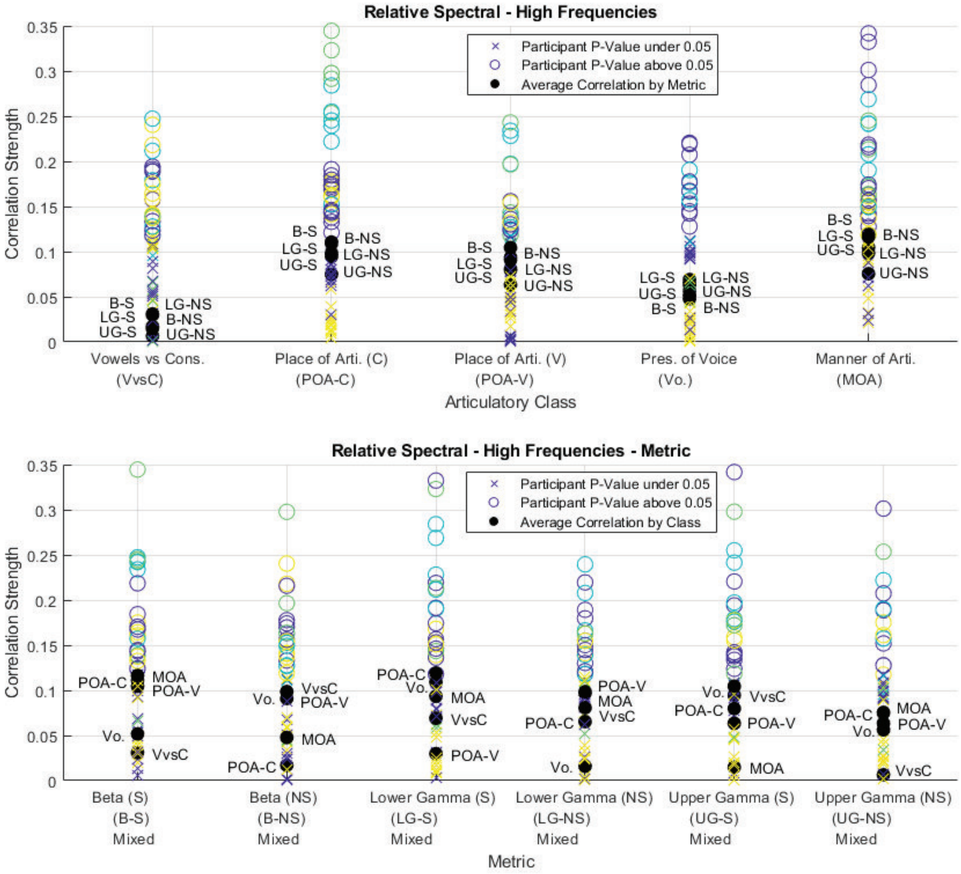


Figure 41: Pearson correlation analysis for the Relative Spectral metrics (high frequencies) and decoding model performance for all classes, viewed as class and metric averages.

although the difference was barely noticeable. Finally, in blind source separation selection, there was little difference in correlation between the methods. However, the use of CCA resulted in less data removal compared to other blind source separation techniques.

Table 16: Reconstructed artifact metrics by cleaning method characteristic.

| | | Artifact Reconstruction | | | |
|---|---|---|---|---|---|
| | | **EMG R-Squared** | **Corr.** | **Avg. Power** | **Highest Power** |
| **Stages** | **Single Stage** | 0.02 | 0.07 | 32.71 | 227.21 |
| | **Two Stage** | 0.01 | 0.05 | 27.68 | 211.44 |
| **Comp. Rej.** | **ICLabel** | 0.00 | 0.00 | 15.12 | 190.64 |
| | **PSD** | 0.01 | 0.06 | 30.10 | 213.87 |
| | **Auto-correlation** | 0.02 | 0.06 | 30.83 | 225.31 |
| **Thresh. Level** | **Low** | 0.01 | 0.04 | 24.12 | 203.67 |
| | **Medium** | 0.01 | 0.06 | 28.59 | 216.70 |
| | **High** | 0.02 | 0.07 | 34.39 | 233.62 |
| **BSS** | **CCA** | 0.02 | 0.06 | 29.15 | 211.99 |
| | **ICA** | 0.02 | 0.06 | 32.20 | 230.81 |

The reconstructed artifact metric correlation analysis is presented with Figure 42. The resulting R-squared and correlation values between the reconstructed artifact and the collected EMG were essentially 0, implying that the reconstructed artifact should be nothing like the EMG signal collected around the mouth for high model performance. This seems counter-intuitive, but only two EMG sensors were employed for collection around the mouth and the number of muscles involved during speech production is much higher [36]. The average correlation strength for the average and highest power of the reconstructed artifact was moderately negative, implying that less cleaning leads to better model performance. While this is likely true, as less cleaning means contamination would still be present for decoding, this finding may not be as informative due to the low number of EMG channels collected.

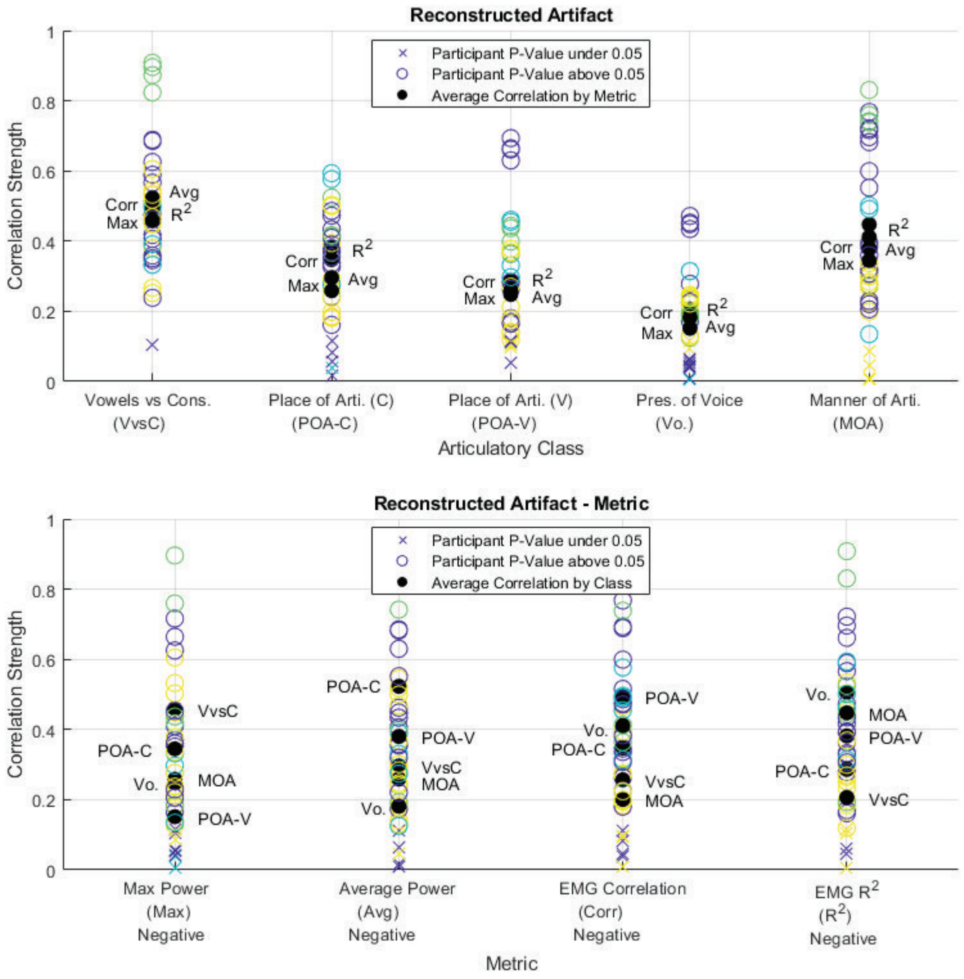


Figure 42: Pearson correlation analysis between the Reconstructed Artifact metrics and decoding model performance for all classes, viewed as class and metric averages.

In our investigation into the inclusion of dipole clusters in speech areas (Table

17), we compared the performance of two-stage and one-stage methods. The results revealed that a larger number of dipole clusters were successfully localized to speech areas using the two-stage approach, further supporting the idea that two-stage cleaning methods are more effective in separating EEG data from EMG data. Furthermore, we examined the impact of component rejection methods on dipole localization in Wernicke and Broca areas, finding the ICLabel rejection method exhibited superior performance in retaining components in both the Broca and Wernicke areas. In the analysis of threshold levels, interestingly, the medium threshold level retained a greater number of dipoles in the Broca area compared to the low or high thresholds, potentially indicating that the low and high thresholds led to under or over cleaning of the signal. Lastly, we evaluated the effect of blind source separation selection on dipole cluster localization, finding that the CCA selection retained more components in Broca area, whereas ICA retained more components in Wernicke area. As Wernicke area is associated with speech comprehension and Broca area is associated with speech production, the inclusion of dipole clusters in Broca area may be a more effective metric in the investigation of speech production.

Table 17: Source analysis metrics by cleaning method characteristic.

| | | Source Analysis | |
|---|---|---|---|
| | | **Wernicke** | **Broca** |
| **Stages** | **Single Stage** | 0.64 | 1.33 |
| | **Two Stage** | 0.94 | 2.21 |
| **Comp. Rej.** | **ICLabel** | 1.70 | 2.59 |
| | **PSD** | 1.00 | 1.97 |
| | **Auto-correlation** | 0.60 | 1.52 |
| **Thresh. Level** | **Low** | 1.25 | 1.63 |
| | **Medium** | 0.86 | 1.90 |
| | **High** | 0.40 | 1.74 |
| **BSS** | **CCA** | 0.32 | 1.72 |
| | **ICA** | 1.02 | 1.24 |

At least half of all participants for each discrete decoding task achieved p-values under 0.05 for a positive correlation between the number of dipole clusters localized in the Broca area and model performance (Figure 43). All participants found the correlation between model performance and the metric for Broca area to be likely the true hypothesis for "Vowels vs. Cons." This was promising and matched our expectations for overt speech production in that retaining EEG data produced in Broca area should lead to higher performance in decoding overt speech characteristics.

In addition to an assessment for the inclusion of sources in speech areas, this source analysis allowed for an understanding of the Brodmann areas activated during overt speech. During the experimental protocol, Brodmann areas localized over brain regions responsible for decision, memory, attention, and sensory functions were activated in addition to speech areas (Table 18).

*9.5.9. Decoding Performance Analysis* In the decoding performance assessment (Table 19), 2-stage methods always outperformed single stage approaches when not

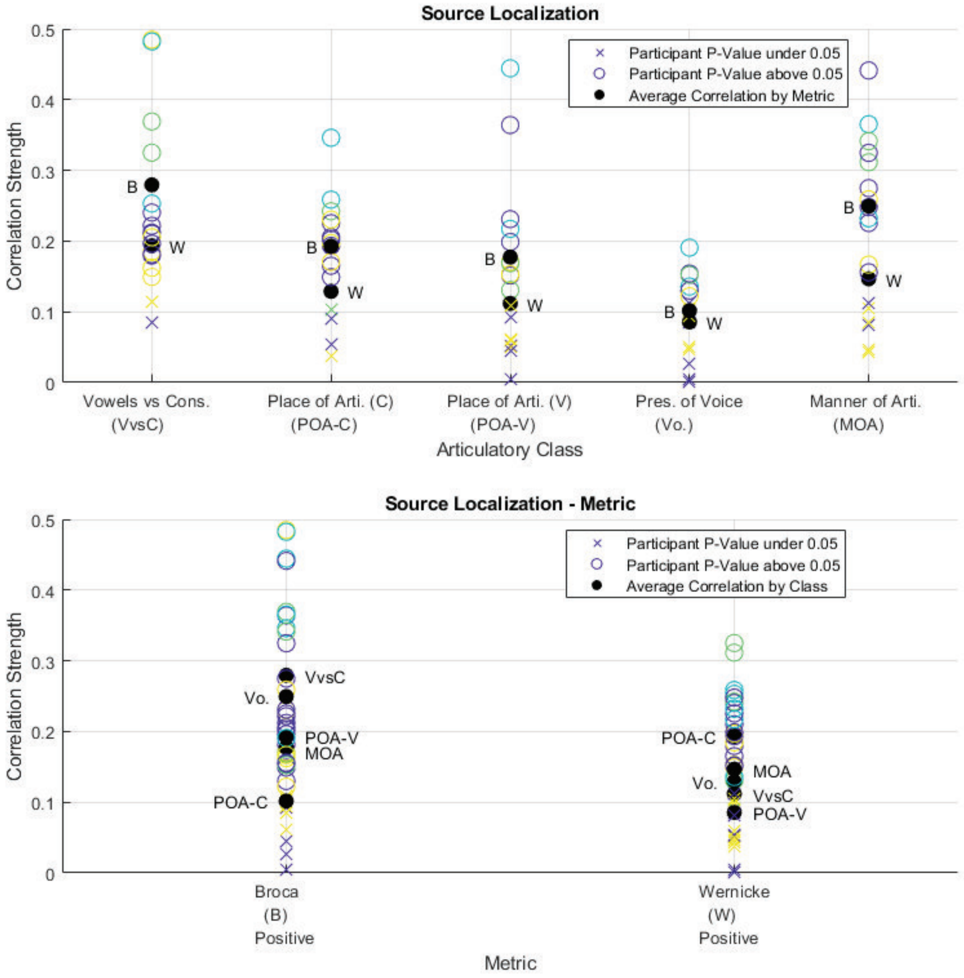


Figure 43: Pearson correlation analysis between the Source analysis metrics and decoding model performance for all classes, viewed as class and metric averages.

considering ICA cleaned with ICLabel EMG removal. ICLabel always outperformed all other cleaning methods, whereas the solely pre-processed data outperformed against every cleaned dataset. ICA (without ICLabel) outperformed CCA both with all implementations together and with just the 2-stage methods included. Lower levels of cleaning always led to higher performance, which makes sense as the model is able to use both EMG and EEG data for decoding. As there exists identical trends across discrete classes, regardless of the type of cleaning method employed, this pointed to the idea that model performance is highly linked to the input data distribution, rather than the output feature class formation.

*9.5.10. Method and Threshold Level Selection* In terms of signal RMSE, on average, data cleaned with two-stage approaches exhibited less distortion of the signal itself. For

Table 18: Brodmann areas activated during the overt speech task.

| BA | Count | Area | Language | Decision | Memory | Attention | Sensory | Motor |
|---|---|---|---|---|---|---|---|---|
| BA11 | 290 | Orbitofrontal cortex | | X | | | | |
| BA23 | 288 | Parahippocampal gyrus | | | X | | | |
| BA32 | 255 | Anterior cingulate cortex | | X | | X | | |
| BA24 | 217 | Anterior cingulate cortex | | X | | | | |
| BA26-29-30 | 189 | Cingulate cortex (all) | | X | X | X | | |
| BA18 | 149 | Visual cortex | | | | | X | |
| BA28 | 149 | Entorhinal cortex | | | X | | | |
| BA45 | 140 | Prefrontal cortex | X | | | | | |
| BA17 | 123 | Visual cortex | | | | | X | |
| BA22 | 122 | Temporal gyrus | X | | | | | |
| BA27 | 111 | Piriform cortex | | | | | X | |
| BA19 | 108 | Lateral occipital cortex | | | | | X | |
| BA37 | 98 | Fusiform gyrus | X | | X | | X | |
| BA7 | 89 | Posterior parietal cortex | | | | | | X |
| BA5 | 80 | Superior parietal lobule | | | | | X | |
| BA25 | 79 | Anterior cingulate cortex | | | | | | |
| BA40 | 72 | Inferior parietal lobule | X | | | X | X | |
| BA9 | 71 | Prefrontal cortex | | | X | | | |
| BA46 | 67 | Prefonrtal cortex | X | | | | | |
| BA35-36 | 49 | Parahippocampal gyrus | | | X | | X | |
| BA6 | 35 | Premotor cortex | | | | | | X |
| BA2 | 24 | Postcentral gyrus | | | | | X | X |
| BA4 | 24 | Precentral gyrus | | | | | | X |
| BA8 | 15 | Dorsolateral prefrontal cortex | | X | | X | | |
| BA44 | 9 | Pars opercularis | X | | | | | |
| BA39 | 7 | Angular gyrus | X | | X | | | |

Table 19: Decoding model performances for all discrete classes by cleaning method characteristic. Results are presented as percentage accuracy above chance level.

| | | **Model Performance by Discrete Decoding Class** | | | | | |
|---|---|---|---|---|---|---|---|
| | | **MOA** | **Voice** | **POA (V)** | **POA (C)** | **V vs C** | **Average** |
| **Stages** | **Single Stage** | 11.8% | 4.7% | 8.4% | 6.1% | 12.8% | 8.8% |
| | **Two Stage** | 14.5% | 5.9% | 10.1% | 7.5% | 15.1% | 10.6% |
| **Comp. Rej.** | **ICLabel** | 15.5% | 6.3% | 10.2% | 8.0% | 17.1% | 11.4% |
| | **PSD** | 12.9% | 5.2% | 9.2% | 6.7% | 13.6% | 9.5% |
| | **Auto-correlation** | 12.3% | 5.0% | 8.9% | 6.4% | 13.2% | 9.2% |
| **Thresh. Level** | **Low** | 14.0% | 5.6% | 9.7% | 7.2% | 15.1% | 10.3% |
| | **Medium** | 12.9% | 5.2% | 9.1% | 6.6% | 13.8% | 9.5% |
| | **High** | 11.1% | 4.5% | 8.2% | 5.8% | 11.8% | 8.3% |
| | **No Clean** | 17.4% | 6.9% | 11.0% | 8.4% | 18.4% | 12.4% |
| **BSS** | **CCA (all)** | 12.0% | 4.8% | 8.5% | 6.2% | 12.9% | 8.9% |
| | **ICA (all, no Iclabel)** | 12.1% | 4.8% | 8.9% | 6.3% | 13.0% | 9.0% |
| | **CCA (2-stage)** | 14.5% | 5.8% | 10.1% | 7.4% | 15.0% | 10.6% |
| | **ICA (2-stage)** | 14.6% | 6.0% | 10.2% | 7.5% | 15.1% | 10.7% |

component rejection methods, auto-correlation thresholds resulted in a higher difference between RMSE for speech periods over RMSE for non-speech periods than either spectral thresholds or ICLabel. While the difference metric is negatively correlated with model performance, a higher RMSE difference implied more cleaning relative to the speech condition, which was an expectation for an effective EMG cleaning method as EMG contamination is greater during speech periods.

In the spectral RMSE and relative power analysis, lower distortion, as measured by spectral RMSE, was observed in the Delta and Theta bands and these metrics were found to be highly negatively correlated with performance. In terms of relative power changes, the changes experienced in the Theta and Delta bands for speech and non-speech periods were positively correlated with performance. For component rejection methods, spectral thresholds had higher relative changes in the Delta band, whereas the use of auto-correlation had higher relative changes in the Theta bands.

The analysis of the reconstructed artifact revealed a moderate negative correlation between the average and maximum power of the artifact. In this context, the use of CCA led to lower values for both metrics, providing support that the use of CCA may lead to higher performing decoding models.

Regarding source analysis, it was found that medium thresholds generally, two-stage approaches, and the use of CCA retained the largest number of components in the Broca area. Additionally, there was high confidence for at least half of all participants that the metric for Broca area was positively correlated with model performance, with all participants finding confidence values under 0.05 for the inclusion of clusters in Broca area for the "Vowels vs. Consonants" decoding task. Table 20 presents the summary of these metrics.

Table 20: Summary of emphasized metrics for the cleaning method selection.

| | | **RMSE Signal** | | |
|---|---|---|---|---|
| | | **S** | **NS** | **Diff.** |
| **Correlation Direction** | | **Neg.** | **Neg.** | **Neg.** |
| **Stages** | **Single Stage** | 6.35 | 5.02 | 1.34 |
| | **Two Stage** | 5.96 | 4.61 | 1.35 |
| **BSS** | **CCA** | 5.99 | 4.70 | 1.29 |
| | **ICA** | 6.38 | 4.94 | 1.44 |
| **Comp. Rej.** | **ICLabel** | 2.91 | 2.42 | 0.49 |
| | **PSD** | 5.98 | 4.83 | 1.14 |
| | **Auto-correlation** | 6.34 | 4.84 | 1.50 |

| | | **RMSE Spectral** | | | |
|---|---|---|---|---|---|
| | | **Delta** | | **Theta** | |
| | | **S** | **NS** | **S** | **NS** |
| **Correlation Direction** | | **Neg.** | **Neg.** | **Neg.** | **Neg.** |
| **Stages** | **Single Stage** | 5.29 | 3.08 | 5.94 | 1.02 |
| | **Two Stage** | 1.05 | 0.59 | 0.98 | 0.38 |

| | | **Relative Power Differences** | | | |
|---|---|---|---|---|---|
| | | **Delta** | | **Theta** | |
| | | **S** | **NS** | **S** | **NS** |
| **Correlation Direction** | | **Pos.** | **Pos.** | **Pos.** | **Pos.** |
| **Stages** | **Single Stage** | 0.57 | 0.52 | 0.46 | 0.36 |
| | **Two Stage** | 0.85 | 0.72 | 0.87 | 0.72 |
| **Comp. Rej.** | **ICLabel** | 0.19 | 0.18 | 0.21 | 0.20 |
| | **PSD** | 0.70 | 0.62 | 0.62 | 0.51 |
| | **Auto-correlation** | 0.69 | 0.60 | 0.66 | 0.52 |

| | | **Artifact Reconstruction** | |
|---|---|---|---|
| | | **Average Power** | **Highest Power** |
| **Correlation Direction** | | **Neg.** | **Neg.** |
| **BSS** | **CCA** | 29.15 | 211.99 |
| | **ICA** | 32.20 | 230.81 |

| | | **Source Analysis** | |
|---|---|---|---|
| | | **Wernicke** | **Broca** |
| **Correlation Direction** | | **Pos.** | **Pos.** |
| **Stages** | **Single Stage** | 0.64 | 1.33 |
| | **Two Stage** | 0.94 | 2.21 |
| **Thresh. Level** | **Low** | 1.25 | 1.63 |
| | **Medium** | 0.86 | 1.90 |
| | **High** | 0.40 | 1.74 |
| **BSS** | **CCA** | 0.32 | 1.72 |
| | **ICA** | 1.02 | 1.24 |

When evaluating the model performance, the collected performances pointed to a

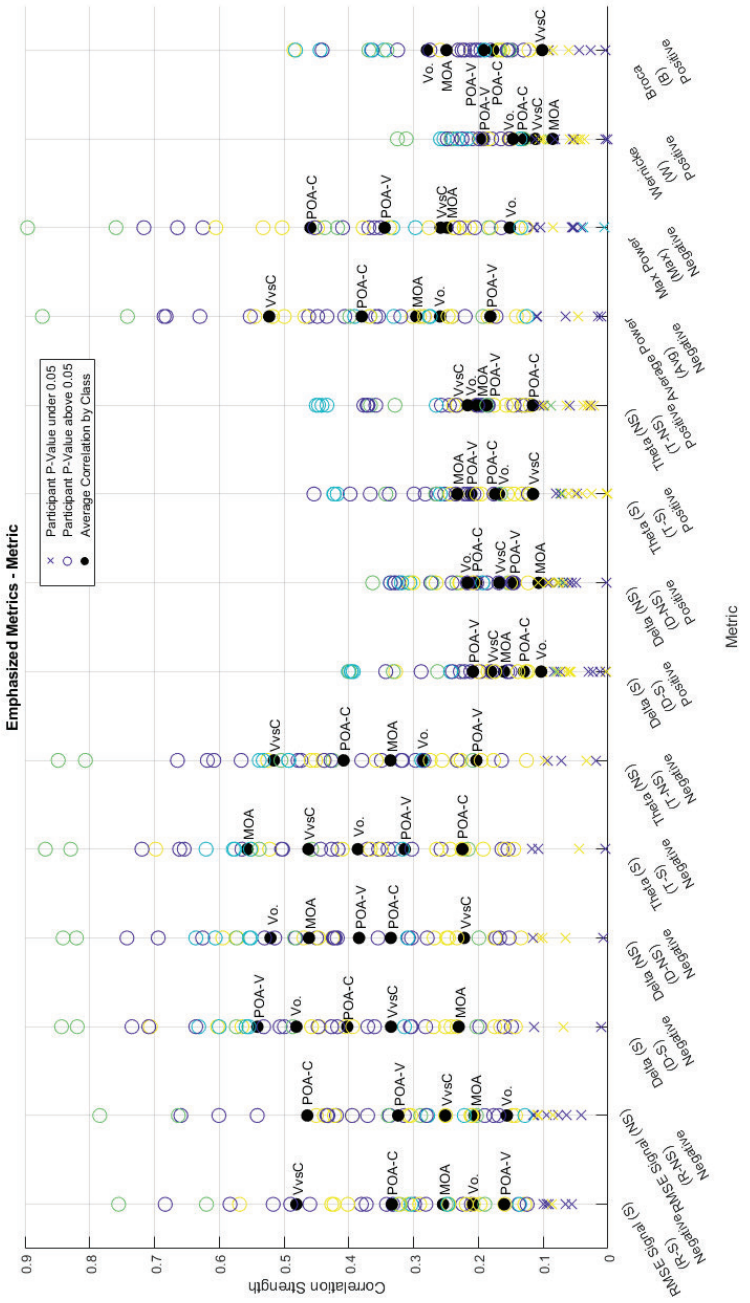


Figure 44: Emphasized metrics for the cleaning method selection.

selection of ICA-ICLabel, the pre-processed data, and a two-stage approach with ICA and low power spectral density (PSD) correlation threshold as method characteristics more closely associated with higher decoding performance.

Based on these findings, it is suggested that the datasets with the highest likelihood of more completely removing EMG contamination were based on either EEMD-ICA or EEMD-CCA. Datasets that have the greatest likelihood of obtaining high performance for speech-related decoding tasks were EEMD-CCA with medium thresholds, EEMD-ICA with low PSD thresholds, ICLabel with high thresholds (low cleaning), and the pre-processed signal itself.
